# Comprehensive insight into the alterations in the gut microbiome and the intestinal barrier as a consequence of iron deficiency anaemia

**DOI:** 10.1101/2023.04.10.536197

**Authors:** Ana Soriano-Lerma, María García-Burgos, Wiley Barton, María José Muñoz-Alférez, Jorge Valentín Crespo-Pérez, Miguel Soriano, Inmaculada López-Aliaga, Paul D. Cotter, José Antonio García-Salcedo

**Author notes:** Correspondence and requests for materials should be addressed to M.S. or I.L.A; (M.S.) (I.L.A).

## Abstract

**Background:** Iron is an essential micronutrient for all living organisms, and as such, iron deficiency is the top leading cause of anaemia. Iron supplements have been shown to be detrimental to the gut microbiome and the intestinal epithelium, triggering dysbiosis and an impaired gut barrier. However, a comprehensive analysis of these two aspects have not been performed during IDA. This study aims to delve further into the analysis of the gut microbiome in an animal model of IDA and to relate microbial changes to the biological processes occurring in the colonic epithelium, with a special focus on the gut barrier. This in-depth analysis might mean a step forward minimising the negative impact of iron supplements on intestinal health during IDA.

**Methods:** IDA was experimentally induced in an animal model through the use of an iron deficient diet. Shotgun sequencing was used to gain insight into alterations of the gut microbiome in the most affected intestinal region during IDA, the colon. Histological analyses, mRNA sequencing (RNA-Seq), qPCR and immunofluorescence were used to study transcriptionally deregulated processes in the colonic epithelium. Determinations of lipopolysaccharide and bacteria-specific immunoglobulins were performed to assess microbial translocation.

**Results:** Microbial metabolism in the colon shifted towards an increased production of certain amino acids, short chain fatty acids and nucleotides, with *Clostridium* species being enriched during IDA. Structural alterations of the colonic epithelium were shown by histological analysis. RNA-Seq revealed a downregulation of extracellular matrix-associated genes and proteins and an overall underdeveloped epithelium. Increased levels of serum LPS in the anaemic animals and an increased immune response against IDA dysbiotic bacteria support an impairment in the integrity of the gut barrier.

**Conclusions:** IDA negatively impacts the gut microbiome and the intestinal barrier, triggering an increased microbial translocation. This study emphasizes the deterioration of gut health during IDA and the fact that it should be addressed when treating the disease.

## 1. Background

Anaemia is a public health concern that affects a quarter of the total population, including 47% of children under 5 years of age and 30% of pregnant women. Despite being globally distributed, anaemia is overrepresented in low-income countries[1], imposing a considerable economic burden.

Iron deficiency is the top leading cause of anaemia[2] and the most common micronutrient deficiency worldwide [3]. Although growing children, pregnant and adult women, and the elderly have been classically considered at risk populations, there is increasing evidence to suggest that iron deficiency anaemia (IDA) can often affect patients suffering a wide variety of disorders, such as inflammatory bowel disease or *Helicobacter pylori* infection [4, 5].

The gut microbiome is gaining traction as a target to consider in the pathogenesis of IDA due to its capacity to modulate iron metabolism. Specific microbial metabolites have been shown to impair iron absorption both *in vitro* and *in vivo* through the degradation of hypoxia inducible factor 2 _α_ subunit [6]; others, however, promote iron absorption, such as p-hydroxyphenyllactic acid [7].

Iron supplements have been shown to be detrimental to the gut microbiome and the intestinal barrier [8], although studies analysing these aspects during IDA are scarce, especially in relation to the gut barrier. There is increasing evidence to suggest that a state of gut dysbiosis exists in the large intestine of patients suffering IDA, with changes in short chain fatty acid (SCFA) - producing genera [9, 10], a depletion in the phylum Proteobacteria [11], and a loss of members of the family Ruminococcaceae, order Clostridiales, class Clostridia and the genus *Faecalibacterium* [12]. Altered relative abundances of streptococci were also reported for iron deficient pregnant women[13]. Studies in animals also reveal a gut dysbiosis in the large intestine during IDA [14–16].

The gut microbiome is one of the key components of the intestinal barrier [17]. A wide variety of antigens, toxins and microorganisms are normally found in the intestinal lumen, with the intestinal barrier preventing their leakage into the bloodstream and the consequent disruption of tissue homeostasis. Considerable evidence supports the fact that the loss of integrity in the intestinal barrier along with microbial translocation to extraintestinal sites are involved in the pathogenesis or aggravation of chronic, non-infectious diseases[18]. Iron shortage impairs cell cycle progression and DNA replication[2]. In this sense, intestinal epithelial cells, the mainstay of the intestinal barrier, are characterized by a high turnover rate [17] and are likely to be affected during IDA. Hypoxia inducible factor 1 _α_ subunit (HIF1_α_) is also a key regulator of the intestinal barrier[19]. HIF1_α_ levels are maintained via mitochondrial iron-dependent mechanisms involving the bacterial production of SCFAs in the large intestine[20].

There are few studies that investigate the effect of iron deficiency on the intestinal barrier functionality [21], and none of them provide a comprehensive analysis. Most studies are focused on the implications of iron overload [21, 22]. Similarly, existing studies in animal models analysing how the gut microbiome responds to an experimentally induced IDA use 16S rRNA sequencing, which only allows a taxonomic characterisation at the genus level [14–16].

This study aims to address these current gaps in our knowledge providing an in-depth functional and structural characterisation of the colonic microbiome in response to IDA through shotgun sequencing. The intestinal barrier and the extent of microbial translocation will also be analysed and considered within the context of the gut microbiome. Analysing these hallmarks of intestinal health might be important in the clinical management of IDA to take gut protective approaches during the treatment period of the disease, contributing to its recovery.

## 2. Material and Methods

### 2.1 Animal model

Animal housing, care, handling procedures, and experimental protocols were approved by the Ethics Committee of the University of Granada and the local government Junta de Andalucía (ref 06/06/2019/100) in accordance with European guidelines (Declaration of Helsinki; Directive 2010/63/EU). Animal experiments were performed in the Animal Service of the University of Granada, with controlled sanitary and environmental parameters. Twenty weaned male Wistar rats, purchased from Charles River Laboratories (France), were used for the study, with diets and deionized water available *ad libitum*. Animals were housed in groups, using ventilated, thermoregulated cages with controlled temperature (23 ± 2 °C), humidity (60 ± 5%), and a twelve-hour circadian rhythm.

IDA was experimentally induced through an iron deficient diet for a period of 40 days, as previously described [23]. Animals were randomly divided into control (n=11) or anaemic (n=9) group, receiving the first one the AIN-93G diet[24] and the last one, the iron deficient counterpart. At the end of the experimental period, blood samples were collected from the caudal vein, using EDTA as anticoagulant to control haematological parameters. Animals were then intraperitoneally anesthetized using sodium pentobarbital (Richter Pharma AG, Austria) and bled out by cardiac puncture. Serum samples were obtained from total clotted blood via centrifugation (3000g, 10 min, 4°C). Colon segments were isolated, contents collected and immediately frozen at −80°C until analysis. Colonic mucous samples underwent fixation for immunostaining and histological analysis or snap freezing in liquid nitrogen for gene expression analysis.

### 2.2 Haematological tests

Red blood cell count, haemoglobin concentration, haematocrit, mean corpuscular volume, mean corpuscular haemoglobin, mean corpuscular haemoglobin concentration, leukocyte and platelet count were measured using an automated haematology analyser Mythic 22CT (C2 Diagnostics, France).

### 2.3 Determination of iron metabolism parameters

Serum iron, total iron binding capacity (TIBC) and ferritin were determined as biomarkers of iron deficiency.

Serum iron and TIBC were determined using the Total Iron-Binding Capacity and Serum Iron Assay Kit (MAK394, Sigma Aldrich, USA), according to the manufacturer’s instructions. Ferritin was determined in serum samples via ELISA, using Rat Ferritin ELISA Kit (ab157732, Abcam, UK).

### 2.4 Shotgun sequencing and bioinformatic analysis

DNA was isolated as described by Soriano-Lerma et al., (2022)[14]. Libraries were prepared using the DNA prep protocol (1000000025416, Illumina, USA) with 25-500ng of input DNA, according to manufactureŕs instructions. All samples were quantified using Qubit system 2.0 (Thermo Fisher Scientific, USA) and pooled according to the lowest concentration. Final pool was sequenced using the 2×151 bp P1 reagent (20050264, Illumina, USA) and NextSeq2000 sequencer, obtaining a total of 150M raw reads.

Quality processing of fastq files was performed using KneadData pipeline, based on Trimmomatic for quality trimming and Bowtie2 to remove host contaminating sequences belonging to human and rat genomes. Resulting fastq files were converted to fasta using IDBA-UD and processed using HUMAnN 3.0 pipeline[25]. Taxonomy at the species level was assigned via alignment against ChocoPhlAn v30 database and MetaPhlAn classifier. Translated search and functional profiling was performed using DIAMOND aligner and UNIPROT database, clustering sequences at 90% of homology to ensure good representation of gene families in the UniRef clusters; gene families were then regrouped using the Kyoto Encyclopedia of Genes and Genomes (KEGG) orthologs database.

Statistically significant KEGG orthologs (KO) were selected using “edgeR” package and p-adjusted values below 0.05. Selected KOs were then mapped to KEGG pathways, and pathways of interest were coloured according to log2 fold change (log_2_FC) values using KEGG Search & Colour pathway tool.

Statistically significant species were determined using ALDEx2 package and p values below 0.05.

### 2.5 Quantification of bacterial load

To quantify the total bacterial load, 16S rRNA gene-targeted quantitative PCR (qPCR) was performed. Power SYBR green PCR (4309155, Thermo Fisher Scientific, USA) was used in a total reaction mixture volume of 10 µL. The universal bacterial primers were F: 5’-AAACTCAAAKGAATTGACGGGG-3’ and R: 5’-GGGTTGCGCTCGTTRYGG-3’ [26]. Primers in a final concentration of 500 nM each and DNA volume of 1 µl (<50ng) were added to the PCR master mix in MicroAmp Fast 96-Well reaction plates (4346907, Thermo Fisher Scientific, USA). qPCRs were performed using QuantStudio 6 system (Thermo Fisher Scientific, USA) and cycling conditions included 95°C for 10 min and 40 cycles consisting of denaturalization at 95°C for 15 seconds and annealing-extension at 60°C for 1 min. Negative controls containing no template DNA were subjected to the same procedures. The specificity of the amplified products was determined by analysis of melting curves. The number of 16S copies per sample were obtained via interpolation in the standard curve, for which known concentrations of *Escherichia coli* 16S gene were used.

### 2.6 SCFAs determination

200mg of colonic contents were weighed and homogenised in 1.8mL of saline solution. Suspensions were centrifuged and filtered (0.22µm) to eliminate suspended particles. Supernatants were transferred to a vial for high performance liquid chromatography (HPLC) analysis using the Acquity UPLC-I Class System (Waters Corporation, USA) with an UV–vis detector set at 210 nm (TUV Detector). Dilutions of SCFAs standards (Acetic acid:A6283, Sigma-Aldrich; Propionic acid: 81,910, Sigma-Aldrich; Butyric acid:108,111,000, Acros Organics) were prepared in saline solution at concentrations ranging from 87 to 0.087 mM for acetic acid, 67–0.067 mM for propionic acid and 54.5–0.0545 mM in the case of butyric acid. A Waters CORTECS™ C18 column (2.1 × 100 mm, 1.6 _μ_m) was used at room temperature, at a flow rate of 0.2 mL/min; water buffer (solvent A)/acetonitrile (solvent B) gradient elution was performed as follows: from 1 to 100% B and down to 1% B, 0–7.5 min. The injected sample volume was 10uL.

### 2.7 Histological analysis

Histological analysis was performed as previously described [20], with colon fragments being collected and fixed 8 h (RT) in 4% paraformaldehyde (P6148, Sigma Aldrich, USA). Tissue dehydration, paraffin embedding, sectioning and haematoxylin&eosin staining were performed by Atrys Health S.A (Granada, Spain). Images were obtained using the Olympus BX43 microscope and analyzed blindly, calculating histological scores for each parameter described by Fachi et al., (2019) [20].

### 2.8 RNA isolation and qPCR

Total cellular RNA was isolated from colonic mucous samples with Trizol Reagent (15596, Invitrogen). Reverse transcription was performed using RevertAid First Strand cDNA Synthesis Kit (K1622, Thermo Fisher Scientific, USA) with Oligo(dT) primers according to the manufacturer’s protocol. Quantitative PCR was conducted on QuantStudio 6 (Thermo Fisher Scientific, USA) with SYBR Green (4309155, Thermo Fisher Scientific), a final concentration of primers of 500 nM and using 2µL of previously diluted cDNA (1:10). Target mRNA levels were normalized in relation to basic transcription factor 3 (BTF3) or anti-cyclophilin B (PPIB). Primers used for this study are listed in Supplementary Table 1.

### 2.9 mRNA sequencing (RNA-Seq)

RNA quality was assessed through the determination of the RNA Integrity Number (RIN) via 2100 Bioanalyzer Instrument (Agilent Technologies, USA). mRNA libraries were prepared using 500-1000 ng of total RNA as an input and the polyA selection protocol (TruSeq stranded mRNA Library Prep, ref. 20020595 / 20020594) at the Centre for Genomic Regulation (Spain). Libraries were then sequenced using the Hiseq2500 platform (Illumina, USA) and the paired-end 50bp format.

A total of 40 million raw reads per sample were obtained and mapped using “Rsubread”. Reads were aligned and annotated using the rat genome rn6 as a reference, downloaded from http://hgdownload.soe.ucsc.edu/downloads.html#rat. Further analyses in R software were performed using “edgeR” package. Gene set enrichment analysis (GSEA) was performed using “fgsea” package and log_2_FC as the ranking parameter. Gene Ontology (GO) was used as reference database (including Biological process, Cellular component and Molecular function categories). Differentially expressed GO terms during IDA were selected using p-adjusted values below 0.05; upregulation or downregulation of each pathway was assessed with positive and negative values of the Normalized Enrichment Score (NES), respectively. Visualization of parental terms was performed using REVIGO tool [27]; results derived from GSEA were also plotted using GOplot package[28].

### 2.10 Immunofluorescence

Colon fragments were harvested, washed and fixed 8 h (RT) in 4% paraformaldehyde (P6148, Sigma Aldrich, USA). Tissues were embedded in Tissue-Tek OCT Compound (Sakura Finetek, USA), frozen using isopentane and dry ice and stored at −80°C. Sections were cut (4 µm) using a cryostat and labelled overnight at 4°C with COL6 antibody (1:500) (MA5-32412, Thermo Fisher Scientific, USA) after blocking nonspecific binding sites with 10% bovine serum albumin (A7906, Sigma Aldrich, USA), 0.5% (v/v) Triton-X100 for 30 min. A secondary antibody conjugated with AlexaFluor-555 (A-31572, Thermo Fisher Scientific, USA) was used for detection (1:1000). Counterstaining was performed using DAPI (0.5µg/mL) (Invitrogen, USA). Sections were mounted in Vectashield medium (Vector Laboratories, USA). Images were acquired through a Confocal Zeiss LSM 710 inverted microscope. Immunofluorescence images were quantified measuring the area positive for both DAPI and antibody signal using ImageJ. To allow comparability, instrument settings were equally adjusted across samples.

### 2.11 Detection of bacteria-specific IgG, IgM and IgA

The determination of faecal bacteria-specific immunoglobulins was performed via ELISA as previously described [29]. Faecal bacteria belonging to the anaemic or control group were obtained using 200mg of faecal pellets. After homogenization in sterile PBS, filtration through a 100 _μ_m strainer, bacteria were separated from debris/rat cells by removing the pellet after centrifugation at 400g for 10 min (4°C). Faecal bacteria were washed, heat-killed at 85°C for 1 h, and resuspended in 20 mL PBS, and 100 _μ_L of this suspension was added to each well of a 96-well ELISA plate for overnight coating at 4°C. Number of bacteria in each suspension was determined by spectrophotometry at 600nm and adjusted across experimental groups. Wells were then blocked with 1% (w/v) BSA in PBS for 2 hr at room temperature. Rat sera were diluted at 1:100 and incubated overnight at 4°C for detection of IgG, IgM, and IgA. Incubation with secondary Anti-rat IgG, IgM and IgA (A110-143P, BioNova científica S.L., Spain) (1:10000) for 1.5h in darkness was followed by the addition of HRP substrate (11112422001, Roche Applied Science, Germany). Absorbance was measured using NanoQuant Infinite M200 Pro multi-plate reader (Tecan, Switzerland).

### 2.12 Lypopolysaccharide (LPS) detection

LPS determination was performed using LAL chromogenic assay (A39552, Thermo Fisher Scientific, USA), according to manufacturer’s instructions. Serum samples were diluted 1:20 and heat inactivated at 70°C for 15 min.

### 2.13 Statistical analysis

Pearson correlations between SCFAs and statistically significant species were calculated using R software. Network diagrams were plotted using Gephi 0.9.2.

Statistical analysis was performed using Prism GraphPad software, or R software in case of RNA-Seq and shotgun experiments. Statistical significance was assessed using Student’s two-tailed t test or a non-parametric alternative in case data were not normally distributed. In RNA-Seq and shotgun analysis, p-adjusted values were used to determine differentially expressed GO terms or statistically significant KO genes.

For all tests, p values below 0.05 were considered significant (unless otherwise indicated) and expressed as follows: *p < 0.05, **p < 0.01, ***p < 0.001 and ****p<0.0001.

## 3. Results

### 3.1 Determination of haematological parameters and study of iron metabolism confirmed the induction of IDA

A decrease in the number of red blood cells, haemoglobin concentration, haematocrit, mean corpuscular volume and mean corpuscular haemoglobin concentration by day 40 (d40) confirmed that IDA had been correctly induced (Table 1).

**Table 1.** Haematological parameters during the development of iron deficiency anaemia (day 40). Means and standard deviations are shown for each group and parameter. Statistical significance is expressed as follows: *p < 0.05, **p < 0.01 and ***p < 0.001

| Haematological parameter | Control (n=11)<br>Day 40 | Anaemia (n=9)<br>Day 40 |
| --- | --- | --- |
| Red blood cells (10<sup>6</sup>/μL) | 8.12±0.45 | 2.15±0.51* |
| Haemoglobin (g/dL) | 16.05±0.89 | 4.16±0.52*** |
| Haematocrit (%) | 47.89±2.75 | 21.34±9.27*** |
| Mean corpuscular volume (fL) | 59.14±1.54 | 40.22±0.65*** |
| Mean corpuscular haemoglobin (pg) | 19.82±0.67 | 20.60±6.27 |
| Mean corpuscular haemoglobin concentration(g/dL) | 33.49±0.75 | 25.95±17.11** |
| Leukocytes (10<sup>3</sup>/μl) | 13.23±2.25 | 10.51±1.48 |
| Platelets (10<sup>3</sup>/μL) | 889.64±445.55 | Over range |

A decrease in serum iron and ferritin along with an increase in TIBC also confirmed iron deficiency in the anaemic animals (Table 2)

**Table 2.** Study of iron metabolism. Means and standard deviations are shown for each group and parameter. Statistical significance is expressed as follows: *p < 0.05, **p < 0.01 and ***p < 0.001

| Parameters |  | Control (n=10)<br>Day 40 | Anaemia (n=8)<br>Day 40 |
| --- | --- | --- | --- |
| Serum iron (µg/L) |  | 1293 ± 80.12 | 613 ± 43.98*** |
| TIBC (µg/L) |  | 2740 ± 189 | 18110 ± 570*** |
| Ferritin (µg/L) |  | 81.5± 2.6 | 48.3± 2.01*** |
| 531 TIBC: | total iron binding capacity |  |  |

### 3.2 Microbial metabolism and gut microbiome structure shifted in response to IDA

Having previously provided a comprehensive view of gut microbial alterations occurring during IDA along the gastrointestinal tract[14], the colon was identified as the region showing the greatest dysbiosis and its microbiome was characterized using shotgun sequencing.

A total of 142M reads were obtained after bioinformatic processing. To analyse the microbial functional traits, differentially abundant KO genes between the control and anaemic groups were determined (Supplementary Figure 1, Supplementary Table 2); most genes showed positive values of log_2_FC indicating a higher abundance during IDA (Supplementary Figure 1). To assess whether these bacterial genes clustered specifically in pathways, differentially abundant KOs were selected and mapped to KEGG pathways (Table 3). Additional information on mapped KOs is displayed in Supplementary Table 3.

**Table 3.** Statistically significant KOs mapped to KEGG pathways; KOs with higher and lower abundance during IDA are shown. Pathways are wn in decreasing order of mapped KOs. Only pathways containing between 50-20 KOs were considered for subsequent analysis to exclude most general and the least abundant ones.

| KEGG pathways | Number of mapped KOs | KOs with higher abundance in<br>IDA | KOs with lower abundance in<br>IDA |
| --- | --- | --- | --- |
| Biosynthesis of amino acids | 47 | 30 | 17 |
| Pyruvate metabolism | 35 | 29 | 6 |
| Purine metabolism | 29 | 27 | 2 |
| Quorum sensing | 27 | 24 | 3 |
| Butanoate metabolism | 24 | 22 | 2 |
| Amino sugar and nucleotide<br>metabolism | 24 | 18 | 6 |
| Nucleotide metabolism | 23 | 23 | 0 |
| Propanoate metabolism | 23 | 21 | 2 |
| Glyoxylate and dicarboxylate<br>metabolism | 22 | 19 | 3 |
| Glucolysis / gluconeogenesis | 22 | 17 | 5 |
| Pyrimidine metabolism | 21 | 20 | 1 |
| Ribosome | 21 | 1 | 20 |
| Carbon fixation pathways in<br>prokaryotes | 21 | 15 | 6 |
| Sulfur metabolism | 21 | 17 | 4 |
| Oxidative phosphorylation | 21 | 15 | 6 |

Pathways of interest were then coloured according to log_2_FC values of differentially abundant KO genes, revealing a general increased abundance of genes related to biosynthesis of amino acids (Supplementary Figure 2), amino sugar and nucleotide metabolism (Supplementary Figure 3), glyclolysis and gluconeogenesis (Supplementary Figure 4), pyruvate metabolism (Supplementary Figure 5), butanoate metabolism (Figure 1), propanoate metabolism (Figure 2), purine metabolism (Supplementary Figure 7) and pyrimidine metabolism (Supplementary Figure 8) during IDA.

**Figure 1.**
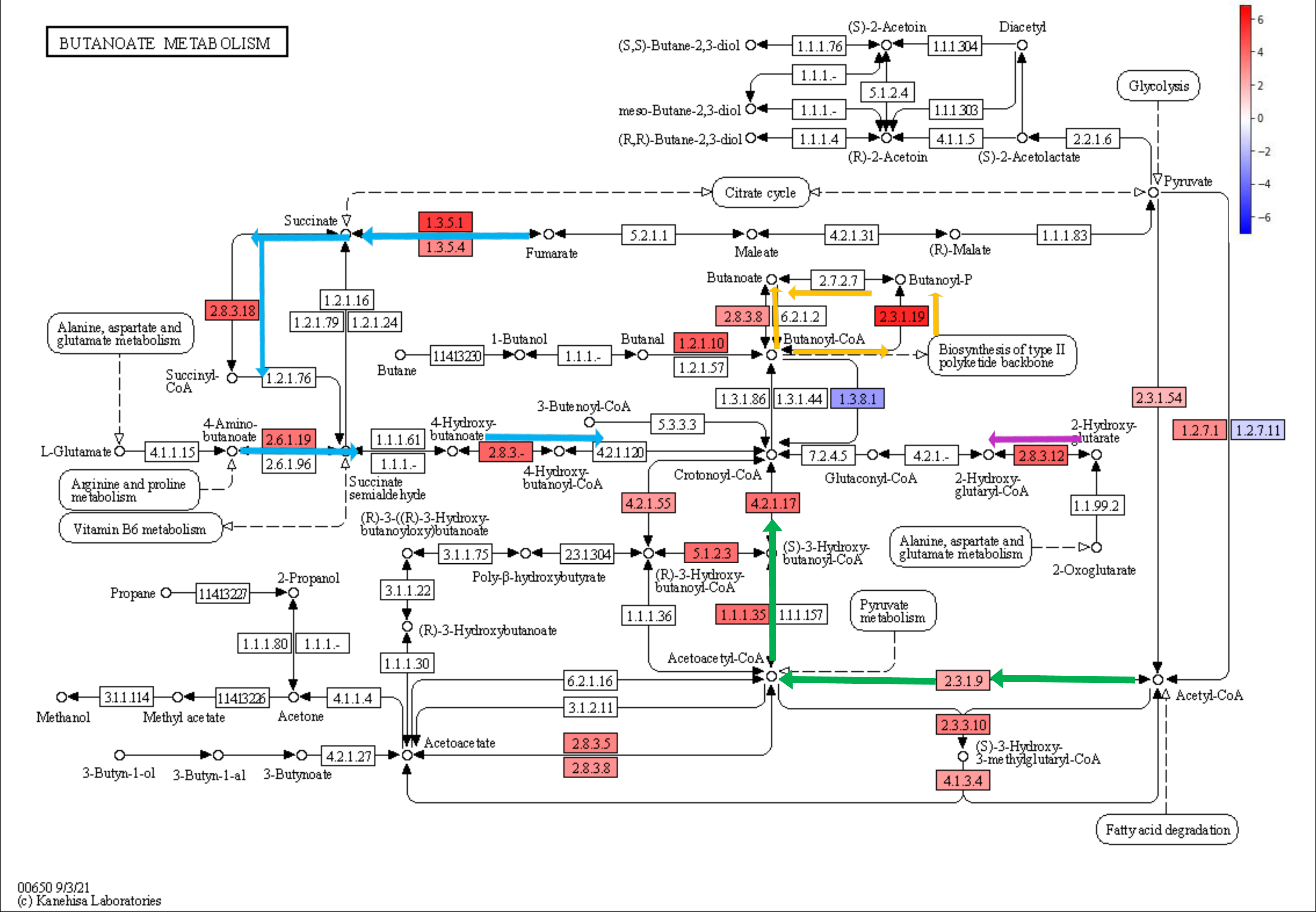
Butanoate metabolism pathway (KEGG) coloured according to log_2_FC values of statistically significant KO genes. Colour legend: 4-aminobutanoate/succinate pathway (blue), Acetyl-CoA pathway (green), glutarate pathway (purple), Butanoyl-CoA – Butanoate conversion (orange).

**Figure 2.**
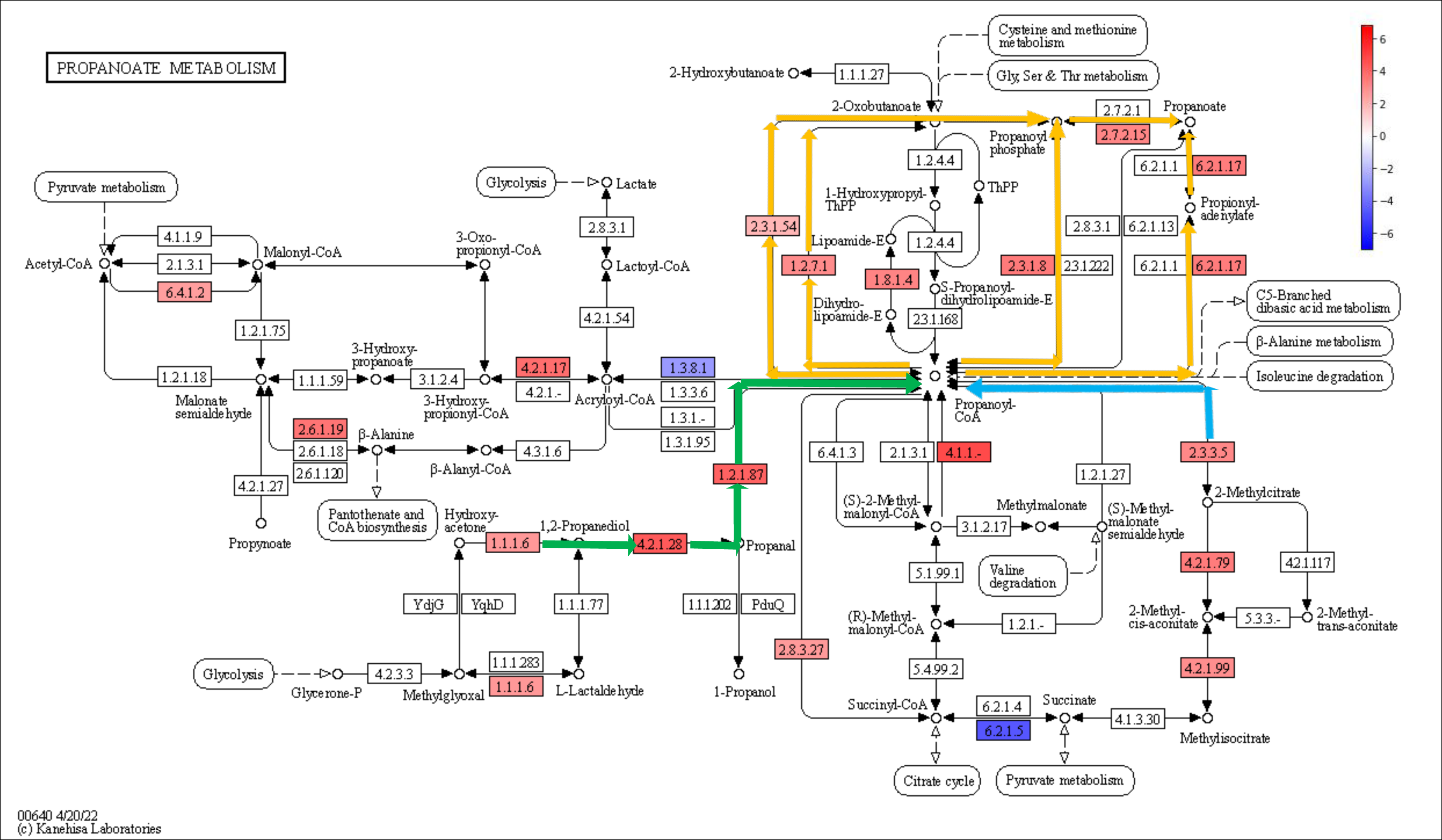
Propanoate metabolism pathway (KEGG) coloured according to log_2_FC values of statistically significant KO genes. Colour legend: 1,2-propanediol pathway (green), 2-methylcitrate pathway (blue), Propanoyl-CoA – Propanoate conversion (orange).

KO genes involved in the synthesis of certain amino acids were more abundant during IDA, such as tyrosine, phenylalanine, cysteine, methionine or alanine (Supplementary Figure 2, highlighted in green squares), while KOs involved in the synthesis of valine, leucine, or isoleucine were decreased (Supplementary Figure 2, highlighted in orange squares).

Analysis of amino sugar and nucleotide metabolism revealed a higher abundance of genes associated with peptidoglycan production (Supplementary Figure 3, highlighted in green) and with the synthesis of metabolic sugar intermediates, such as Glucose-6-P, Fructose-6-P or Mannose-6-P (Supplementary Figure 3, highlighted in blue). As far as glycolysis and gluconeogenesis is concerned, KO genes related to the transformation of monosaccharides and sugar intermediates, such as the above-mentioned ones, into pyruvate and acetyl-CoA were enriched in anaemic animals (Supplementary Figure 4, highlighted in blue). KO genes involved in the metabolism of pyruvate showed also increased abundances during IDA, leading to the formation of formate, succinate (Supplementary Figure 5, highlighted in blue) and lactate and acetate (Supplementary Figure 5, highlighted in green). Most KO genes belonging to the acetyl-CoA and 4-aminobutanoate/succinate pathway were increased within the butanoate metabolism during IDA (Figure 1, highlighted in green and blue respectively). Similarly, part of the glutarate pathway also showed higher abundance (Figure 1, highlighted in purple), as well as last steps involving the transformation of butanoyl-CoA into butanoate (Figure 1, highlighted in orange). In the case of propanoate metabolism, most pathways involved in the transformation of propanoyl-CoA into propanoate were more abundant in the anaemic group (Figure 2, highlighted in orange), along with the 1,2-propanediol pathway (Figure 2, highlighted in green) and the 2-methylcitrate pathway (Figure 2, highlighted in blue). Determination of butanoate, propanoate and acetate in the colon of control and anaemic animals confirmed an increased production of SCFAs during IDA (Supplementary Figure 6).

Lastly, purine and pyrimidine pathways were characterized by an increased abundance of genes related to the production of guanine, xanthine, hypoxanthine and adenine in the case of purine metabolism (Supplementary Figure 7, highlighted in green), and uracil, cytosine and thymine in the case of pyrimidine metabolism (Supplementary Figure 8, highlighted in green), KO genes involved in the production cyclic AMP and GMP, along with hyperphosphorylated guanine derivatives (pppGpp and ppGpp), were also more abundant during IDA (Supplementary Figure 7, highlighted in purple).

Since high synthesis of nucleotides suggest a high division rate, bacterial load was quantified by qPCR in colonic content samples belonging to anaemic and control animals. A higher number of 16S rRNA copies was found during IDA (Supplementary Figure 9).

Analysis of the differentially abundant bacterial taxa showed that *Clostridium* species, among others, were more abundant during IDA (Figure 3A). Since members of the genus *Clostridium* are among the major producers of SCFAs in the large intestine[30], Pearson correlations were calculated between SCFAs and differentially abundant species and plotted in a network diagram (Figure 3B). Node size was adjusted according to their connectivity, and so was colour using a blue-to-red scale. Correlations were plotted as edges using the same colour scale, with blue and red edges indicating negative and positive correlations, respectively. All *Clostridium* species (*Clostridium simbiosum*, *Clostridium disporicum*, *Clostridium perfringens, Clostridium innocuum* and *Clostridium bolteae*) showed positive correlations to butanoate and propanoate, and negative correlations to acetate.

**Figure 3.**
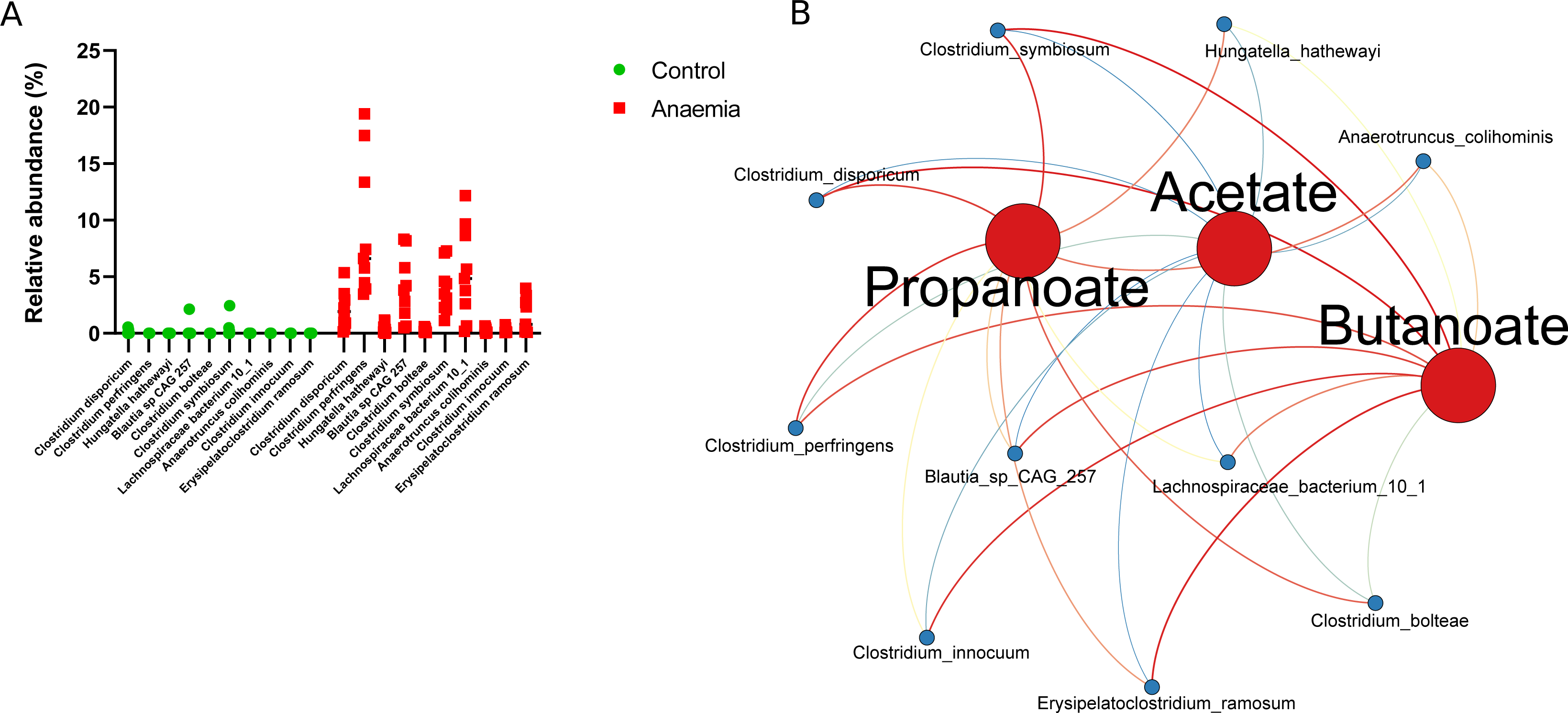
Taxonomic analysis at the species level and Pearson correlations between statistically significant species and SCFAs levels. (A) Scatter plot of differentially abundant species (p<0.05) in the control and anaemic group determined by ALDEx2 (B) Network diagram illustrating Pearson correlations between species in A and SCFAs levels. Node size and colour have been adjusted according to its connectivity in a blue-to-red scale; edges have been coloured using the same scale, whereby blue and red indicate negative and positive correlations.

### 3.3 Changes in colonic metabolism during IDA reveals hypoxia-independent alterations in the intestinal barrier

First, histological analysis was performed to provide a comprehensive view of the integrity of the colonic epithelium. Considerable epithelial damage and ulceration was found during IDA, along with leukocyte infiltration into the lamina propia and submucosa (Supplementary Figure 10A, red and black arrows respectively). A depletion of goblet cells and cells undergoing mitosis was also noticed in anaemic animals (Supplementary Figure 10A, asterisks and green circles respectively). Histological scores, indicating an altered structure of the epithelium, were calculated for both groups, with increased values in the anaemic group (Supplementary Figure 10B)

RNA-Seq was performed to gain insight into alterations in gene expression taking place during IDA in the colonic epithelium.

Gene set enrichment analysis (GSEA) was next applied to assess whether deregulated genes during anaemia clustered in specific Gene Ontology (GO) terms (Supplementary Table 4). GO database was used including all categories (Biological process, Cellular component and Molecular function). Significant parental terms resulting from GSEA were divided into upregulated (Normalized Enrichment Score, NES > 0) (Figure 4A, 4C, 4E) or downregulated pathways (NES < 0) (Figure 4B, 4D, 4F) for each category, and represented in a bubble plot. Each bubble represents a significantly deregulated parental GO term, grouped by semantic similarity. The colour of the bubbles is indicative of the NES value; the brighter the blue, the higher value of NES, which indicates that particular GO term is strongly deregulated. Only the most representative GO terms (larger sizes, see legend) were displayed to facilitate visualization.

**Figure 4.**
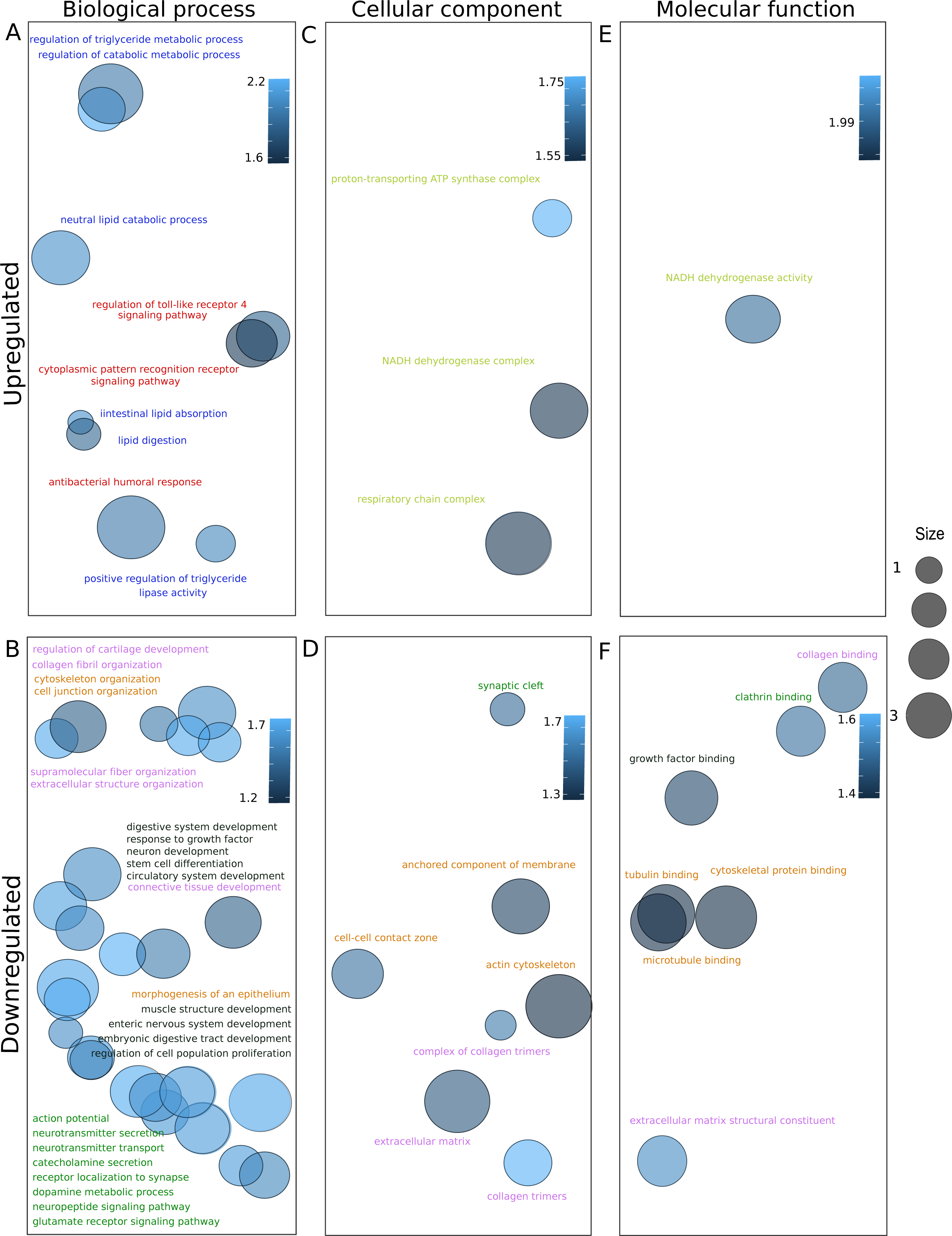
Bubble plots representing parental upregulated (Normalized Enriched Score, NES>0) or downregulated (NES<0) GO terms grouped by semantic similarity. Bubble colour have been adjusted according to NES values: the brighter the blue, the higher absolute value of NES. Bubble size is indicative of how general the GO term is in the Gene Ontology Annotation database; larger sizes indicate more general terms. Only the most general GO terms were displayed to facilitate visualization. A and B represents Biological process category, C and D Cellular component category and E and F Molecular Function category. (A) Upregulated parental GO terms during IDA (Biological process category). (B) Downregulated parental GO terms during IDA (Biological process category). (C) Upregulated parental GO terms during IDA (Cellular component category). (D) Downregulated parental GO terms during IDA (Cellular component category). (E) Upregulated parental GO terms during IDA (Molecular function category). (F) Downregulated parental GO terms during IDA (Molecular function category). Colour legend: blue (lipid metabolism); red (host-microbe interactions); yellow (mitochondrial related pathways); orange (cell junction, integrity, cytoskeletal organization); green (synaptic signalling); black (development of enteric nervous system and digestive tract); pink (extracellular matrix).

Downregulated GO terms outnumbered upregulated ones. It is notable the fact that upregulated GO terms included those involved in lipid metabolism (highlighted in blue), metabolic pathways in response to microorganisms (highlighted in red) and mitochondria-related pathways (highlighted in yellow). On the contrary, GO terms related to the development of the enteric nervous system and the digestive tract (highlighted in black) were downregulated, along with those involved in the neural signalling within the enteric nervous system (highlighted in green). Cell junction assembly and cell integrity GO terms (highlighted in orange), along with extracellular matrix-associated ones (highlighted in pink) were also downregulated.

To analyse individual gene deregulation and since the extracellular matrix is a key component of the intestinal barrier [31, 32], some GO terms related to collagen metabolism were represented in a chord diagram, along with GO terms associated with colonic intestinal dysbiosis and SCFAs (lipid) metabolism during IDA. On the left part, genes and their corresponding log_2_FC are associated to one or more selected GO terms, shown on the right (Figure 5). Downregulated genes belonged to extracellular matrix-associated pathways while upregulated ones belonged to GO terms involved in host-microbial interactions and lipid metabolism (Figure 5). Interestingly, a considerable number of genes relating to the collagen family of proteins were downregulated. Namely, *Adipocyte enhancer binding protein 1* (*AEBP1*), *collagen VI alpha 1 chain* (*COL6A1*), *fibronectin 1* (*FN1*), *lumican* (*LUM*) and *fibroblast growth factor 13* (*FGF13*) (highlighted in red) were selected along the whole spectrum of downregulation to be validated by quantitative PCR (qPCR) (Figure 6A). *COL6A1* was the most significantly downregulated gene during IDA (Figure 6A), and COL6 protein levels were assessed via immunofluorescence (Figure 6B), finding also a significant decrease (Figure 6C).

**Figure 5.**
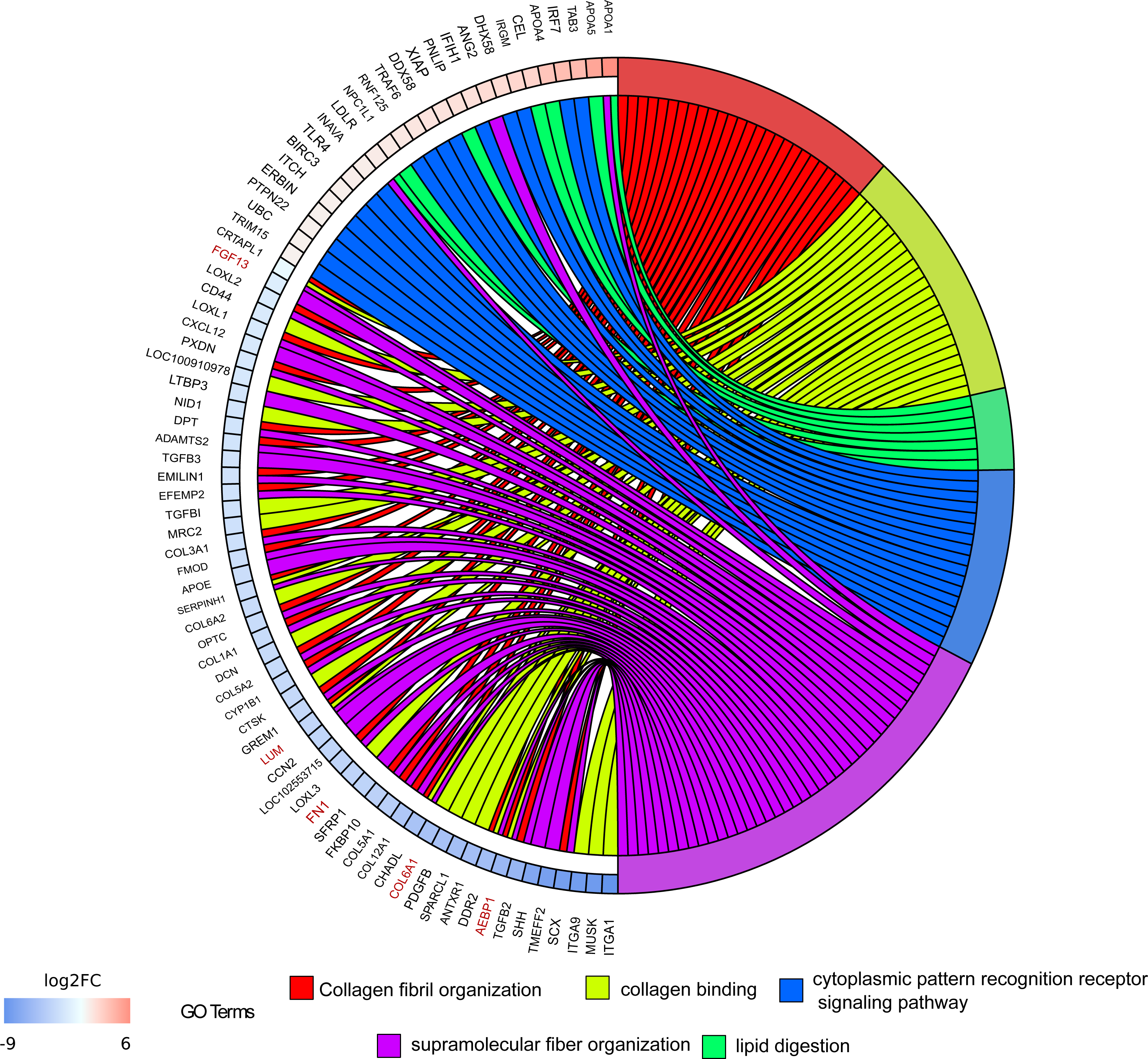
Identification of key altered genes in the colonic mucous during IDA. Chord diagram illustrating the relationships between the list of selected GO terms and their leading-edge subset of genes obtained from GSEA, including log_2_FC values. Left half of chord diagram displays whether genes are up-or downregulated during IDA; genes are linked to one or several GO terms (right half) by coloured bands, according to the legend. Highlighted in red are the selected genes for qPCR validation.

**Figure 6.**
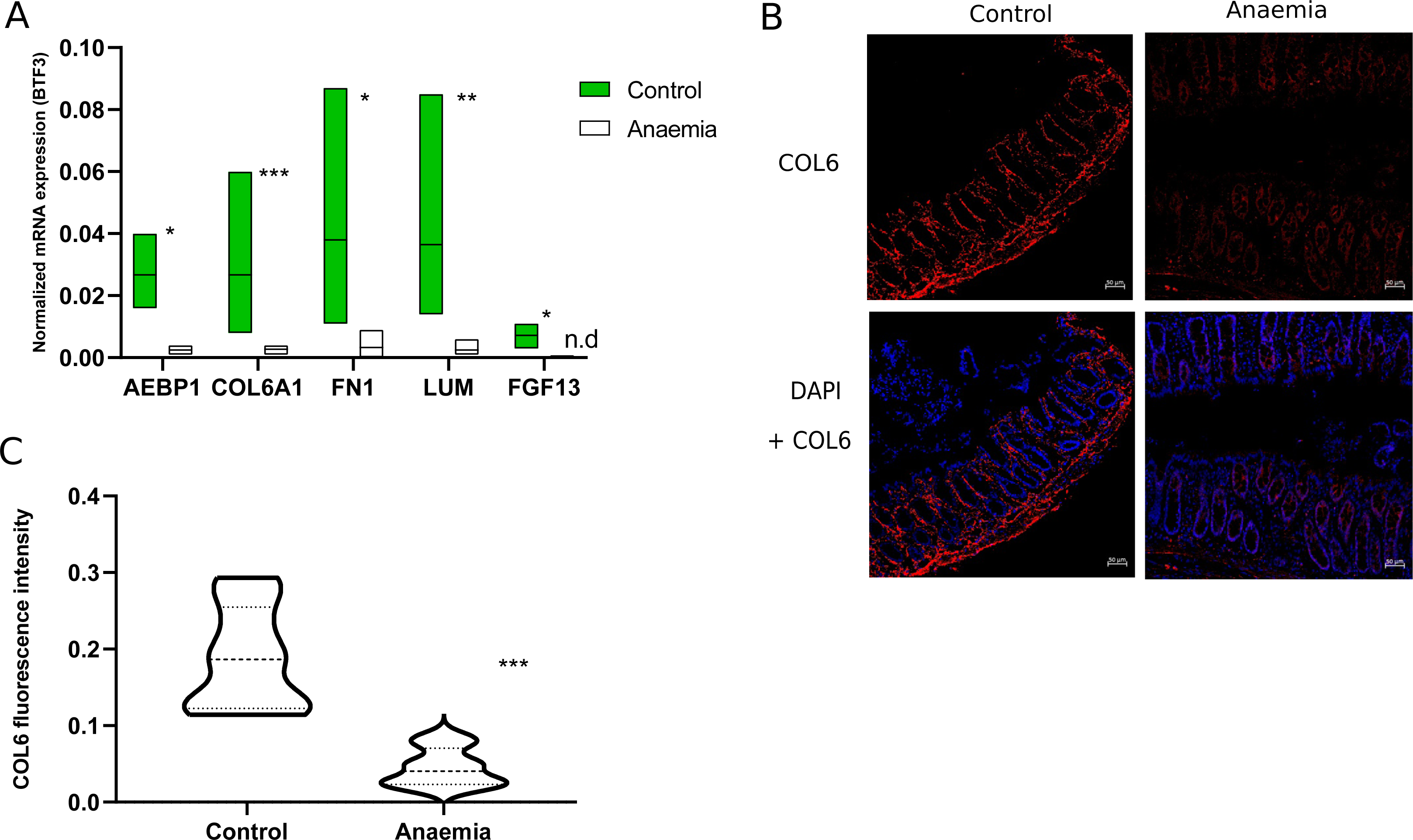
Validation of genes and proteins selected from RNA-Seq. (A) qPCR validation of previously selected genes. Normalized mRNA expression was calculated using BTF3 as housekeeping gene. *, ** or *** indicate the level of statistical significance between experimental groups. n.d.: not detected (B) Immunofluorescence showing COL6 staining (red) in colonic mucous slides belonging to anaemic and control animals; counterstaining was performed using DAPI (blue); scale bar = 50µm (C) COL6 fluorescence level quantification from sections in (B) by Image J (n = 10 images per experimental group)

Given the importance of HIF1_α_ in the maintenance of the intestinal barrier and its dependence on SCFAs levels and mitochondrial iron containing complexes, HIF1_α_ target genes were also studied by qPCR during IDA in the colonic epithelium.

Specific HIF1_α_ targets related to the intestinal barrier were selected, namely *5’-nucleotidase ecto* (*NT5E*), *ectonucleoside triphosphate diphosphohydrolase 1* (*ENTPD1*), *claudin 1* (*CLD1*), *multidrug resistance 1 gene* (*MDR1*) and *mucin 2* (*MUC2*) [19, 33, 34], and their expression measured in the colonic mucous of anaemic and control animals. Surprisingly, no significant changes were observed in any gene between experimental groups (Supplementary Figure 11).

### 3.4 Increased LPS translocation and immune response towards dysbiotic bacteria is observed during IDA as a consequence of the impaired gut barrier

LPS translocation was analysed in serum samples, along with bacteria-specific immunoglobulins, to assess the consequences of the impaired gut barrier.

LPS detection in serum samples was significantly higher for anaemic rats compared to control ones (Figure 7A). When assessing immune response, immunoglobulin detection against autologous faecal bacteria was greater for anaemic animals compared to control (Figure 7B, “Paired faeces”). However, no significant differences were found in the heterologous immune response of the control and anaemic animals against bacteria obtained from faeces belonging to the control group (Figure 7B, “Control faeces”).

**Figure 7.**
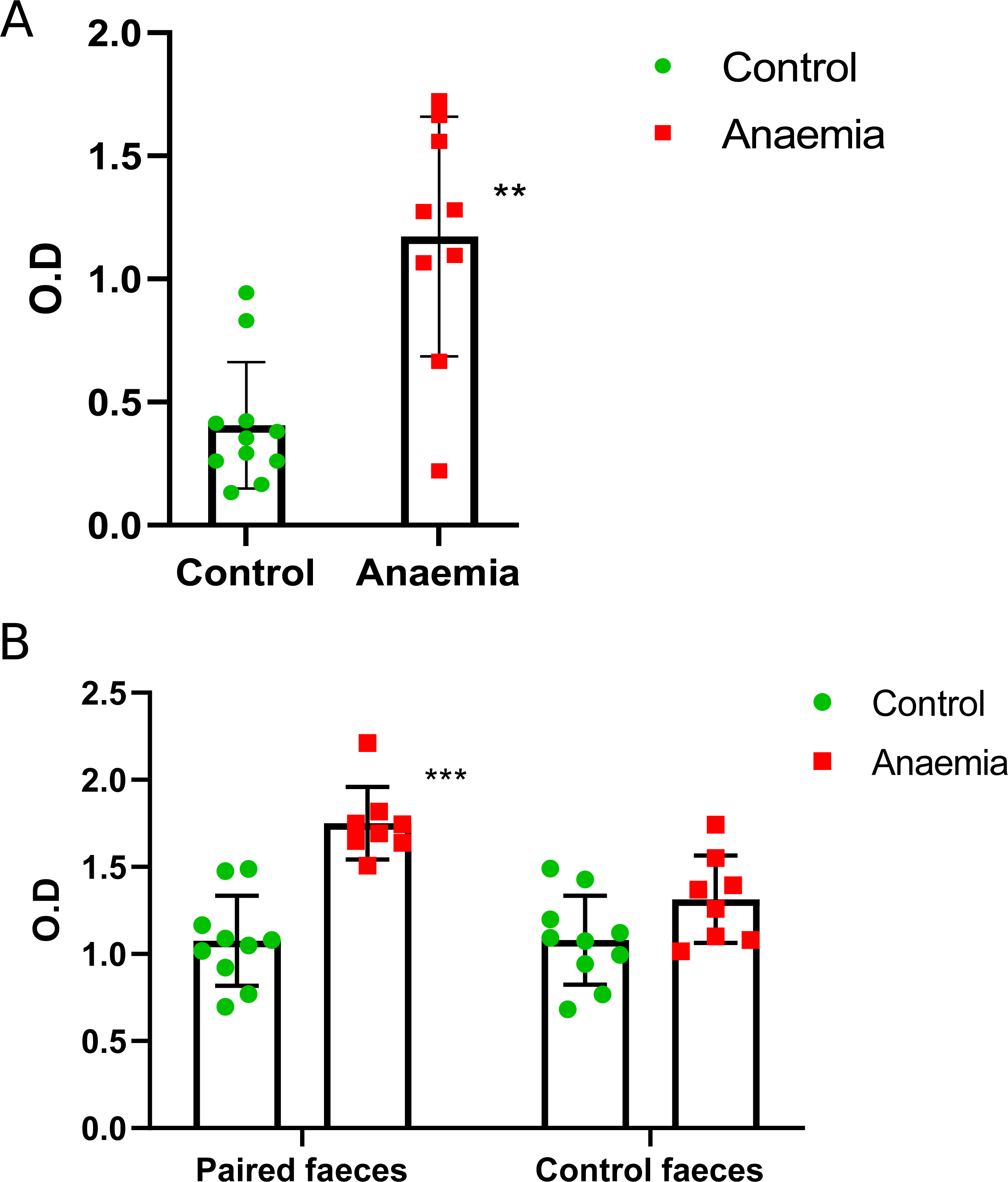
Study of microbial translocation biomarkers in serum samples belonging to anaemic and control animals. Mean and standard deviations, along with individual values for each sample (green circles for the control group and red squares for the anaemic group) are shown for biological replicates. Statistical significance is expressed as follows: *p < 0.05; **p < 0.01 and ***p < 0.001 (A) LPS levels in both experimental groups. (B) Bacteria-specific IgG, IgA and IgM in serum samples of both experimental groups. Left bar diagram shows the autologous immune response of control and anaemic animals against bacteria obtained from faecal pellets belonging to their own experimental group (*Paired faeces*). Right bar diagram represents the heterologous immune response of control and anaemic animals against bacteria obtained from faecal pellets belonging to the control group (*Control faeces*).

## 4. Discussion

Complex interactions take place between the host and the gut microbiome, especially when the state of health is compromised. This study aims to provide an in-depth insight into the gut microbiome in the most affected intestinal region during IDA through shotgun sequencing. A comprehensive analysis of the transcriptional changes occurring in the colonic epithelium was also carried out, including a structural analysis of the intestinal barrier and the study of intestinal barrier biomarkers such as extracellular matrix associated genes and proteins and HIF1_α_ target genes. Lastly, microbial translocation was assessed as an indicator of the barrier permeability.

According to Malinowska et al., (2022)[35], diet is a key influential factor on the gut microbiome functioning. During iron deficiency, metabolism of carbohydrates leaned towards an increased processing of monosaccharides through the amino sugar and nucleotide pathway (Supplementary Figure 3, highlighted in green and blue), glycolysis and gluconeogenesis (Supplementary Figure 4, highlighted in blue) and pyruvate metabolism (Supplementary Figure 5, highlighted in green and blue). End products from these pathways included acetyl-CoA, formate or succinate. In this sense, crossfeeding mechanisms might contribute to SCFAs production since the acetyl-CoA and 4-aminobutanoate/succinate pathways within butanoate metabolism were increased during IDA (Figure 1, highlighted in green and blue respectively). Similarly, certain amino acids, such as cysteine, methionine and alanine whose synthesis pathways were increased during IDA (Supplementary Figure 2, highlighted in green squares), can also be utilized in the production of SCFAs [36].

Moreover, microbial metabolism in the colon shifted towards an increased production of nucleotides and nucleotide derivatives (Supplementary Figure 7 highlighted in green and purple, Supplementary Figure 8 highlighted in green). Synthesis of hyperphosphorylated guanine derivatives have been related to cellular reprogramming occurring during stress in bacteria [37], such as iron-limiting conditions. These molecules, as well as cyclic nucleotides, can also be involved in quorum sensing signalling, regulating bacterial communication with the environment [38].

Structural alterations of the colonic epithelium were observed during IDA, including depletion of goblet cells, epithelial damage and leukocyte infiltration into the lamina propia and submucosa (Supplementary Figure 10A). mRNA isolated from colonic mucous samples was sequenced in an attempt to identify the most important changes at the transcriptional level and relate them to the gut microbiome structure and functioning and the intestinal barrier. The gut barrier is considerably affected during IDA, and our results suggest the gut microbiome might be exerting trade-off effects through the production of SCFAs. In fact, GO terms related to lipid metabolism and host-microbial interactions are upregulated in the colonic epithelium, suggesting a crosstalk occurring between the host and the microbiome (Figure 4A, highlighted in blue and red respectively, Figure 5).

GSEA revealed a general downregulation during IDA (Figure 4B, 4D, 4F), which might be indicative of an impaired gut barrier. Among others, the development of the enteric nervous system was negatively affected during IDA (Figure 4B) and some authors have proposed glial cells as modulators of barrier permeability [18]. In this sense, SCFAs might be involved in increasing enteric neural survival and neurogenesis [39] as well as in promoting natural turnover of colonocytes to make up for an underdeveloped epithelium [40]. More importantly, extracellular matrix-associated pathways and genes were also diminished during IDA (Figure 4B, 4D, 4F highlighted in pink, Figure 6 and Figure 7). Extracellular matrices are three-dimensional networks mainly composed of collagens and other macromolecules. Not only do they provide a physical scaffold for the development of tissues and organs but they also supply cells with chemicals to regulate proliferation, survival and differentiation[41]. Disruption of the extracellular matrix has been related to damage in the intestinal barrier, while its recovery has been associated with the restoration of the intestinal epithelium [31, 32]. Iron containing proteins are required for collagen metabolism [42], which supports the downregulation of collagen genes during IDA (Figure 5). In particular, *COL6A1* mRNA and COL6 protein levels were validated by qPCR and immunofluorescence, confirming a decrease in the colonic mucous of anaemic animals (Figure 6A, 6B and 6C).

Despite the increase in SCFAs during IDA (Supplementary Figure 6) and the upregulation of GO terms related to the electron transport chain, no differences in HIF1_α_ targets were found between the anaemic and control groups (Supplementary Figure 11). HIF1_α_ stabilization by SCFAs is mediated by iron-containing proteins in the mitochondria[43], and iron deficiency might therefore impair mitochondria-dependent hypoxic responses. Altogether, these results suggest that the observed damage in the intestinal barrier is independent of hypoxia levels. Lastly, LPS and bacteria-recognizing Ig were measured in serum samples to assess whether the impaired barrier functionality was responsible for an increased translocation during IDA. LPS levels were increased in serum samples belonging to anaemic animals compared to control ones (Figure 7A). Immunoglobulin detection of autologous faecal bacteria was also higher for the anaemic group, while no differences between anaemic and control animals were found when assessing immune response towards bacteria obtained from faecal control samples (Figure 7B). These results suggest an increased immune response is occurring against dysbiotic gut bacteria during IDA. Even though SCFAs production seems to play a beneficial role in this context, intestinal dysbiosis is known to trigger inflammatory and immune responses[18].

## 5. Conclusions

The results presented in this study show that IDA has a considerable impact on the gut microbiome, colonic metabolism and the intestinal barrier. An alteration in the structure of colonic microbial communities occurred in response to IDA, with a predominance of *Clostridium* species, high production of SCFAs and increased bacterial load. An overall deteriorated state of the colonic epithelium was also found, with structural components of the gut barrier being affected by iron deficiency, such as extracellular matrix-associated genes and proteins. As a result, increased levels of LPS were found in serum samples of anaemic animals along with an increased immune response towards gut dysbiotic bacteria. Further investigation is still needed with regards to the effects of SCFAs and the implication of hypoxia as a modulator of the gut barrier functionality during IDA. The existing intestinal dysbiosis and the intestinal barrier functionality should be restored during the treatment period of IDA and might even lead to a more efficient recovery of the disease.

## Supplementary materials

Supplementary materials from this study will be included separately from the main manuscript.

## Acknowledgements

The authors would like to thank Dr Fiona Crispie, Teagasc Agriculture and Food Development Authority (Ireland) and the Genomics Unit at the Center for Genomic Regulation (Spain) for their assistance in the sequencing service. The authors are grateful to Atrys Health S.A (Granada, Spain) for their assistance in the histological analysis.

## Author contributions

M.S, I.L.A and J.A.G.S developed the original idea and provided financial support. A.S.L, M.G.B and M.J.M.A carried out the in vivo study and contributed to the production of experimental data. P.C supervised shotgun sequencing; A.S.L. and W.B. performed bioinformatic analysis. J.V.C.P. performed the histological analysis. A.S.L wrote the original draft; M.S., I.L.A, P.C and J.A.G.S edited the final version of the manuscript.

## Data availability statement

All datasets supporting the conclusions of this article will be made available in the Sequence Read Archive (SRA) of the National Centre for Biotechnology Information (NCBI) upon publication. Authors can confirm that all relevant data are included in the article and/or its supplementary information files.

## Abbreviations

AEBP1: Adipocyte enhancer binding protein 1
BTF3: Basic transcription factor 3
CLD1: Claudin 1
COL6: Collagen VI
COL6A1: Collagen VI alpha 1 chain
D40: Day 40 during the induction of iron deficiency anaemia
ENTPD1: Ectonucleoside triphosphate diphosphohydrolase 1
F: Forward
FGF13: Fibroblast growth factor 13
FN1: Fibronectin 1
GO: Gene onthology
GSEA: Gene set enrichment analysis
HIF1α: Hypoxia inducible factor 1 α
HPLC: High performance liquid chromatography
IDA: Iron deficiency anaemia
Igs: Immunoglobulins
IgA: Immunoglobulin A
IgG: Immunoglobulin G
IgM: Immunoglobulin M
KEGG: Kyoto Encyclopedia of Genes and Genomes
KO: KEGG orthologs
MDR1: Multidrug resistance gene 1
MUC2: Mucin 2
NES: Normalized enrichment score
NT5E: 5’-nucleotidase ecto
Log_2_FC: Log2 fold change
LPS: Lipopolysaccharide
LUM: Lumican
PPIB: Anti-cyclophilin B
R: Reverse
SCFA: Short chain fatty acids
SOP: Standard operating procedure
TIBC: Total iron binding capacity

