## Supplementary material for "Comprehensive insight into the alterations in the gut microbiome and the intestinal barrier as a consequence of iron deficiency anaemia"

[a]Department of Physiology (Faculty of Pharmacy, Campus Universitario de Cartuja), Institute of Nutrition and Food Technology "José Mataix Verdú", University of Granada, E-18071 Granada, Spain. [b]GENYO. Centre for Genomics and Oncological Research: Pfizer / University of Granada / Andalusian Regional Government, PTS Granada, E-18016, Granada, Spain. [c]Instituto de Investigación Biosanitaria ibs.GRANADA, Granada, Spain. [d]VistaMilk, Ireland. [e]Teagasc Food Research Centre, Moorepark, Fermoy, Cork, Ireland. [f]Service of Anatomical pathology, Intercenter Regional Unit Granada, Granada, Spain. [g]Center for Intensive Mediterranean Agrosystems and Agri-food Biotechnology (CIAIMBITAL), University of Almeria, E-04001, Almería, Spain. [h]APC Microbiome Ireland, Cork, Ireland. [i]Microbiology Unit, University Hospital Virgen de las Nieves, Granada, Spain

**Supplementary Table 1.** qPCR primers used in this study. A list of all forward (F) and reverse (R) primers and their respective sequences. Cycling conditions included 95°C for 10 min and 40 cycles consisting of denaturalization at 95°C for 15 seconds and annealing-extension at 60°C for 1 min. For NT5E, ENTPD1 and CLD1 genes, the annealing-extension temperature was adjusted to 50°C to optimise performance.

| Name | Gene | Pair | Sequence (5' → 3') |
|---|---|---|---|
| BTF3 | basic transcription factor 3 | F | TGGCAGCAAACACCTTCACC |
| | | R | AGCTTCAGCCAGTCTCCTCAAAC |
| PPIB | anti-cyclophilin B | F | TCAAGCTGAAGCACTATGGG |
| | | R | GCAATGGCAAAGGGTTTCTC |
| AEBP1 | adipocyte enhancer binding protein 1 | F | GGGTCCAGTGGAGAAAATCA |
| | | R | GTTGTCCTCAATGCGGTGT |
| COL6A1 | collagen VI alpha 1 chain | F | GCGATTGCCTTCCAAGACT |
| | | R | CCTCAAGGCCACACTCTCC |
| FN1 | fibronectin 1 | F | GATGCCGATCAGAAGTTTGG |
| | | R | GGTCGTGCAGATCTCCTCAT |
| LUM | lumican | F | CCTTCAACACAACCAGCTCA |
| | | R | GTGACTTAAGGCCTTTCAGAGAA |
| FGF13 | fibroblast growth factor 13 | F | AGGCAGATGGAACCATTGAT |
| | | R | CCCACAGGGATGAGGTTAAA |
| NT5E | 5'-nucleotidase ecto | F | CTTCTGAACAGCACCATTCG |
| | | R | CCTCCACTGGTTAATGTCTGC |
| ENTPD1 | ectonucleoside triphosphate diphosphohydrolase 1 | F | GTCGTCTCACACCAACCTGT |
| | | R | GGTGTCTGGTGCTGTTTGGA |
| CLD1 | claudin 1 | F | TGTGTCCACCATTGGCATGA |
| | | R | ACAGGGGTCATGGGGTCATA |
| MUC2 | Mucin 2 | F | GCCAGATCCCGAAACCA |
| | | R | TATAGGAGTCTCGGCAGTCA |
| MDR1 | multidrug resistance 1 gene | F | GCTTCTTCCAAAGTGTACATCTTG |
| | | R | TGCCCATCTTTGAGAAGTTCTTG |

**Supplementary Table 2.** Statistically significant KOs between the control and anaemic group (padj < 0.05). $Log_2FC$ values were calculated using the control group as a reference.

| KO number | log2FC | padj |
|---|---|---|
| K00005 | 2.67712 | 0.01349049 |
| K00009 | 4.44472 | 0.00586344 |
| K00010 | 5.79709 | 0.00293278 |
| K00030 | -3.3807 | 0.01391732 |
| K00042 | 5.48833 | 0.00232026 |
| K00053 | -1.9317 | 0.00926502 |
| K00060 | -3.3695 | 0.00656207 |
| K00073 | 3.274 | 0.01840139 |
| K00075 | 2.07828 | 0.00905887 |
| K00077 | 3.99261 | 0.00379344 |
| K00087 | 4.20071 | 0.00634498 |
| K00094 | 3.27801 | 0.01583079 |
| K00098 | 3.21621 | 0.02066331 |
| K00101 | 3.69023 | 0.01685147 |
| K00104 | 2.07979 | 0.03595368 |
| K00108 | 4.22927 | 0.00581135 |
| K00111 | 2.51573 | 0.01573715 |
| K00113 | 3.39666 | 0.02153984 |
| K00116 | 3.83559 | 0.00778852 |
| K00117 | 2.7305 | 0.04624617 |
| K00121 | 3.98637 | 0.01132152 |
| K00123 | 3.70825 | 0.00542926 |
| K00124 | 3.56133 | 0.00670795 |
| K00127 | 2.99892 | 0.01460036 |
| K00131 | 5.72998 | 0.00232026 |
| K00138 | 3.44351 | 0.01344649 |
| K00146 | 2.66321 | 0.03739153 |
| K00161 | -3.0295 | 0.03083665 |
| K00164 | 2.67376 | 0.03916421 |
| K00170 | 3.0141 | 0.01840139 |
| K00175 | -1.3124 | 0.03806254 |
| K00180 | -3.0101 | 0.01746165 |
| K00198 | -2.3292 | 0.0365319 |
| K00208 | 2.93512 | 0.01970772 |
| K00219 | 3.21495 | 0.0158158 |
| K00230 | 4.91251 | 0.00466824 |
| K00242 | 5.15947 | 0.00293278 |
| K00244 | 3.04272 | 0.0173509 |
| K00245 | 4.57035 | 0.00466824 |
| K00247 | 4.35268 | 0.00796851 |
| K00248 | -2.8156 | 0.03113942 |
| K00254 | -3.8568 | 0.00615134 |
| K00259 | 3.15414 | 0.04649492 |

| KO number | log2FC | padj |
|---|---|---|
| K05846 | 3.80291 | 0.00751 |
| K05873 | 5.43797 | 0.00263 |
| K05878 | 3.5541 | 0.01251 |
| K05887 | 3.27338 | 0.03941 |
| K05895 | 5.74103 | 0.00232 |
| K05896 | 2.3626 | 0.01997 |
| K05916 | 3.70086 | 0.008 |
| K05919 | 2.61106 | 0.04305 |
| K05939 | 3.4292 | 0.01764 |
| K05942 | -3.2029 | 0.02284 |
| K05966 | 3.30264 | 0.01226 |
| K05979 | 5.52063 | 0.00232 |
| K05982 | 3.46262 | 0.01332 |
| K05989 | 2.86951 | 0.02761 |
| K05997 | 2.63695 | 0.04291 |
| K06001 | -2.7 | 0.00488 |
| K06006 | 4.75529 | 0.0047 |
| K06019 | 5.73701 | 0.00272 |
| K06023 | 2.68253 | 0.0087 |
| K06024 | 2.87697 | 0.00644 |
| K06039 | 3.16334 | 0.03493 |
| K06048 | 4.59218 | 0.00775 |
| K06073 | 3.59683 | 0.02853 |
| K06074 | 2.93381 | 0.02283 |
| K06075 | 3.03147 | 0.02966 |
| K06077 | 4.39796 | 0.0064 |
| K06078 | 5.10315 | 0.00552 |
| K06080 | 3.71499 | 0.00767 |
| K06122 | 5.68572 | 0.00232 |
| K06131 | 2.73929 | 0.00897 |
| K06133 | 3.27975 | 0.02264 |
| K06140 | 4.91492 | 0.0042 |
| K06144 | 2.74386 | 0.03988 |
| K06145 | 3.29909 | 0.0188 |
| K06149 | 3.50331 | 0.00899 |
| K06155 | 2.98098 | 0.02612 |
| K06157 | 3.77697 | 0.01262 |
| K06162 | 4.14212 | 0.00799 |
| K06167 | -4.708 | 0.00586 |
| K06175 | 4.87987 | 0.00531 |
| K06176 | 3.61374 | 0.01434 |
| K06177 | 3.75598 | 0.00999 |
| K06179 | 2.12078 | 0.03036 |

| | | |
|---|---|---|
| K00261 | -3.3701 | 0.02459467 |
| K00262 | -1.6158 | 0.00686273 |
| K00265 | -1.4807 | 0.04870107 |
| K00266 | 1.97983 | 0.02355585 |
| K00275 | 4.71166 | 0.00586344 |
| K00278 | 3.70383 | 0.00761685 |
| K00281 | -2.8205 | 0.01472815 |
| K00282 | 3.89121 | 0.01186039 |
| K00285 | 3.83113 | 0.00761685 |
| K00290 | -2.4153 | 0.00854133 |
| K00294 | -5.1054 | 0.00371482 |
| K00313 | 4.55261 | 0.00586344 |
| K00324 | 3.7821 | 0.00972285 |
| K00325 | 4.38503 | 0.00609582 |
| K00331 | -2.4466 | 0.01301388 |
| K00333 | -5.9112 | 0.00334265 |
| K00334 | 2.39415 | 0.0302802 |
| K00338 | 4.73317 | 0.00466824 |
| K00340 | -3.4713 | 0.01408669 |
| K00362 | 2.91404 | 0.02604288 |
| K00370 | 2.51992 | 0.0230307 |
| K00371 | 2.82547 | 0.02177878 |
| K00372 | 5.47175 | 0.00262888 |
| K00373 | 3.16514 | 0.01130848 |
| K00374 | 2.70051 | 0.01710558 |
| K00380 | 3.48101 | 0.01933834 |
| K00381 | 3.22596 | 0.0173414 |
| K00382 | 3.20707 | 0.0095326 |
| K00385 | 6.00628 | 0.00232026 |
| K00389 | 5.44555 | 0.00330101 |
| K00394 | -4.3082 | 0.00665826 |
| K00425 | 2.67167 | 0.01163158 |
| K00427 | 4.39455 | 0.00586344 |
| K00432 | 4.95543 | 0.00530562 |
| K00526 | 2.84991 | 0.03090499 |
| K00548 | 3.4059 | 0.00805161 |
| K00558 | -2.723 | 0.00371604 |
| K00563 | 2.74058 | 0.02459467 |
| K00564 | 3.31008 | 0.01426131 |
| K00568 | 3.5143 | 0.01334336 |
| K00573 | 3.18121 | 0.01726531 |
| K00574 | 3.00524 | 0.00839813 |
| K00595 | 3.79624 | 0.00761685 |
| K00598 | 3.14672 | 0.02460164 |
| K00620 | 2.92012 | 0.01483334 |
| K00625 | 1.58718 | 0.04739611 |
| K00626 | 2.24334 | 0.01840139 |
| K00627 | -2.8798 | 0.01454205 |
| K06182 | 6.05813 | 0.00232 |
| K06186 | 3.51739 | 0.01336 |
| K06188 | 3.50086 | 0.01557 |
| K06189 | 3.36075 | 0.01737 |
| K06190 | 3.0346 | 0.02892 |
| K06191 | 3.14718 | 0.0179 |
| K06194 | 3.03583 | 0.01764 |
| K06195 | 3.49515 | 0.03065 |
| K06199 | -1.113 | 0.03601 |
| K06200 | 3.98262 | 0.00488 |
| K06202 | 4.6472 | 0.00586 |
| K06203 | 4.0745 | 0.00986 |
| K06205 | 3.25363 | 0.03823 |
| K06206 | 4.76761 | 0.00272 |
| K06209 | 2.67271 | 0.00777 |
| K06211 | 4.86581 | 0.00467 |
| K06214 | 4.06643 | 0.00865 |
| K06216 | 2.90077 | 0.03823 |
| K06222 | 3.93514 | 0.00985 |
| K06223 | -1.9909 | 0.02332 |
| K06281 | 2.8024 | 0.02077 |
| K06282 | 4.0674 | 0.0049 |
| K06283 | -1.923 | 0.04144 |
| K06297 | 3.70401 | 0.01464 |
| K06298 | -4.5885 | 0.00488 |
| K06346 | 2.50127 | 0.01409 |
| K06381 | 2.86705 | 0.00984 |
| K06382 | 3.31751 | 0.00649 |
| K06383 | 3.99543 | 0.00488 |
| K06384 | 5.75356 | 0.00232 |
| K06387 | 1.97992 | 0.02316 |
| K06390 | 2.28618 | 0.02892 |
| K06391 | 3.95244 | 0.01251 |
| K06392 | -1.6301 | 0.04812 |
| K06393 | 3.84063 | 0.00531 |
| K06394 | 2.79378 | 0.02287 |
| K06396 | 2.57181 | 0.03722 |
| K06400 | 5.35242 | 0.00265 |
| K06409 | 5.14586 | 0.00232 |
| K06410 | 2.36108 | 0.03062 |
| K06415 | 5.66527 | 0.00232 |
| K06416 | -3.1587 | 0.01262 |
| K06438 | 5.20082 | 0.00272 |
| K06518 | 2.22221 | 0.02852 |
| K06726 | 2.63858 | 0.01364 |
| K06859 | 2.73124 | 0.04302 |
| K06861 | -1.7546 | 0.01645 |
| K06866 | 3.08189 | 0.03916 |

| | | | | | |
|---|---|---|---|---|---|
| K00631 | 3.97114 | 0.00774545 | K06867 | 2.52406 | 0.04631 |
| K00632 | 3.58561 | 0.00635964 | K06878 | 3.15459 | 0.00751 |
| K00633 | -2.313 | 0.04135602 | K06879 | 4.39136 | 0.00543 |
| K00634 | 5.7631 | 0.00232026 | K06881 | 2.47063 | 0.02818 |
| K00640 | 1.8668 | 0.04065595 | K06890 | 2.50407 | 0.0218 |
| K00641 | 4.2292 | 0.00609582 | K06891 | 3.20074 | 0.01653 |
| K00645 | 2.65337 | 0.01073472 | K06896 | 2.60292 | 0.01454 |
| K00656 | 1.85073 | 0.03891044 | K06897 | 3.37716 | 0.00857 |
| K00674 | 5.47157 | 0.00232026 | K06899 | 3.06931 | 0.01922 |
| K00675 | 3.74952 | 0.01358631 | K06909 | -4.1971 | 0.0061 |
| K00677 | -1.8707 | 0.03105988 | K06910 | -2.4728 | 0.00531 |
| K00681 | 3.51187 | 0.01250024 | K06911 | 4.03066 | 0.00392 |
| K00684 | -2.3666 | 0.04302045 | K06916 | 3.61798 | 0.01341 |
| K00688 | 3.12851 | 0.00684108 | K06919 | -3.2695 | 0.00579 |
| K00700 | 1.88662 | 0.02059078 | K06921 | -3.0301 | 0.00531 |
| K00754 | 2.6843 | 0.04427196 | K06923 | 3.05636 | 0.00854 |
| K00757 | 2.95907 | 0.01963605 | K06925 | 2.10611 | 0.0274 |
| K00769 | 3.61578 | 0.01867853 | K06926 | -3.655 | 0.00353 |
| K00789 | -1.5001 | 0.00854133 | K06956 | 4.29185 | 0.00531 |
| K00795 | 4.64255 | 0.00379344 | K06957 | 2.77701 | 0.04302 |
| K00799 | 2.26581 | 0.041774 | K06958 | 2.44103 | 0.00986 |
| K00806 | 2.17493 | 0.01922581 | K06966 | 3.15232 | 0.01404 |
| K00817 | -1.9459 | 0.01432198 | K06970 | 3.19649 | 0.02702 |
| K00823 | 3.63908 | 0.02154063 | K06972 | 5.33988 | 0.00263 |
| K00832 | 3.85378 | 0.00591536 | K06975 | 2.66647 | 0.03211 |
| K00844 | 3.45559 | 0.01264793 | K06980 | 5.08701 | 0.00455 |
| K00847 | -2.5376 | 0.01160662 | K06997 | 2.94536 | 0.00531 |
| K00848 | 3.87993 | 0.00701675 | K07000 | 4.34602 | 0.00531 |
| K00852 | 3.07002 | 0.00586344 | K07014 | 2.74678 | 0.0192 |
| K00855 | 3.76677 | 0.00774545 | K07019 | 2.99547 | 0.01613 |
| K00858 | 2.0647 | 0.01736684 | K07026 | 4.37572 | 0.00649 |
| K00865 | 3.85457 | 0.00451314 | K07030 | 3.53154 | 0.00455 |
| K00867 | 3.96538 | 0.00454382 | K07034 | 3.59729 | 0.01106 |
| K00878 | -1.6871 | 0.04539774 | K07038 | 4.51647 | 0.00488 |
| K00879 | 2.50563 | 0.04381744 | K07039 | 2.65553 | 0.03377 |
| K00880 | 4.19771 | 0.00656207 | K07040 | -2.0909 | 0.02576 |
| K00882 | 2.11349 | 0.02815799 | K07043 | 6.59314 | 0.00232 |
| K00883 | 3.40197 | 0.03474338 | K07051 | -6.1276 | 0.00286 |
| K00884 | 3.8766 | 0.01391732 | K07053 | 4.4522 | 0.00293 |
| K00885 | 4.40903 | 0.00844667 | K07054 | 2.40921 | 0.02557 |
| K00926 | 2.5516 | 0.01301388 | K07058 | 2.94963 | 0.01312 |
| K00930 | -1.6701 | 0.01705731 | K07067 | 5.69503 | 0.00232 |
| K00931 | 1.6916 | 0.02577995 | K07071 | 4.91507 | 0.00488 |
| K00932 | 3.13582 | 0.01844574 | K07080 | 2.99723 | 0.01175 |
| K00937 | 2.69735 | 0.01583296 | K07082 | 3.51681 | 0.00854 |
| K00950 | 2.62936 | 0.0301977 | K07088 | 1.98118 | 0.00922 |
| K00957 | -1.7052 | 0.04220926 | K07091 | 4.05259 | 0.00512 |
| K00962 | -1.2368 | 0.03594765 | K07095 | 1.96999 | 0.03582 |

| | | | | | |
|---|---|---|---|---|---|
| K00970 | -5.2516 | 0.00376385 | K07099 | 2.51894 | 0.01262 |
| K00974 | 2.55694 | 0.02244835 | K07101 | 3.60613 | 0.01685 |
| K00978 | -3.3581 | 0.01022454 | K07105 | 2.60554 | 0.03941 |
| K00986 | -4.2251 | 0.0075123 | K07107 | -2.0962 | 0.01106 |
| K00989 | 2.84585 | 0.00761685 | K07109 | 2.61777 | 0.04636 |
| K00990 | 2.75636 | 0.03120174 | K07112 | 3.82373 | 0.01024 |
| K00997 | 2.38699 | 0.01460036 | K07113 | 2.34131 | 0.03955 |
| K00998 | 3.44101 | 0.01838941 | K07114 | -1.4545 | 0.03739 |
| K01002 | 3.3357 | 0.01369034 | K07119 | 4.85432 | 0.00492 |
| K01008 | 4.51858 | 0.00359047 | K07121 | 2.82355 | 0.02853 |
| K01012 | -1.7386 | 0.01344724 | K07122 | 3.6452 | 0.00586 |
| K01028 | 3.20768 | 0.02708639 | K07124 | -2.0052 | 0.04215 |
| K01034 | 2.85612 | 0.02175034 | K07127 | 2.7234 | 0.04305 |
| K01035 | 3.08852 | 0.00850519 | K07132 | 4.26752 | 0.00852 |
| K01040 | 4.1662 | 0.0126136 | K07133 | -1.6221 | 0.0212 |
| K01042 | 5.49185 | 0.0023744 | K07136 | 2.54023 | 0.04631 |
| K01048 | 5.12893 | 0.00376385 | K07137 | -2.291 | 0.0084 |
| K01055 | 3.32789 | 0.03099263 | K07141 | 2.02232 | 0.03686 |
| K01058 | 3.98195 | 0.00607733 | K07146 | 4.25338 | 0.00762 |
| K01060 | 3.83827 | 0.0084728 | K07147 | 3.66922 | 0.01374 |
| K01066 | 3.46934 | 0.00992189 | K07149 | -2.6659 | 0.04426 |
| K01070 | 3.60763 | 0.01334109 | K07156 | 3.3807 | 0.02062 |
| K01082 | 2.65551 | 0.02779866 | K07171 | -1.6437 | 0.01949 |
| K01085 | 3.80718 | 0.00994413 | K07173 | 1.75817 | 0.02957 |
| K01087 | 3.91185 | 0.00824052 | K07175 | 3.28171 | 0.01733 |
| K01092 | -2.9137 | 0.01369034 | K07180 | 5.8371 | 0.00232 |
| K01104 | 2.11006 | 0.01349049 | K07184 | 2.99209 | 0.03549 |
| K01118 | 6.84331 | 0.00232026 | K07216 | 3.52075 | 0.01539 |
| K01119 | 4.28107 | 0.00761685 | K07229 | 2.89907 | 0.02816 |
| K01130 | -5.6458 | 0.00371604 | K07232 | 4.33211 | 0.00762 |
| K01138 | 5.0102 | 0.00451314 | K07235 | 4.29796 | 0.00649 |
| K01139 | 4.26026 | 0.00499871 | K07236 | 4.2631 | 0.00829 |
| K01141 | 3.78173 | 0.00774545 | K07248 | 2.81448 | 0.03041 |
| K01142 | 1.82163 | 0.0137402 | K07258 | 2.42219 | 0.00751 |
| K01146 | 3.14667 | 0.0243878 | K07259 | 3.46419 | 0.01997 |
| K01147 | 3.72856 | 0.00855169 | K07261 | 5.27122 | 0.00376 |
| K01150 | 3.26883 | 0.0230307 | K07262 | 3.80037 | 0.00777 |
| K01151 | -2.1072 | 0.00973872 | K07264 | 2.74479 | 0.0485 |
| K01155 | -4.9747 | 0.00232026 | K07273 | 2.58453 | 0.00854 |
| K01159 | -1.7311 | 0.01296947 | K07274 | 4.29536 | 0.00618 |
| K01163 | 2.04588 | 0.0215662 | K07275 | 4.16363 | 0.00748 |
| K01169 | 3.6826 | 0.01369034 | K07278 | 2.85905 | 0.04372 |
| K01193 | 2.03255 | 0.02460133 | K07284 | 3.77271 | 0.00352 |
| K01195 | 6.66106 | 0.00232026 | K07285 | 2.92342 | 0.02645 |
| K01197 | 6.79414 | 0.00232026 | K07289 | 2.73799 | 0.043 |
| K01205 | 3.58559 | 0.01972481 | K07300 | 5.20865 | 0.00303 |
| K01220 | 3.94582 | 0.00867507 | K07301 | 2.34808 | 0.01918 |
| K01226 | 6.62142 | 0.00232026 | K07304 | 3.39108 | 0.0058 |

| | | | | | |
|---|---|---|---|---|---|
| K01235 | 3.57336 | 0.01328107 | K07305 | 4.91584 | 0.0037 |
| K01246 | -2.311 | 0.00586344 | K07306 | 3.36463 | 0.011 |
| K01250 | 4.77077 | 0.00586344 | K07307 | 3.49785 | 0.01369 |
| K01252 | 3.74892 | 0.00761685 | K07311 | 2.45187 | 0.03288 |
| K01255 | 4.49438 | 0.00272423 | K07312 | 4.3032 | 0.00845 |
| K01273 | 3.6778 | 0.01216818 | K07313 | 2.76953 | 0.0309 |
| K01297 | 1.60227 | 0.02782427 | K07318 | 2.7343 | 0.03549 |
| K01304 | 2.14054 | 0.03035715 | K07320 | 2.24557 | 0.03314 |
| K01305 | 3.26457 | 0.01022454 | K07323 | 3.42142 | 0.00775 |
| K01355 | 4.20933 | 0.0078317 | K07337 | 3.94456 | 0.01277 |
| K01356 | 2.07319 | 0.01540485 | K07339 | -3.0687 | 0.01318 |
| K01358 | -1.1425 | 0.02995981 | K07341 | -3.0708 | 0.011 |
| K01387 | 5.32833 | 0.00262888 | K07347 | 3.12403 | 0.0154 |
| K01414 | 3.35092 | 0.0122644 | K07348 | 3.64998 | 0.01763 |
| K01420 | -3.3911 | 0.00446995 | K07351 | 2.88009 | 0.02039 |
| K01421 | 6.03005 | 0.00232026 | K07354 | 2.97143 | 0.02966 |
| K01423 | 3.58934 | 0.00761685 | K07356 | 3.33861 | 0.0194 |
| K01442 | 1.57448 | 0.02713785 | K07387 | 2.02245 | 0.04425 |
| K01443 | 3.04252 | 0.00673221 | K07391 | 2.68582 | 0.02866 |
| K01447 | -7.1538 | 0.00232026 | K07397 | 3.32217 | 0.0107 |
| K01448 | 1.97929 | 0.02892184 | K07399 | 2.83742 | 0.04878 |
| K01462 | 1.53443 | 0.02482213 | K07400 | 4.19652 | 0.00854 |
| K01466 | 3.57161 | 0.0151405 | K07402 | 5.00273 | 0.0033 |
| K01467 | 4.24315 | 0.00609582 | K07405 | -3.2112 | 0.01816 |
| K01478 | 3.04613 | 0.00854133 | K07448 | -4.1662 | 0.006 |
| K01480 | 3.59952 | 0.00530562 | K07455 | 5.83037 | 0.00232 |
| K01484 | 3.96761 | 0.00854133 | K07459 | 4.09141 | 0.00634 |
| K01486 | 4.6259 | 0.00371482 | K07461 | 3.12403 | 0.00593 |
| K01488 | 3.27173 | 0.00629648 | K07462 | 3.38044 | 0.00585 |
| K01491 | -1.0308 | 0.04940258 | K07479 | 3.37266 | 0.01438 |
| K01494 | 3.26171 | 0.01732737 | K07481 | -4.0876 | 0.00454 |
| K01497 | 3.40484 | 0.01979006 | K07490 | 4.71301 | 0.00545 |
| K01512 | 2.31431 | 0.01903483 | K07533 | 2.40958 | 0.01574 |
| K01521 | 3.83365 | 0.00992189 | K07552 | 3.88211 | 0.00531 |
| K01525 | 2.86712 | 0.04603421 | K07559 | 4.71092 | 0.00313 |
| K01531 | 5.33144 | 0.00269654 | K07560 | -1.5502 | 0.01229 |
| K01537 | 2.02898 | 0.01555735 | K07566 | -1.8453 | 0.01758 |
| K01571 | -3.1742 | 0.00635964 | K07568 | -1.4527 | 0.01839 |
| K01577 | 2.8781 | 0.02263964 | K07571 | 3.83832 | 0.00565 |
| K01581 | 2.72506 | 0.03474338 | K07589 | 3.6186 | 0.01468 |
| K01582 | 3.61496 | 0.00986263 | K07590 | 6.04329 | 0.00232 |
| K01590 | 5.64314 | 0.00232026 | K07592 | 3.96869 | 0.01052 |
| K01595 | 2.34487 | 0.02281156 | K07637 | 3.36845 | 0.01879 |
| K01599 | 3.33171 | 0.01128185 | K07638 | 4.1425 | 0.00543 |
| K01609 | 2.5866 | 0.04396872 | K07643 | 4.05563 | 0.00797 |
| K01611 | 4.80846 | 0.00379344 | K07659 | 3.82517 | 0.00838 |
| K01639 | 1.81084 | 0.03151284 | K07663 | 4.98377 | 0.00488 |
| K01640 | 2.55079 | 0.03437897 | K07666 | 3.76879 | 0.00586 |

| | | | | | |
|---|---|---|---|---|---|
| K01641 | 3.32381 | 0.03955991 | K07667 | 1.89253 | 0.04427 |
| K01643 | 2.86398 | 0.00844667 | K07674 | 3.50789 | 0.01824 |
| K01644 | 3.62543 | 0.00530562 | K07687 | 3.72704 | 0.01628 |
| K01646 | 1.77109 | 0.04846209 | K07688 | 4.4572 | 0.00854 |
| K01659 | 3.13198 | 0.0403171 | K07689 | 3.4471 | 0.02177 |
| K01664 | 4.42583 | 0.00775387 | K07690 | 3.4755 | 0.01475 |
| K01665 | 3.81364 | 0.00839319 | K07701 | 3.70325 | 0.01149 |
| K01666 | 3.70455 | 0.00983576 | K07708 | 3.54429 | 0.00853 |
| K01673 | 3.12856 | 0.00861747 | K07711 | 2.84967 | 0.02568 |
| K01679 | 2.07329 | 0.04425244 | K07715 | 3.36172 | 0.01907 |
| K01682 | 2.93992 | 0.03605912 | K07738 | -1.9798 | 0.00748 |
| K01684 | 3.74695 | 0.00933887 | K07740 | 3.4649 | 0.02645 |
| K01687 | -1.4815 | 0.01262288 | K07749 | 2.21742 | 0.03299 |
| K01693 | -2.7358 | 0.01391732 | K07751 | 3.90976 | 0.00992 |
| K01695 | -1.4391 | 0.04371826 | K07771 | 4.49878 | 0.00593 |
| K01698 | 2.09412 | 0.03897742 | K07772 | 3.13646 | 0.02459 |
| K01703 | -2.1253 | 0.00734522 | K07773 | 2.85099 | 0.04812 |
| K01709 | -3.1248 | 0.03838058 | K07782 | 3.34273 | 0.02838 |
| K01715 | 3.12062 | 0.0121596 | K07783 | 4.1477 | 0.01946 |
| K01716 | 3.29062 | 0.02752338 | K07784 | 4.44144 | 0.00634 |
| K01720 | 3.45632 | 0.0136338 | K07791 | 3.86712 | 0.00491 |
| K01725 | 4.15073 | 0.008011 | K07793 | 3.79932 | 0.00644 |
| K01738 | 1.77143 | 0.02932894 | K07798 | 2.17764 | 0.04913 |
| K01740 | -2.5797 | 0.00844667 | K07799 | 4.27196 | 0.00748 |
| K01744 | 3.411 | 0.01036241 | K07803 | 3.15863 | 0.0277 |
| K01749 | 4.51816 | 0.00272423 | K07806 | 4.35585 | 0.00684 |
| K01753 | 2.99566 | 0.03173186 | K07810 | 3.97256 | 0.01518 |
| K01760 | 3.63819 | 0.00623929 | K07812 | 2.63606 | 0.04419 |
| K01766 | 3.50427 | 0.01702879 | K07813 | 4.63626 | 0.0033 |
| K01770 | -1.4077 | 0.01073472 | K08092 | 4.56289 | 0.00543 |
| K01775 | 2.81442 | 0.00586344 | K08137 | 2.89942 | 0.02702 |
| K01776 | 2.57206 | 0.00850519 | K08138 | 3.09377 | 0.01955 |
| K01782 | 3.79144 | 0.00870367 | K08154 | 3.72731 | 0.01224 |
| K01783 | 3.28556 | 0.00446995 | K08156 | -2.688 | 0.00546 |
| K01787 | -2.9261 | 0.01922581 | K08160 | 3.98554 | 0.00871 |
| K01792 | 3.15314 | 0.02459467 | K08161 | 2.20073 | 0.03433 |
| K01816 | 3.29478 | 0.01465597 | K08162 | 2.82671 | 0.03064 |
| K01818 | -2.2135 | 0.00687618 | K08163 | 4.3058 | 0.00916 |
| K01823 | 3.18015 | 0.01541818 | K08172 | 3.1642 | 0.01927 |
| K01838 | 3.62666 | 0.01552803 | K08177 | 4.75759 | 0.00372 |
| K01858 | 2.67959 | 0.04081331 | K08191 | 4.71437 | 0.00579 |
| K01890 | 2.06961 | 0.01653078 | K08217 | -1.5515 | 0.02803 |
| K01897 | 2.15291 | 0.02577995 | K08219 | 3.84715 | 0.00974 |
| K01903 | -4.6492 | 0.00542959 | K08221 | -2.7755 | 0.03823 |
| K01908 | 3.40547 | 0.01930561 | K08222 | 3.26191 | 0.02355 |
| K01910 | 2.33777 | 0.02398133 | K08224 | 5.12761 | 0.00455 |
| K01911 | 2.53249 | 0.02234516 | K08227 | 2.68822 | 0.0424 |
| K01918 | -1.9367 | 0.00946992 | K08281 | 3.75013 | 0.00899 |

| | | | | | |
|---|---|---|---|---|---|
| K01919 | 2.7966 | 0.01137203 | K08296 | 4.36781 | 0.00634 |
| K01921 | 1.59934 | 0.03092987 | K08299 | 4.55437 | 0.00762 |
| K01923 | -1.7332 | 0.01667142 | K08301 | 3.32284 | 0.00868 |
| K01929 | 3.03618 | 0.00542926 | K08304 | 2.42688 | 0.02635 |
| K01958 | 3.50715 | 0.01681914 | K08306 | 3.74937 | 0.01349 |
| K01962 | 2.49652 | 0.01304261 | K08307 | 3.06785 | 0.00992 |
| K01963 | 2.95937 | 0.00796851 | K08310 | 4.09053 | 0.00608 |
| K01974 | 4.86047 | 0.00330101 | K08311 | 3.88364 | 0.00957 |
| K01975 | 4.0074 | 0.00665826 | K08314 | 4.48089 | 0.00634 |
| K01994 | -4.047 | 0.02384074 | K08315 | 4.80899 | 0.0061 |
| K01999 | 4.08213 | 0.00586344 | K08320 | 2.86696 | 0.03824 |
| K02000 | 2.93629 | 0.03182165 | K08326 | 3.55145 | 0.01334 |
| K02003 | -1.0938 | 0.0326321 | K08349 | 4.07018 | 0.0087 |
| K02006 | 3.01694 | 0.00951193 | K08351 | 3.96895 | 0.00851 |
| K02007 | -3.4668 | 0.01037298 | K08368 | 4.11163 | 0.00603 |
| K02008 | 5.12835 | 0.00262888 | K08384 | 2.52245 | 0.03263 |
| K02009 | 4.05651 | 0.00454382 | K08483 | 4.0597 | 0.0033 |
| K02010 | 2.95166 | 0.04483259 | K08485 | 5.25716 | 0.0033 |
| K02017 | 3.69527 | 0.00607733 | K08587 | 5.18219 | 0.00362 |
| K02018 | 2.24697 | 0.04304573 | K08591 | 2.2951 | 0.01247 |
| K02019 | 3.15521 | 0.01493172 | K08602 | 2.0195 | 0.0385 |
| K02020 | 3.68686 | 0.0084524 | K08641 | 3.14914 | 0.01349 |
| K02022 | -5.3913 | 0.00470949 | K08678 | -2.6091 | 0.03959 |
| K02024 | 2.99758 | 0.03524855 | K08722 | 2.80584 | 0.02971 |
| K02026 | -1.6384 | 0.03315961 | K08723 | 2.90771 | 0.02702 |
| K02032 | -3.9671 | 0.00609582 | K08884 | 3.79947 | 0.01342 |
| K02034 | 1.6066 | 0.02769564 | K08970 | 3.33228 | 0.02822 |
| K02035 | 2.11307 | 0.01948865 | K08972 | 2.97987 | 0.04353 |
| K02042 | 3.58985 | 0.00916474 | K08974 | 4.62521 | 0.00379 |
| K02044 | 2.23098 | 0.0301977 | K08981 | 5.7735 | 0.00232 |
| K02045 | 3.08156 | 0.03636892 | K08984 | 3.62023 | 0.01764 |
| K02046 | 3.97819 | 0.00854133 | K08986 | 5.85936 | 0.00232 |
| K02047 | 4.99111 | 0.00446995 | K08989 | 4.34322 | 0.00762 |
| K02049 | 2.17853 | 0.02966164 | K08990 | 4.51486 | 0.0061 |
| K02051 | 3.5463 | 0.00451314 | K08992 | 4.08664 | 0.01409 |
| K02055 | 2.8432 | 0.02296519 | K08993 | 4.70138 | 0.00735 |
| K02056 | 2.02732 | 0.02526491 | K09001 | 2.39602 | 0.03441 |
| K02058 | 2.77876 | 0.02702067 | K09007 | -5.3934 | 0.00481 |
| K02062 | 2.91048 | 0.0230307 | K09009 | 2.64248 | 0.04372 |
| K02066 | -3.0511 | 0.01054403 | K09018 | 2.75483 | 0.03519 |
| K02068 | 3.64902 | 0.00854133 | K09020 | 3.05702 | 0.02751 |
| K02072 | 2.57763 | 0.01250024 | K09023 | 2.79088 | 0.03263 |
| K02078 | -1.5019 | 0.0128364 | K09117 | 2.65318 | 0.02416 |
| K02079 | 3.94556 | 0.00796036 | K09118 | 5.67284 | 0.00232 |
| K02081 | 4.5276 | 0.00451314 | K09121 | -3.8469 | 0.00779 |
| K02083 | 3.05266 | 0.02153984 | K09124 | 3.22632 | 0.02493 |
| K02099 | 2.70705 | 0.0456527 | K09128 | 3.99444 | 0.01774 |
| K02100 | 3.76824 | 0.00870367 | K09131 | 3.84123 | 0.00868 |

| | | | | | |
|---|---|---|---|---|---|
| K02106 | 3.65282 | 0.00839813 | K09137 | 5.67583 | 0.00232 |
| K02107 | 2.96054 | 0.04753758 | K09158 | 3.89087 | 0.00818 |
| K02110 | -1.5556 | 0.0125352 | K09159 | 4.97517 | 0.00358 |
| K02116 | 5.46929 | 0.00232026 | K09160 | 4.44665 | 0.0087 |
| K02122 | 3.2934 | 0.00761685 | K09161 | 3.54866 | 0.01433 |
| K02160 | 1.882 | 0.0426263 | K09167 | 3.87822 | 0.01755 |
| K02167 | 4.92959 | 0.00488139 | K09181 | 3.46642 | 0.00845 |
| K02170 | 4.03852 | 0.00584564 | K09470 | 4.63945 | 0.00543 |
| K02172 | -3.8958 | 0.00488139 | K09471 | 3.54335 | 0.01819 |
| K02189 | 5.53262 | 0.00232026 | K09473 | 3.07605 | 0.02636 |
| K02192 | 4.18703 | 0.01124209 | K09702 | -1.7195 | 0.04291 |
| K02193 | 3.72964 | 0.00609582 | K09705 | 4.77506 | 0.00631 |
| K02194 | 3.05516 | 0.02419945 | K09712 | 3.72242 | 0.01044 |
| K02195 | 4.30328 | 0.00586344 | K09758 | 5.11259 | 0.0033 |
| K02196 | 3.95598 | 0.00992189 | K09760 | 2.34285 | 0.0493 |
| K02198 | 2.93169 | 0.03092987 | K09762 | 2.30806 | 0.0206 |
| K02199 | 2.52412 | 0.02306038 | K09764 | -2.5071 | 0.04671 |
| K02200 | 2.57296 | 0.03598703 | K09767 | 4.2779 | 0.00845 |
| K02221 | 2.9897 | 0.01742986 | K09770 | -3.7585 | 0.01428 |
| K02227 | 4.2829 | 0.00376385 | K09771 | 5.7502 | 0.00272 |
| K02237 | 2.42208 | 0.01369712 | K09773 | 3.14791 | 0.00592 |
| K02245 | 2.47703 | 0.04287432 | K09778 | 5.1971 | 0.00313 |
| K02255 | 3.15125 | 0.02877878 | K09779 | -3.0369 | 0.01428 |
| K02257 | 3.56859 | 0.01224322 | K09786 | 5.84329 | 0.00232 |
| K02283 | -4.4428 | 0.00820127 | K09787 | 1.78573 | 0.03133 |
| K02297 | 3.32122 | 0.01342397 | K09791 | 3.94707 | 0.00992 |
| K02299 | 4.80732 | 0.00469659 | K09802 | 2.89609 | 0.04382 |
| K02300 | 3.14466 | 0.02474668 | K09803 | 3.42275 | 0.02999 |
| K02315 | -1.6231 | 0.02774825 | K09806 | 4.14228 | 0.00725 |
| K02317 | 3.78339 | 0.00928971 | K09811 | 2.50636 | 0.01924 |
| K02337 | -1.7722 | 0.02141566 | K09812 | 1.82481 | 0.03093 |
| K02339 | 4.68896 | 0.00530562 | K09817 | 3.4239 | 0.01574 |
| K02344 | 2.99354 | 0.04735692 | K09819 | -6.3092 | 0.00293 |
| K02358 | -1.0149 | 0.01816438 | K09824 | 3.8446 | 0.01052 |
| K02361 | 3.89538 | 0.01310614 | K09858 | 4.82583 | 0.00638 |
| K02363 | 4.46387 | 0.00586344 | K09862 | 3.80626 | 0.00779 |
| K02379 | 4.12396 | 0.00635964 | K09889 | 3.01536 | 0.01518 |
| K02380 | 4.68304 | 0.00475177 | K09890 | 4.09085 | 0.01711 |
| K02386 | 3.68676 | 0.01149007 | K09891 | 3.43969 | 0.01105 |
| K02389 | 3.34792 | 0.01864367 | K09892 | 5.54032 | 0.00282 |
| K02392 | 2.5318 | 0.02535226 | K09893 | 3.64482 | 0.008 |
| K02395 | 2.73388 | 0.03954487 | K09894 | 3.87228 | 0.01158 |
| K02396 | 2.68525 | 0.02355585 | K09896 | 3.02558 | 0.02989 |
| K02398 | 3.00394 | 0.02284259 | K09897 | 4.67877 | 0.00775 |
| K02399 | 3.28985 | 0.0230307 | K09898 | 5.27871 | 0.00379 |
| K02402 | 3.89958 | 0.01264965 | K09899 | 3.63863 | 0.01045 |
| K02406 | -1.7754 | 0.04765436 | K09900 | 5.12306 | 0.00467 |
| K02408 | 2.07715 | 0.04114735 | K09901 | 3.53996 | 0.01514 |

| | | | | | |
|---|---|---|---|---|---|
| K02409 | 3.05422 | 0.03634269 | K09902 | 5.08652 | 0.0033 |
| K02411 | 3.63278 | 0.01626751 | K09904 | 4.23201 | 0.00723 |
| K02414 | 4.92848 | 0.0035602 | K09906 | 3.35391 | 0.0382 |
| K02424 | 2.83352 | 0.01862745 | K09908 | 3.22329 | 0.03093 |
| K02426 | 2.70941 | 0.02920023 | K09910 | 3.26847 | 0.01711 |
| K02427 | 3.43309 | 0.00780222 | K09911 | 3.51982 | 0.01702 |
| K02428 | -1.319 | 0.02038726 | K09913 | 4.81732 | 0.00735 |
| K02431 | 2.87618 | 0.02411298 | K09915 | 3.15018 | 0.0273 |
| K02435 | -2.2471 | 0.03470931 | K09916 | 2.6992 | 0.03708 |
| K02438 | 2.86152 | 0.02819521 | K09917 | 4.94533 | 0.00796 |
| K02439 | 3.77596 | 0.00670795 | K09918 | 3.4402 | 0.0393 |
| K02440 | 2.00834 | 0.02912542 | K09920 | 4.26154 | 0.00603 |
| K02441 | 3.87927 | 0.0075123 | K09921 | 3.52022 | 0.00671 |
| K02446 | 2.81663 | 0.03253897 | K09925 | 2.97871 | 0.0393 |
| K02459 | 4.20111 | 0.0080935 | K09933 | 2.51847 | 0.04604 |
| K02466 | 2.6756 | 0.04396872 | K09935 | 4.31044 | 0.0061 |
| K02471 | 2.59754 | 0.04758469 | K09946 | 3.36997 | 0.02122 |
| K02472 | 4.14123 | 0.0072473 | K09954 | 3.95186 | 0.01769 |
| K02477 | 2.96725 | 0.02474751 | K09969 | 3.89246 | 0.00563 |
| K02483 | 4.41277 | 0.00656816 | K09970 | 3.86354 | 0.00631 |
| K02485 | 3.17011 | 0.01783911 | K09971 | 3.00332 | 0.02081 |
| K02494 | 3.8354 | 0.00701675 | K09973 | 5.15847 | 0.00272 |
| K02495 | 4.10666 | 0.00552251 | K09975 | 4.01497 | 0.0087 |
| K02499 | 2.84485 | 0.01541818 | K09979 | 3.80404 | 0.01763 |
| K02500 | -1.6843 | 0.02419945 | K09996 | 3.88176 | 0.01755 |
| K02507 | 2.63744 | 0.04846209 | K09999 | 4.56407 | 0.00586 |
| K02510 | 3.12406 | 0.01218982 | K10001 | 3.18556 | 0.02297 |
| K02518 | -1.0819 | 0.02559826 | K10004 | 3.44229 | 0.01037 |
| K02521 | 3.44009 | 0.01790324 | K10009 | 2.5731 | 0.01883 |
| K02523 | 4.16972 | 0.00541922 | K10011 | 2.84196 | 0.03988 |
| K02526 | 2.83739 | 0.0243878 | K10012 | 3.7665 | 0.00865 |
| K02527 | 2.85402 | 0.02041967 | K10015 | 3.81163 | 0.00776 |
| K02532 | 3.11005 | 0.01674505 | K10017 | 4.56225 | 0.00586 |
| K02533 | 2.78877 | 0.01754712 | K10036 | 2.60756 | 0.03097 |
| K02535 | 3.18658 | 0.01589521 | K10110 | 4.87643 | 0.00491 |
| K02549 | 2.68673 | 0.03581696 | K10111 | 3.79459 | 0.00797 |
| K02551 | -2.3559 | 0.01824322 | K10439 | 1.80663 | 0.02822 |
| K02557 | 3.02907 | 0.01404277 | K10441 | 1.64903 | 0.04671 |
| K02558 | 3.83071 | 0.00586344 | K10537 | 3.14691 | 0.01949 |
| K02560 | 3.10619 | 0.02473517 | K10543 | 2.61433 | 0.03299 |
| K02565 | 2.81425 | 0.03446515 | K10544 | 3.59284 | 0.01035 |
| K02566 | 5.03422 | 0.00530562 | K10555 | 3.64874 | 0.01219 |
| K02567 | 3.40291 | 0.00987284 | K10558 | 2.57703 | 0.04964 |
| K02570 | 4.41102 | 0.0075883 | K10563 | 2.09054 | 0.03184 |
| K02572 | 2.56166 | 0.02850525 | K10680 | 4.05905 | 0.00862 |
| K02573 | 2.95902 | 0.01617968 | K10709 | 4.82391 | 0.0047 |
| K02574 | 3.69211 | 0.00673221 | K10711 | 3.63051 | 0.01975 |
| K02575 | 2.47568 | 0.02434996 | K10716 | 1.82781 | 0.04953 |

| | | | | | |
|---|---|---|---|---|---|
| K02588 | 3.95879 | 0.00808742 | K10763 | 3.47171 | 0.0124 |
| K02598 | 2.77707 | 0.03881571 | K10778 | 3.82255 | 0.01353 |
| K02610 | 4.50336 | 0.00900665 | K10797 | 2.94642 | 0.03435 |
| K02611 | 3.89146 | 0.00854133 | K10804 | 4.56409 | 0.00531 |
| K02617 | 3.40826 | 0.02355585 | K10914 | 3.61974 | 0.00845 |
| K02650 | 4.75272 | 0.00422722 | K10939 | 3.36961 | 0.02435 |
| K02654 | 2.92015 | 0.03379377 | K10973 | 4.03555 | 0.00671 |
| K02664 | 5.41299 | 0.00253596 | K10974 | 3.81065 | 0.00905 |
| K02669 | 2.04689 | 0.04425244 | K11057 | 6.51881 | 0.00232 |
| K02679 | 5.28796 | 0.00379344 | K11065 | 3.23959 | 0.00805 |
| K02681 | 5.04795 | 0.00586344 | K11066 | 3.29813 | 0.01767 |
| K02682 | 3.57563 | 0.01026489 | K11073 | 3.90187 | 0.00787 |
| K02686 | 6.04059 | 0.00232026 | K11076 | 3.27567 | 0.02554 |
| K02742 | 3.36805 | 0.0220474 | K11102 | 5.47703 | 0.00232 |
| K02745 | 3.05309 | 0.01301388 | K11104 | 2.87481 | 0.02371 |
| K02750 | 2.99883 | 0.02395613 | K11106 | 3.81922 | 0.00845 |
| K02768 | 3.46851 | 0.0043623 | K11139 | 2.99844 | 0.03143 |
| K02769 | 2.48204 | 0.01554598 | K11175 | 2.34198 | 0.01322 |
| K02770 | 3.35 | 0.00542926 | K11178 | 3.60776 | 0.011 |
| K02775 | 2.81014 | 0.02662682 | K11183 | 3.8304 | 0.00735 |
| K02777 | 3.00411 | 0.01115265 | K11191 | 2.83293 | 0.03263 |
| K02784 | 3.65096 | 0.01426131 | K11192 | 2.83293 | 0.03263 |
| K02796 | 2.43888 | 0.01965658 | K11201 | 3.37853 | 0.00986 |
| K02798 | 3.61826 | 0.00644444 | K11202 | 2.95246 | 0.01101 |
| K02799 | 5.10706 | 0.00330101 | K11203 | 3.45991 | 0.01224 |
| K02800 | 5.10706 | 0.00330101 | K11209 | 4.52719 | 0.00465 |
| K02803 | 3.10697 | 0.03286657 | K11264 | 4.79863 | 0.0047 |
| K02806 | 3.39848 | 0.00867507 | K11392 | 3.04485 | 0.02816 |
| K02818 | 5.03129 | 0.00526331 | K11472 | 4.21211 | 0.00762 |
| K02819 | 4.91306 | 0.00542926 | K11473 | 2.62249 | 0.0454 |
| K02821 | 5.05973 | 0.00272423 | K11474 | 3.92591 | 0.00986 |
| K02823 | 6.05331 | 0.00232026 | K11477 | 3.43988 | 0.02625 |
| K02835 | -1.052 | 0.03891044 | K11534 | 3.71537 | 0.00657 |
| K02836 | 1.41185 | 0.04771966 | K11535 | 3.96813 | 0.00845 |
| K02838 | -1.35 | 0.01809457 | K11615 | 2.96357 | 0.03529 |
| K02841 | 2.30125 | 0.04114735 | K11685 | 3.76872 | 0.0094 |
| K02846 | 3.83955 | 0.01109856 | K11719 | 4.49702 | 0.00531 |
| K02851 | 2.23404 | 0.03563365 | K11734 | 3.84963 | 0.00868 |
| K02856 | 4.58361 | 0.00530562 | K11735 | 3.12327 | 0.02657 |
| K02858 | 4.37707 | 0.00530562 | K11741 | 3.14899 | 0.03424 |
| K02867 | -1.1965 | 0.02039189 | K11742 | 2.71306 | 0.04221 |
| K02874 | -1.4673 | 0.00916474 | K11745 | 2.61503 | 0.04494 |
| K02876 | -1.2177 | 0.01483334 | K11747 | 4.18383 | 0.00929 |
| K02878 | -1.1169 | 0.02142701 | K11748 | 4.37036 | 0.00656 |
| K02879 | -1.1436 | 0.02416593 | K11750 | 3.31602 | 0.01189 |
| K02887 | -1.0353 | 0.02296443 | K11751 | 3.1869 | 0.02154 |
| K02890 | -0.9964 | 0.04468668 | K11752 | 2.91343 | 0.02753 |
| K02892 | -1.038 | 0.04940258 | K11921 | 3.3217 | 0.02534 |

| | | |
|---|---|---|
| K02897 | -1.2904 | 0.0326321 |
| K02899 | -1.2949 | 0.01369034 |
| K02919 | -2.4707 | 0.0354711 |
| K02933 | -1.2465 | 0.02061651 |
| K02946 | -1.3313 | 0.01158949 |
| K02948 | -1.1996 | 0.02460164 |
| K02952 | -1.0803 | 0.03076157 |
| K02959 | -1.2975 | 0.0212226 |
| K02965 | -1.5713 | 0.00629648 |
| K02970 | -1.1296 | 0.03378532 |
| K02982 | -1.3295 | 0.01027704 |
| K02992 | -1.4399 | 0.00906364 |
| K03046 | -1.1305 | 0.03672708 |
| K03060 | 2.62037 | 0.00586344 |
| K03071 | 4.20748 | 0.00536711 |
| K03072 | 5.28869 | 0.00232026 |
| K03076 | -1.1245 | 0.04682172 |
| K03079 | 4.76474 | 0.00521987 |
| K03098 | 3.93731 | 0.00796851 |
| K03100 | 2.596 | 0.00629648 |
| K03101 | 2.59832 | 0.00776355 |
| K03116 | -2.341 | 0.01579517 |
| K03117 | 3.33407 | 0.00917432 |
| K03118 | -1.9423 | 0.01352505 |
| K03119 | 5.35878 | 0.00488139 |
| K03148 | 3.97625 | 0.00752797 |
| K03151 | 2.52522 | 0.00673221 |
| K03169 | -1.5179 | 0.0354711 |
| K03181 | 4.77739 | 0.00508927 |
| K03182 | 3.37536 | 0.01438165 |
| K03186 | 2.48563 | 0.04453913 |
| K03190 | 3.72994 | 0.02769564 |
| K03205 | -4.1641 | 0.00232026 |
| K03215 | 1.40033 | 0.03653552 |
| K03271 | -3.2733 | 0.00933648 |
| K03273 | 3.80585 | 0.00754087 |
| K03286 | 3.27845 | 0.01099713 |
| K03289 | 3.87221 | 0.00986263 |
| K03290 | 3.03012 | 0.03066748 |
| K03291 | 3.92952 | 0.00865166 |
| K03297 | 4.09484 | 0.00971033 |
| K03299 | 2.60163 | 0.03437897 |
| K03305 | 2.15145 | 0.04591323 |
| K03311 | 2.07899 | 0.04910859 |
| K03314 | 2.81368 | 0.04120844 |
| K03319 | 3.84209 | 0.00747638 |
| K03326 | 3.2838 | 0.03916421 |
| K03327 | 3.38842 | 0.01763264 |

| | | |
|---|---|---|
| K11922 | 4.16285 | 0.00779 |
| K11924 | 3.89015 | 0.01229 |
| K11926 | 2.64292 | 0.04291 |
| K11927 | 4.09924 | 0.0061 |
| K11929 | 3.83486 | 0.01247 |
| K11932 | 2.96243 | 0.03106 |
| K11933 | 3.47642 | 0.01404 |
| K11934 | 2.72029 | 0.04034 |
| K11938 | 2.62012 | 0.04657 |
| K11939 | 4.59636 | 0.00593 |
| K12056 | 2.5965 | 0.04765 |
| K12132 | 2.28108 | 0.03768 |
| K12138 | 3.80701 | 0.01374 |
| K12146 | 3.20454 | 0.02043 |
| K12147 | 3.64797 | 0.0083 |
| K12148 | 3.24324 | 0.02154 |
| K12151 | 2.80957 | 0.03256 |
| K12152 | 3.73281 | 0.01102 |
| K12264 | 2.77323 | 0.0278 |
| K12265 | 3.07309 | 0.02356 |
| K12266 | 2.75256 | 0.03525 |
| K12267 | -2.8063 | 0.01682 |
| K12291 | 2.85457 | 0.02459 |
| K12297 | 2.79623 | 0.0431 |
| K12299 | 3.40792 | 0.011 |
| K12340 | 4.37281 | 0.00586 |
| K12369 | 3.35945 | 0.01336 |
| K12371 | 4.06144 | 0.00793 |
| K12452 | -4.2347 | 0.00379 |
| K12507 | 3.3618 | 0.02039 |
| K12524 | 2.50413 | 0.03739 |
| K12527 | 4.12522 | 0.006 |
| K12529 | 3.84463 | 0.00845 |
| K12574 | 1.92072 | 0.02671 |
| K12582 | 3.39701 | 0.02482 |
| K12684 | 4.25844 | 0.01734 |
| K12940 | 5.23472 | 0.00293 |
| K12943 | 3.55827 | 0.011 |
| K12957 | 2.7551 | 0.03162 |
| K12961 | 5.39375 | 0.00338 |
| K12963 | 4.58218 | 0.00857 |
| K12972 | 3.7495 | 0.01785 |
| K12973 | 2.52702 | 0.02859 |
| K12974 | 3.16526 | 0.02091 |
| K13004 | 3.12497 | 0.02635 |
| K13012 | 4.03095 | 0.00921 |
| K13014 | 4.20062 | 0.00726 |
| K13038 | 2.59451 | 0.00901 |

| | | | | | |
|---|---|---|---|---|---|
| K03329 | 3.06043 | 0.0264104 | K13040 | 3.63454 | 0.04177 |
| K03335 | 3.33162 | 0.00635964 | K13043 | -2.0848 | 0.00629 |
| K03336 | 2.56688 | 0.014279 | K13069 | 3.50464 | 0.01453 |
| K03337 | 4.5191 | 0.00462472 | K13244 | 2.9979 | 0.03493 |
| K03338 | 5.6559 | 0.00232026 | K13255 | 5.71394 | 0.00273 |
| K03385 | 3.02242 | 0.01763672 | K13256 | 4.08835 | 0.00488 |
| K03386 | 3.53584 | 0.00644444 | K13281 | 5.39028 | 0.00272 |
| K03387 | 2.84645 | 0.03234244 | K13283 | 4.31128 | 0.00644 |
| K03399 | 5.38582 | 0.00317745 | K13292 | 2.98278 | 0.00608 |
| K03406 | -4.5091 | 0.00778852 | K13301 | 3.95403 | 0.01208 |
| K03414 | 3.85483 | 0.01106441 | K13483 | 3.86897 | 0.01102 |
| K03415 | -2.606 | 0.02577995 | K13497 | 2.61941 | 0.04574 |
| K03427 | -1.1803 | 0.02822195 | K13498 | 3.88855 | 0.01251 |
| K03429 | 2.52992 | 0.02841342 | K13542 | 3.43677 | 0.0063 |
| K03439 | 1.38082 | 0.03765589 | K13566 | 2.6959 | 0.03593 |
| K03446 | 2.90715 | 0.0342442 | K13574 | 2.5813 | 0.04372 |
| K03449 | 2.45958 | 0.03196874 | K13628 | 4.13778 | 0.0087 |
| K03458 | 5.81152 | 0.00232026 | K13631 | 4.28154 | 0.00897 |
| K03459 | 3.77044 | 0.01642675 | K13638 | 4.45032 | 0.00745 |
| K03465 | 3.50986 | 0.00726267 | K13639 | 5.03991 | 0.0042 |
| K03474 | -2.0529 | 0.01629137 | K13641 | 4.43574 | 0.00531 |
| K03477 | 2.56938 | 0.03891044 | K13643 | 2.75845 | 0.01878 |
| K03478 | 4.21197 | 0.00586344 | K13650 | 4.34023 | 0.01092 |
| K03480 | 6.4135 | 0.00232026 | K13686 | 3.40381 | 0.02195 |
| K03481 | 5.65222 | 0.00232026 | K13695 | 3.21842 | 0.02377 |
| K03485 | 4.22675 | 0.0075883 | K13771 | 4.17494 | 0.00657 |
| K03487 | 2.84641 | 0.0342442 | K13788 | 3.46426 | 0.00868 |
| K03488 | 2.73842 | 0.02434996 | K13890 | 3.51307 | 0.01349 |
| K03497 | -1.52 | 0.0111523 | K13891 | 3.49423 | 0.02311 |
| K03500 | 4.45108 | 0.00293278 | K13919 | 4.37252 | 0.00748 |
| K03501 | 1.54388 | 0.04554291 | K13920 | 3.50012 | 0.01369 |
| K03516 | 2.41253 | 0.03822876 | K13922 | 4.10639 | 0.00868 |
| K03521 | -2.5374 | 0.00369568 | K13953 | 4.64222 | 0.00586 |
| K03522 | 1.85744 | 0.03524855 | K13954 | 4.74837 | 0.00522 |
| K03523 | 3.45378 | 0.00852725 | K13963 | 2.76064 | 0.02899 |
| K03526 | -1.3282 | 0.03813486 | K14056 | 2.43328 | 0.04591 |
| K03529 | 4.64639 | 0.00330101 | K14061 | 2.86147 | 0.04804 |
| K03531 | 2.27234 | 0.01343836 | K14062 | 4.03687 | 0.00839 |
| K03533 | 4.06444 | 0.01352505 | K14064 | 3.97541 | 0.01054 |
| K03543 | 2.85034 | 0.02932894 | K14065 | 3.69519 | 0.00649 |
| K03546 | 2.34734 | 0.04804367 | K14155 | 3.48007 | 0.00686 |
| K03547 | 2.24589 | 0.01358631 | K14170 | 3.87329 | 0.00543 |
| K03548 | 3.23255 | 0.02542672 | K14187 | 4.08435 | 0.00893 |
| K03549 | 1.80919 | 0.03850109 | K14261 | 2.69646 | 0.04671 |
| K03551 | -1.1257 | 0.02877878 | K14287 | 4.01941 | 0.0092 |
| K03557 | 3.50605 | 0.0151405 | K14347 | 3.86264 | 0.0084 |
| K03558 | 2.51824 | 0.03884027 | K14348 | 2.49889 | 0.04772 |
| K03560 | 3.54564 | 0.01310614 | K14414 | 4.74304 | 0.00379 |

| | | |
|---|---|---|
| K03562 | 3.69507 | 0.00867507 |
| K03564 | -1.5323 | 0.04461289 |
| K03566 | 4.20566 | 0.00774545 |
| K03568 | 4.3444 | 0.00339646 |
| K03569 | -1.2016 | 0.03739153 |
| K03571 | 1.87399 | 0.03323242 |
| K03573 | 2.77694 | 0.03682479 |
| K03575 | 3.38698 | 0.00830062 |
| K03576 | 3.7324 | 0.00882746 |
| K03578 | 2.64177 | 0.04158552 |
| K03579 | 3.96886 | 0.00983576 |
| K03582 | 3.02646 | 0.02183068 |
| K03583 | 2.91964 | 0.01921525 |
| K03586 | 2.90744 | 0.01767404 |
| K03587 | 3.6536 | 0.00793554 |
| K03592 | 3.90042 | 0.00499871 |
| K03593 | 2.35603 | 0.02877878 |
| K03597 | 3.06188 | 0.01838941 |
| K03598 | 2.8916 | 0.03120174 |
| K03599 | 2.77212 | 0.01661052 |
| K03603 | 2.4676 | 0.03959128 |
| K03605 | 3.51226 | 0.01038785 |
| K03607 | 4.05991 | 0.00774545 |
| K03608 | 3.59602 | 0.00542926 |
| K03610 | 5.04567 | 0.00285777 |
| K03612 | 1.83868 | 0.0365319 |
| K03618 | 3.77976 | 0.00829359 |
| K03619 | 4.13769 | 0.00701675 |
| K03623 | 2.72299 | 0.02679797 |
| K03628 | -3.5642 | 0.00628977 |
| K03630 | 1.7187 | 0.03708494 |
| K03632 | 2.78364 | 0.04371826 |
| K03633 | 3.42013 | 0.0150381 |
| K03634 | 3.67843 | 0.00992189 |
| K03635 | 2.58828 | 0.03133353 |
| K03637 | 3.79627 | 0.00530562 |
| K03639 | 5.03971 | 0.00235042 |
| K03640 | 2.60974 | 0.03011302 |
| K03641 | 4.4206 | 0.00521987 |
| K03642 | -5.2644 | 0.0032463 |
| K03643 | 4.04977 | 0.00986263 |
| K03645 | 4.05718 | 0.00586344 |
| K03646 | 2.61279 | 0.04631187 |
| K03651 | 2.85199 | 0.02107811 |
| K03652 | 2.48419 | 0.01537651 |
| K03656 | 4.47712 | 0.00551338 |
| K03657 | 2.00915 | 0.01112944 |
| K03658 | 2.9833 | 0.03850342 |
| K14441 | -1.5293 | 0.02853 |
| K14445 | -3.8419 | 0.00332 |
| K14540 | 2.36094 | 0.03287 |
| K14623 | -2.8484 | 0.02799 |
| K14682 | 3.2023 | 0.0168 |
| K14744 | 3.21157 | 0.02264 |
| K14762 | 3.23092 | 0.03424 |
| K15256 | 4.2534 | 0.00762 |
| K15461 | 3.50647 | 0.00796 |
| K15520 | 3.09279 | 0.02916 |
| K15524 | 2.79658 | 0.03654 |
| K15540 | 4.5603 | 0.00586 |
| K15547 | 3.88576 | 0.01028 |
| K15553 | 4.01533 | 0.00992 |
| K15554 | 4.14042 | 0.00656 |
| K15555 | 2.52591 | 0.04796 |
| K15580 | 2.39569 | 0.03683 |
| K15581 | -2.0124 | 0.0274 |
| K15583 | 2.03659 | 0.02666 |
| K15586 | 2.27757 | 0.0331 |
| K15633 | -1.3239 | 0.01607 |
| K15634 | 2.89444 | 0.02803 |
| K15722 | 2.5533 | 0.03838 |
| K15724 | 3.39369 | 0.01159 |
| K15737 | 3.64535 | 0.01022 |
| K15773 | 3.51408 | 0.02272 |
| K15827 | 3.80901 | 0.01998 |
| K15828 | 2.75543 | 0.02803 |
| K15831 | 4.83856 | 0.00531 |
| K15832 | 4.23103 | 0.00809 |
| K15833 | 3.33404 | 0.01438 |
| K15834 | 3.36063 | 0.01042 |
| K15836 | 3.75991 | 0.01021 |
| K15866 | 3.62735 | 0.01149 |
| K15977 | -2.4308 | 0.00686 |
| K15986 | 2.82139 | 0.0084 |
| K15987 | -4.4261 | 0.00352 |
| K16013 | 2.27232 | 0.0312 |
| K16074 | 3.67825 | 0.01436 |
| K16088 | 2.98857 | 0.0277 |
| K16090 | 2.8419 | 0.03997 |
| K16092 | 3.7296 | 0.01374 |
| K16138 | 3.71982 | 0.00765 |
| K16203 | 2.25591 | 0.04677 |
| K16212 | -2.1403 | 0.04459 |
| K16322 | 2.69971 | 0.02267 |
| K16328 | 3.66706 | 0.00686 |
| K16329 | 2.30906 | 0.04629 |

| | | | | | |
|---|---|---|---|---|---|
| K03660 | -2.3721 | 0.04603421 | K16345 | 2.58135 | 0.04174 |
| K03665 | -1.5991 | 0.03028191 | K16346 | 3.89294 | 0.00971 |
| K03666 | 4.89553 | 0.00232026 | K16370 | 3.40232 | 0.01262 |
| K03669 | 3.10035 | 0.0233207 | K16650 | 3.10017 | 0.02412 |
| K03674 | 4.45989 | 0.00629648 | K16693 | 3.37832 | 0.01231 |
| K03683 | 3.3231 | 0.0181366 | K16711 | 3.53389 | 0.02559 |
| K03684 | 4.19233 | 0.00609582 | K16789 | 3.37749 | 0.00543 |
| K03687 | 2.55304 | 0.01193397 | K16881 | 4.12534 | 0.00829 |
| K03690 | 2.49408 | 0.03151284 | K16898 | 2.30373 | 0.01628 |
| K03696 | 2.71885 | 0.0136338 | K16899 | 2.57826 | 0.011 |
| K03716 | 5.14874 | 0.00293278 | K16906 | 2.93602 | 0.04372 |
| K03719 | 3.42262 | 0.0159576 | K16923 | 5.49961 | 0.00235 |
| K03720 | 5.41739 | 0.00272423 | K16924 | 2.05126 | 0.03415 |
| K03722 | 2.47804 | 0.01768854 | K16925 | 3.72786 | 0.01727 |
| K03732 | 3.25063 | 0.01754712 | K16950 | 5.56714 | 0.00263 |
| K03733 | -2.891 | 0.01253288 | K17234 | -3.7714 | 0.01127 |
| K03735 | 2.40401 | 0.01674505 | K17236 | -4.2961 | 0.00854 |
| K03737 | -1.7128 | 0.02853389 | K17247 | 4.17339 | 0.00765 |
| K03745 | 5.76564 | 0.00262888 | K17319 | -2.0857 | 0.03891 |
| K03746 | 3.7858 | 0.01022454 | K17675 | -2.6836 | 0.01971 |
| K03748 | -1.7848 | 0.04910859 | K17680 | 3.02261 | 0.03263 |
| K03749 | 4.02773 | 0.00680859 | K17686 | 4.50497 | 0.00543 |
| K03752 | 3.51349 | 0.00530562 | K17713 | 2.68145 | 0.02205 |
| K03753 | 4.48882 | 0.00371482 | K17733 | 3.43572 | 0.01819 |
| K03755 | 3.48457 | 0.01754679 | K17734 | -3.0195 | 0.04682 |
| K03756 | 3.14498 | 0.00987284 | K17837 | 3.86047 | 0.01301 |
| K03758 | 5.53183 | 0.00232026 | K17865 | 2.67183 | 0.02114 |
| K03759 | 3.39071 | 0.01732737 | K17899 | 4.65198 | 0.00649 |
| K03760 | 2.55373 | 0.02779866 | K17948 | 3.29985 | 0.0277 |
| K03761 | 4.09746 | 0.00673221 | K18012 | 2.61901 | 0.02803 |
| K03762 | 4.46043 | 0.00679664 | K18013 | 2.86529 | 0.01364 |
| K03764 | 3.05314 | 0.02386871 | K18118 | 4.24652 | 0.00767 |
| K03767 | 4.06557 | 0.00824052 | K18122 | 4.08061 | 0.00801 |
| K03772 | 2.78125 | 0.02061651 | K18123 | 4.16645 | 0.00671 |
| K03776 | 4.04813 | 0.00854133 | K18138 | 3.58739 | 0.02318 |
| K03777 | 2.90759 | 0.0326321 | K18139 | 2.88141 | 0.0436 |
| K03779 | 4.02231 | 0.0082651 | K18140 | 4.27132 | 0.00824 |
| K03781 | -4.1035 | 0.00530562 | K18220 | -1.8074 | 0.00634 |
| K03783 | 2.4969 | 0.01417815 | K18344 | -4.3598 | 0.00986 |
| K03784 | 2.62955 | 0.01027704 | K18345 | -3.8019 | 0.01475 |
| K03785 | 3.46648 | 0.00597264 | K18350 | 3.27236 | 0.01374 |
| K03786 | 3.92714 | 0.0057887 | K18478 | 3.81791 | 0.01051 |
| K03789 | 3.0075 | 0.01404277 | K18581 | -3.1276 | 0.02534 |
| K03796 | 3.40362 | 0.01773767 | K18657 | 3.33866 | 0.01237 |
| K03800 | 2.95552 | 0.01178338 | K18692 | 3.21405 | 0.0277 |
| K03803 | 2.61683 | 0.01664521 | K18702 | 3.20205 | 0.01275 |
| K03807 | 2.48461 | 0.03310193 | K18765 | 3.40844 | 0.01789 |
| K03808 | 3.68609 | 0.01218982 | K18778 | 2.83792 | 0.01497 |

| | | | | | |
|---|---|---|---|---|---|
| K03814 | 4.93222 | 0.00358085 | K18815 | -2.4786 | 0.03634 |
| K03818 | -4.0472 | 0.00586344 | K18828 | 4.32935 | 0.00762 |
| K03820 | 2.85433 | 0.02474668 | K18829 | 3.34242 | 0.02999 |
| K03824 | 2.66633 | 0.04122202 | K18840 | 4.6638 | 0.00751 |
| K03826 | 2.06349 | 0.0359183 | K18856 | 3.08714 | 0.03117 |
| K03831 | 4.33738 | 0.00530562 | K18862 | 4.77772 | 0.0084 |
| K03833 | 5.58207 | 0.00232026 | K18866 | 2.60361 | 0.04115 |
| K03840 | 5.27531 | 0.00376385 | K18888 | -3.0282 | 0.01906 |
| K03841 | 2.7947 | 0.02061651 | K18889 | -2.8343 | 0.01922 |
| K03852 | -4.8995 | 0.00599652 | K18898 | 2.92758 | 0.03117 |
| K03855 | 3.7685 | 0.01016021 | K18919 | 6.64565 | 0.00371 |
| K03856 | 2.12065 | 0.04682172 | K18921 | 4.77157 | 0.0274 |
| K03892 | 2.19274 | 0.02419945 | K18922 | 6.46813 | 0.00382 |
| K03923 | 4.44584 | 0.00358085 | K18928 | 3.45514 | 0.0274 |
| K03969 | 3.61871 | 0.00530562 | K18979 | 4.95508 | 0.00352 |
| K03970 | 5.03812 | 0.0049058 | K19000 | 3.17512 | 0.01369 |
| K03971 | 5.00641 | 0.00466824 | K19055 | 5.12135 | 0.00379 |
| K03973 | 4.55046 | 0.00599652 | K19075 | 3.305 | 0.03637 |
| K04014 | 3.40823 | 0.00607733 | K19155 | 4.89245 | 0.0047 |
| K04021 | 3.20081 | 0.0212226 | K19159 | 2.54763 | 0.03292 |
| K04025 | 4.69374 | 0.00541922 | K19162 | 3.46401 | 0.02616 |
| K04028 | 2.48472 | 0.01344681 | K19165 | -5.0037 | 0.00754 |
| K04030 | 3.49051 | 0.0084524 | K19168 | 5.23921 | 0.00392 |
| K04031 | 2.82753 | 0.01369976 | K19169 | -5.5833 | 0.00394 |
| K04034 | 5.49742 | 0.00272423 | K19170 | -6.3155 | 0.00232 |
| K04042 | 2.24796 | 0.01242165 | K19171 | -3.9213 | 0.0087 |
| K04043 | -0.8859 | 0.04682172 | K19172 | -6.4384 | 0.00232 |
| K04062 | 2.90305 | 0.04758469 | K19221 | 3.91189 | 0.01403 |
| K04070 | 3.85608 | 0.00599652 | K19222 | 4.0639 | 0.01059 |
| K04073 | 4.31138 | 0.00483735 | K19228 | 4.33264 | 0.00631 |
| K04075 | 3.38036 | 0.00530562 | K19230 | 5.12621 | 0.00423 |
| K04076 | 5.66058 | 0.00232026 | K19236 | 3.2343 | 0.02303 |
| K04077 | -0.986 | 0.03570058 | K19265 | 4.7233 | 0.00543 |
| K04080 | 2.96462 | 0.02411766 | K19267 | 3.9583 | 0.00805 |
| K04081 | 3.12773 | 0.01301388 | K19270 | 3.38172 | 0.01324 |
| K04082 | 3.19388 | 0.0137402 | K19294 | 2.18383 | 0.04838 |
| K04083 | 1.87983 | 0.02966435 | K19303 | 4.18024 | 0.00875 |
| K04088 | 3.47867 | 0.01073472 | K19334 | 4.02424 | 0.01052 |
| K04335 | 4.03394 | 0.01160662 | K19335 | 3.60679 | 0.01667 |
| K04337 | 2.65708 | 0.03288099 | K19336 | 4.53367 | 0.00592 |
| K04485 | 2.26938 | 0.0222117 | K19350 | 2.25501 | 0.04454 |
| K04516 | -1.6567 | 0.04135602 | K19354 | 4.03158 | 0.00644 |
| K04517 | 2.51875 | 0.02600768 | K19405 | 5.58596 | 0.0025 |
| K04565 | 3.61346 | 0.00929169 | K19411 | 4.30785 | 0.00613 |
| K04651 | 3.66605 | 0.00599652 | K19419 | 2.70234 | 0.03925 |
| K04653 | 4.39318 | 0.00483735 | K19431 | 3.07907 | 0.04197 |
| K04654 | 3.16869 | 0.0151405 | K19577 | 2.66691 | 0.04282 |
| K04691 | 3.99597 | 0.00761685 | K19591 | 2.85385 | 0.03133 |

| | | | | | |
|---|---|---|---|---|---|
| K04744 | 2.73571 | 0.02317096 | K19611 | 4.38467 | 0.00602 |
| K04750 | 3.14727 | 0.01037298 | K19688 | 3.5129 | 0.01428 |
| K04752 | 4.29509 | 0.0078317 | K19776 | 3.5672 | 0.00868 |
| K04754 | 4.74305 | 0.00530562 | K19777 | 4.97641 | 0.00616 |
| K04755 | 2.72904 | 0.02556247 | K19778 | 4.21029 | 0.00992 |
| K04759 | 2.3341 | 0.01054093 | K19784 | 2.55838 | 0.04305 |
| K04770 | 2.83015 | 0.0301977 | K19789 | 4.02553 | 0.00854 |
| K04771 | 2.93145 | 0.00761685 | K19802 | 2.62595 | 0.03332 |
| K04774 | 3.29524 | 0.01996671 | K19804 | 3.84649 | 0.00586 |
| K04775 | 2.7629 | 0.04984752 | K19955 | 2.58252 | 0.01369 |
| K04835 | -3.1849 | 0.01979006 | K19956 | -2.6634 | 0.01997 |
| K05245 | 3.35918 | 0.01537651 | K20107 | 4.52814 | 0.00531 |
| K05337 | 3.10577 | 0.00867507 | K20108 | 4.52814 | 0.00531 |
| K05341 | -3.2682 | 0.01902299 | K20265 | 4.18627 | 0.00488 |
| K05365 | 2.55321 | 0.04870107 | K20542 | 2.60903 | 0.02108 |
| K05396 | 2.9273 | 0.02998692 | K20881 | 2.76858 | 0.02469 |
| K05499 | 3.01404 | 0.02802893 | K20885 | 3.54744 | 0.01675 |
| K05501 | 3.29888 | 0.00910465 | K20904 | 2.8936 | 0.01658 |
| K05516 | 2.82498 | 0.02644688 | K21029 | 4.34691 | 0.00488 |
| K05526 | 4.22866 | 0.00761685 | K21065 | 3.47491 | 0.01322 |
| K05527 | 4.23572 | 0.00711873 | K21085 | 2.71741 | 0.04139 |
| K05539 | 2.80681 | 0.02190753 | K21086 | 2.77148 | 0.0309 |
| K05540 | 2.71487 | 0.01314796 | K21088 | 4.07718 | 0.00702 |
| K05591 | 3.73294 | 0.01095206 | K21394 | 4.03344 | 0.00697 |
| K05594 | 3.83276 | 0.01519593 | K21395 | 4.63152 | 0.00531 |
| K05595 | 2.51584 | 0.02853389 | K21399 | 4.68309 | 0.01054 |
| K05596 | 4.26716 | 0.00644444 | K21405 | 4.01785 | 0.01079 |
| K05709 | 4.34722 | 0.007236 | K21469 | 3.50458 | 0.02402 |
| K05710 | 4.18652 | 0.00992189 | K21514 | -2.9198 | 0.01658 |
| K05713 | 2.59538 | 0.04918977 | K21637 | 4.51505 | 0.00883 |
| K05714 | 3.56485 | 0.01475665 | K21638 | 4.18434 | 0.01037 |
| K05775 | 4.10907 | 0.00586344 | K21695 | 6.39739 | 0.00232 |
| K05776 | 2.82396 | 0.03600819 | K21742 | 2.59021 | 0.02635 |
| K05777 | 2.49793 | 0.03958926 | K21759 | 4.10842 | 0.00854 |
| K05779 | 5.01316 | 0.00530562 | K21901 | 2.53164 | 0.03916 |
| K05782 | 3.56328 | 0.01410262 | K21908 | 3.6571 | 0.0158 |
| K05785 | 3.82222 | 0.00854133 | K21929 | -1.9189 | 0.02986 |
| K05786 | 3.95481 | 0.00580391 | K21965 | 3.22131 | 0.01464 |
| K05787 | 4.30053 | 0.00696579 | K21975 | 3.73672 | 0.01879 |
| K05788 | 4.6656 | 0.00530562 | K21976 | 4.40226 | 0.00868 |
| K05798 | 2.75929 | 0.04574482 | K21977 | -5.9858 | 0.00313 |
| K05799 | 4.16854 | 0.00381623 | K21990 | 2.81885 | 0.03925 |
| K05803 | 2.70069 | 0.03117343 | K21993 | 2.30911 | 0.03447 |
| K05804 | 4.1874 | 0.00670795 | K22015 | 4.64896 | 0.00634 |
| K05805 | 2.66865 | 0.02258846 | K22041 | 3.75417 | 0.01105 |
| K05807 | 2.56153 | 0.03399376 | K22044 | 3.91077 | 0.00406 |
| K05809 | 2.92943 | 0.0429125 | K22110 | 3.52202 | 0.00986 |
| K05810 | 2.58098 | 0.01418801 | K22111 | 4.01073 | 0.0084 |

| | | |
|---|---|---|
| K05812 | 4.49263 | 0.00609582 |
| K05813 | 2.51417 | 0.02815799 |
| K05814 | 4.32603 | 0.00613025 |
| K05815 | 3.7307 | 0.00864767 |
| K05816 | 3.73385 | 0.0104469 |
| K05818 | 3.11604 | 0.01879564 |
| K05822 | 3.13746 | 0.03044527 |
| K05845 | 2.41923 | 0.03437897 |
| K22468 | 3.14738 | 0.03762015 |
| K22718 | 5.52892 | 0.00761685 |
| K22719 | 2.91726 | 0.02317096 |

| | | |
|---|---|---|
| K22130 | 4.39616 | 0.0061 |
| K22132 | 2.35344 | 0.02966 |
| K22214 | 2.59607 | 0.03891 |
| K22227 | 2.74176 | 0.03755 |
| K22304 | 4.98221 | 0.00649 |
| K22373 | 2.59257 | 0.03916 |
| K22443 | 2.55178 | 0.04372 |
| K22444 | 3.10062 | 0.02663 |
| K23007 | 3.22471 | 0.01454 |
| K23015 | 3.06317 | 0.0325 |

**Supplementary Table 3.** Statistically significant KOs mapped to KEGG pathways. Pathways are shown in decreasing order of mapped KOs. Only pathways containing between 50-20 KOs were considered for subsequent analysis to exclude the most general and the least abundant ones. KO numbers and symbols have been included for each gene.

| KEGG pathways | Number of mapped KOs | Mapped KOs |
|---|---|---|
| Biosynthesis of aminoacids | 47 | K00030 IDH3,K00053 ilvC,K00265 gltB,K00266 gltD,K00290 LYS1,K00548 metH,K00620 argJ,K00640 cysE,K00641 metX,K00674 dapD,K00789 metK,K00817 hisC,K00832 tyrB,K00930 argB,K00931 proB,K01609 trpC,K01682 acnB,K01687 ilvD,K01693 hisB,K01695 trpA,K01703 leuC,K01738 cysK,K01760 metC,K01783 rpe,K01958 PC,K02500 hisF,K03785 aroD,K03786 aroQ,K03856 AROA2,K04516 AROA1,K04517 tyrA2,K05822 dapH,K05942 E2.3.3.3,K06001 trpB,K06209 pheB,K07173 luxS,K09758 asdA,K12524 thrA,K13497 trpGD,K13498 trpCF,K14155 patB,K14170 pheA,K14187 tyrA,K14682 argAB,K15633 gpmI,K15634 gpmB,K16370 pfkB |
| Pyruvate metabolism | 35 | K00101 lldD,K00116 mqo,K00121 frmA,K00138 aldB,K00161 PDHA,K00170 porB,K00175 korB,K00244 frdA,K00245 frdB,K00247 frdD,K00382 DLD,K00625 pta,K00626 ACAT,K00627 DLAT,K00656 E2.3.1.54,K01512 acyP,K01571 oadA,K01595 ppc,K01679 E4.2.1.2B,K01958 PC,K01962 accA,K01963 accD,K02160 accB,K03737 por,K03777 dld,K04021 eutE,K04073 mhpF,K07248 aldA,K12957 ahr,K12972 ghrA,K13788 pta,K13953 adhP,K13954 yiaY,K18118 aarC,K22373 larA |
| Purine metabolism | 29 | K00073 allD,K00087 ygeS,K00526 E1.17.4.1B,K00769 gpt,K00926 arcC,K00957 cysD,K01119 cpdB,K01139 spoT,K01466 allB,K01486 ade,K01488 add,K01525 apaH,K01923 purC,K02083 allC,K02566 nagD,K03651 cpdA,K03783 punA,K03784 deoD,K05810 LACC1,K05873 cyaB,K06966 ppnN,K07127 uraH,K08722 yfbR,K09913 ppnP,K11175 purN,K11178 yagS,K11751 ushA,K13483 yagT,K20881 yrfG |
| Quorum sensing | 27 | K01497 ribA,K01897 ACSL,K01999 livK,K02032 ddpF,K02034 ABC.PE.P1,K02035 ABC.PE.S,K02055 ABC.SP.S,K02402 flhC,K03071 secB,K03076 secY,K03666 hfq,K07173 luxS,K07666 qseB,K07667 kdpE,K07711 glrK,K07715 glrR,K07782 sdiA,K07813 agrB,K10555 lsrB,K10558 lsrA,K10914 crp,K11752 ribD,K15580 oppA,K15581 oppB,K15583 oppD,K18139 oprM,K20265 gadC |
| Butanoate metabolism | 24 | K00170 porB,K00175 korB,K00242 sdhD,K00244 frdA,K00245 frdB,K00247 frdD,K00248 ACADS,K00626 ACAT,K00634 ptb,K00656 E2.3.1.54,K00823 puuE,K01028 scoA,K01034 atoD,K01035 atoA,K01040 gctB,K01640 HMGCL,K01641 HMGCS |

| | | |
|---|---|---|
| | | ,K01715 crt,K01782 fadJ,K03737 por,K04073 mhpF,K17865 croR,K18118 aarC,K18122 cat2 |
| Amino sugar and nucleotide metabolism | 24 | K00075 murB,K00844 HK,K00847 E2.7.1.4,K00884 NAGK,K00885 nanK,K00978 rfbF,K01443 nagA,K01639 E4.1.3.3,K01709 rfbG,K01787 RENBP,K02472 wecC,K02777 crr,K02796 manZ,K04042 glmU,K06859 pgi1,K07806 arnB,K08678 UXS1,K09001 anmK,K10011 arnA,K10012 arnC,K11192 murP,K12452 ascC,K13014 arnD,K16881 EC:2.7.7.13 5.4.2.8 |
| Nucleotide metabolism | 23 | K00087 ygeS,K00526 E1.17.4.1B,K00757 udp,K00769 gpt,K01250 rihA,K01486 ade,K01488 add,K01494 dcd,K02566 nagD,K03465 thyX,K03783 punA,K03784 deoD,K05810 LACC1,K06966 ppnN,K07043 upp,K08320 nudG,K08722 yfbR,K08723 yjjG,K09913 ppnP,K11178 yagS,K11751 ushA,K13483 yagT,K20881 yrfG |
| Propanoate metabolism | 23 | K00005 gldA,K00170 porB,K00248 ACADS,K00382 DLD,K00625 pta,K00656 E2.3.1.54,K00823 puuE,K00932 tdcD,K01659 prpC,K01682 acnB,K01720 prpD,K01782 fadJ,K01903 sucC,K01908 ACSS3,K01962 accA,K01963 accD,K02160 accB,K11264 scpB,K13788 pta,K13919 pduD,K13920 pduE,K13922 pduP,K22214 scpC |
| Glyoxylate and dicarboxylate metabolism | 22 | K00042 garR,K00104 glcD,K00123 fdoG,K00124 fdoH,K00127 fdoI,K00281 GLDC,K00282 gcvPA,K00382 DLD,K00626 ACAT,K00865 glxK,K01577 oxc,K01682 acnB,K01816 hyi,K03779 ttdA,K03781 katE,K04835 mal,K07248 aldA,K11472 glcE,K11473 glcF,K12972 ghrA,K17865 croR,K18123 HOGA1 |
| Glycolysis / gluconeogenesis | 22 | K00121 frmA,K00131 gapN,K00138 aldB,K00161 PDHA,K00170 porB,K00175 korB,K00382 DLD,K00627 DLAT,K00844 HK,K01085 agp,K01792 E5.1.3.15,K02446 glpX,K02777 crr,K03737 por,K03841 FBP,K06859 pgi1,K12957 ahr,K13953 adhP,K13954 yiaY,K15633 gpmI,K15634 gpmB,K16370 pfkB |
| Pyrimidine metabolism | 21 | K00254 DHODH,K00526 E1.17.4.1B,K00757 udp,K01119 cpdB,K01250 rihA,K01494 dcd,K02566 nagD,K02823 pyrDII,K03465 thyX,K06966 ppnN,K07043 upp,K08320 nudG,K08722 yfbR,K08723 yjjG,K09018 rutA,K09020 rutB,K09023 rutD,K09913 ppnP,K11751 ushA,K16328 psuK,K16329 psuG |
| Ribosome | 21 | K02867 RP-L11,K02874 RP-L14,K02876 RP-L15,K02878 RP-L16,K02879 RP-L17,K02887 RP-L20,K02890 RP-L22,K02892 RP-L23,K02897 RP-L25,K02899 RP-L27,K02919 RP-L36,K02933 RP-L6,K02946 RP-S10,K02948 RP-S11,K02952 RP-S13,K02959 RP-S16,K02965 RP-S19,K02970 RP-S21,K02982 RP-S3,K02992 RP-S7,K07590 RP-L7A |
| Carbon fixation | 21 | K00170 porB,K00175 korB,K00198 cooS,K00242 sdhD,K00244 frdA,K00245 frdB,K00247 frdD,K00625 pta,K00626 ACAT,K01491 folD,K01595 ppc,K01679 E4.2.1.2B,K01682 acnB,K0 |

| | | |
|---|---|---|
| pathways in prokaryotes | | 1903 sucC,K01958 PC,K01962 accA,K01963 accD,K02160 accB,K03737 por,K13788 pta,K22015 fdhF |
| Sulfur metabolism | 21 | K00380 cysJ,K00381 cysI,K00385 asrC,K00394 aprA,K00640 cysE,K00641 metX,K00957 cysD,K01082 cysQ,K01738 cysK,K02045 cysA,K02046 cysU,K02047 cysW,K02439 glpE,K03119 tauD,K06881 nrnA,K07306 dmsA,K07307 dmsB,K15553 ssuA,K15554 ssuC,K15555 ssuB,K16950 asrA |
| Oxidative phosphorylation | 21 | K00242 sdhD,K00244 frdA,K00245 frdB,K00247 frdD,K00331 nuoB,K00333 nuoD,K00334 nuoE,K00338 nuoI,K00340 nuoK,K00425 cydA,K00937 ppk1,K02107 ATPVG,K02110 ATPF0C,K02122 ATPVF,K02257 COX10,K02297 cyoA,K02299 cyoC,K02300 cyoD,K06019 ppaX,K15986 ppaC,K22468 ppk2 |

**Supplementary Table 4**. Significantly deregulated GO terms between the anaemic and control groups. GO IDs, category within the GO database, p-adjusted values, normalized enrichment scores (NES) and number of genes contributing to deregulation (leading edge genes) are included for each GO term

| GO ID | Category | padj | NES | size |
|---|---|---|---|---|
| GO:0051966 | Biological process | 1.64E-05 | -1.7409691 | 44 |
| GO:0007269 | Biological process | 7.16E-09 | -1.73741838 | 131 |
| GO:0007416 | Biological process | 7.16E-09 | -1.73690509 | 123 |
| GO:0086002 | Biological process | 9.19E-05 | -1.73631829 | 39 |
| GO:0051965 | Biological process | 2.10E-05 | -1.73577385 | 44 |
| GO:0035637 | Biological process | 7.16E-09 | -1.73082969 | 149 |
| GO:0060078 | Biological process | 1.16E-07 | -1.73021452 | 92 |
| GO:0099174 | Biological process | 0.00041623 | -1.72969854 | 22 |
| GO:0086003 | Biological process | 1.72E-05 | -1.72206483 | 55 |
| GO:0097553 | Biological process | 2.75E-08 | -1.72148186 | 108 |
| GO:0061337 | Biological process | 1.70E-08 | -1.71668855 | 111 |
| GO:0035249 | Biological process | 1.39E-05 | -1.71420965 | 58 |
| GO:0006836 | Biological process | 7.16E-09 | -1.7139718 | 166 |
| GO:0001508 | Biological process | 5.66E-08 | -1.71373466 | 104 |
| GO:0062237 | Biological process | 0.00027127 | -1.71183561 | 37 |
| GO:0051279 | Biological process | 6.81E-06 | -1.71063506 | 65 |
| GO:0050432 | Biological process | 0.00014334 | -1.70802244 | 40 |
| GO:0051963 | Biological process | 6.08E-06 | -1.70769923 | 76 |
| GO:0051588 | Biological process | 1.47E-06 | -1.70669418 | 89 |
| GO:1901019 | Biological process | 7.78E-06 | -1.70447316 | 72 |
| GO:0099565 | Biological process | 4.48E-06 | -1.70153632 | 81 |
| GO:0099560 | Biological process | 0.00114395 | -1.6999854 | 19 |
| GO:0086010 | Biological process | 0.00062751 | -1.69913588 | 30 |
| GO:0019226 | Biological process | 6.30E-05 | -1.69884007 | 47 |
| GO:0015837 | Biological process | 1.47E-05 | -1.69756628 | 67 |
| GO:0099536 | Biological process | 7.16E-09 | -1.68909723 | 531 |
| GO:1903169 | Biological process | 1.04E-07 | -1.68702192 | 116 |
| GO:0060401 | Biological process | 7.16E-09 | -1.68643044 | 141 |
| GO:0001505 | Biological process | 7.16E-09 | -1.68611058 | 175 |
| GO:0086026 | Biological process | 0.00275934 | -1.68072358 | 15 |
| GO:1903779 | Biological process | 0.00010719 | -1.67875805 | 60 |
| GO:0099172 | Biological process | 0.00060394 | -1.67865005 | 35 |
| GO:0016079 | Biological process | 2.58E-06 | -1.67804305 | 90 |
| GO:0050808 | Biological process | 7.16E-09 | -1.67793032 | 326 |
| GO:2001257 | Biological process | 2.81E-08 | -1.67703564 | 130 |
| GO:0015844 | Biological process | 0.00018278 | -1.67619778 | 59 |
| GO:0099625 | Biological process | 0.00353985 | -1.67563363 | 20 |
| GO:0050806 | Biological process | 1.70E-08 | -1.67194222 | 129 |
| GO:2000300 | Biological process | 0.00021116 | -1.67167274 | 59 |

| | | | | |
|---|---|---|---|---|
| GO:0099623 | Biological process | 0.00274619 | -1.67134794 | 19 |
| GO:0007612 | Biological process | 4.78E-07 | -1.6687611 | 112 |
| GO:0098693 | Biological process | 9.95E-06 | -1.66811348 | 85 |
| GO:0050905 | Biological process | 0.00014048 | -1.66685568 | 69 |
| GO:0070252 | Biological process | 1.03E-05 | -1.66652374 | 86 |
| GO:0030199 | Biological process | 0.00050084 | -1.66627781 | 42 |
| GO:0010522 | Biological process | 3.26E-05 | -1.66538681 | 81 |
| GO:0043266 | Biological process | 0.00015214 | -1.66441716 | 69 |
| GO:0061037 | Biological process | 0.00275934 | -1.66285797 | 24 |
| GO:0098815 | Biological process | 0.00153374 | -1.66279797 | 37 |
| GO:0099622 | Biological process | 0.00274619 | -1.6627623 | 27 |
| GO:0042391 | Biological process | 7.16E-09 | -1.6608888 | 311 |
| GO:0060048 | Biological process | 2.76E-06 | -1.66066286 | 101 |
| GO:0007218 | Biological process | 0.0017039 | -1.66033501 | 39 |
| GO:0070050 | Biological process | 0.00289258 | -1.65952738 | 27 |
| GO:0003015 | Biological process | 7.16E-09 | -1.6591499 | 219 |
| GO:0099177 | Biological process | 7.16E-09 | -1.65843834 | 346 |
| GO:0035418 | Biological process | 8.09E-05 | -1.65764266 | 64 |
| GO:0086019 | Biological process | 0.00405396 | -1.6554423 | 23 |
| GO:0086065 | Biological process | 0.00099542 | -1.65466851 | 43 |
| GO:0070839 | Biological process | 0.00289028 | -1.65439143 | 10 |
| GO:0014059 | Biological process | 0.00405396 | -1.65384418 | 23 |
| GO:0086001 | Biological process | 0.00034283 | -1.65292274 | 56 |
| GO:0050654 | Biological process | 0.00268999 | -1.64762883 | 36 |
| GO:0006813 | Biological process | 1.25E-08 | -1.64451935 | 156 |
| GO:0099003 | Biological process | 7.16E-09 | -1.64269336 | 166 |
| GO:1904427 | Biological process | 0.00047481 | -1.64236848 | 53 |
| GO:0086009 | Biological process | 0.00191428 | -1.64138154 | 35 |
| GO:0042417 | Biological process | 0.00581026 | -1.63952246 | 24 |
| GO:0060314 | Biological process | 0.00309832 | -1.6384807 | 25 |
| GO:0051926 | Biological process | 0.00143213 | -1.63831605 | 42 |
| GO:0070296 | Biological process | 0.00419114 | -1.63783391 | 29 |
| GO:1903514 | Biological process | 0.00435001 | -1.63735069 | 26 |
| GO:0006941 | Biological process | 9.10E-07 | -1.63694711 | 125 |
| GO:0097120 | Biological process | 0.00145114 | -1.63690703 | 42 |
| GO:1901890 | Biological process | 7.76E-05 | -1.6366132 | 75 |
| GO:0050803 | Biological process | 7.16E-09 | -1.63598381 | 182 |
| GO:0015872 | Biological process | 0.00336714 | -1.63577707 | 31 |
| GO:0051953 | Biological process | 0.00573814 | -1.63236063 | 18 |
| GO:0051281 | Biological process | 0.00375289 | -1.63204679 | 31 |
| GO:0031644 | Biological process | 4.27E-06 | -1.63198911 | 116 |
| GO:0060306 | Biological process | 0.00407879 | -1.63047526 | 25 |
| GO:1903522 | Biological process | 7.16E-09 | -1.62963975 | 221 |
| GO:0007613 | Biological process | 2.10E-05 | -1.62897901 | 95 |
| GO:0007205 | Biological process | 0.009281 | -1.62881476 | 22 |
| GO:0072132 | Biological process | 0.00191428 | -1.62754724 | 42 |
| GO:0006936 | Biological process | 7.16E-09 | -1.62544565 | 263 |
| GO:0086012 | Biological process | 0.01288059 | -1.62255296 | 20 |

| | | | | |
|---|---|---|---|---|
| GO:0032331 | Biological process | 0.00977624 | -1.62224006 | 19 |
| GO:1904062 | Biological process | 7.16E-09 | -1.62167711 | 259 |
| GO:0051208 | Biological process | 4.46E-05 | -1.62116928 | 98 |
| GO:0099084 | Biological process | 0.00442933 | -1.61898106 | 28 |
| GO:0051899 | Biological process | 0.00040056 | -1.61885208 | 74 |
| GO:0099175 | Biological process | 8.37E-05 | -1.61869132 | 87 |
| GO:0098900 | Biological process | 0.00241079 | -1.61807356 | 44 |
| GO:0070509 | Biological process | 0.00050296 | -1.61668247 | 64 |
| GO:0050885 | Biological process | 0.00509103 | -1.6166647 | 37 |
| GO:0043113 | Biological process | 0.00218834 | -1.61649893 | 49 |
| GO:0031115 | Biological process | 0.01100156 | -1.6147378 | 13 |
| GO:1905874 | Biological process | 0.01024076 | -1.61254328 | 12 |
| GO:0001755 | Biological process | 0.0018527 | -1.61188115 | 48 |
| GO:1902414 | Biological process | 0.00010313 | -1.6109143 | 84 |
| GO:0010524 | Biological process | 0.00448694 | -1.61076962 | 43 |
| GO:0034765 | Biological process | 7.16E-09 | -1.60611913 | 352 |
| GO:0070588 | Biological process | 7.16E-09 | -1.60516265 | 235 |
| GO:0035235 | Biological process | 0.02215798 | -1.60499212 | 15 |
| GO:0099637 | Biological process | 0.00895528 | -1.60475308 | 36 |
| GO:0051480 | Biological process | 7.16E-09 | -1.60452229 | 249 |
| GO:0021795 | Biological process | 0.00733228 | -1.60449834 | 31 |
| GO:0007156 | Biological process | 0.0003002 | -1.60399233 | 75 |
| GO:0043271 | Biological process | 3.77E-05 | -1.60358311 | 107 |
| GO:0034767 | Biological process | 7.14E-06 | -1.6035033 | 122 |
| GO:1903170 | Biological process | 0.00890021 | -1.60253447 | 30 |
| GO:0001941 | Biological process | 0.00755336 | -1.60157776 | 32 |
| GO:0098801 | Biological process | 0.01356916 | -1.6009436 | 27 |
| GO:0033555 | Biological process | 0.00200872 | -1.59887595 | 56 |
| GO:0090102 | Biological process | 0.00588382 | -1.59437104 | 35 |
| GO:0014047 | Biological process | 0.01160874 | -1.59398846 | 34 |
| GO:0021695 | Biological process | 0.00709992 | -1.59366465 | 41 |
| GO:0042596 | Biological process | 0.01024076 | -1.59266 | 26 |
| GO:0055078 | Biological process | 0.01211344 | -1.59193466 | 36 |
| GO:0060291 | Biological process | 0.00099542 | -1.59125012 | 65 |
| GO:0097484 | Biological process | 0.01320475 | -1.5909312 | 29 |
| GO:0006942 | Biological process | 0.00057576 | -1.59064224 | 70 |
| GO:0010959 | Biological process | 7.16E-09 | -1.59006661 | 289 |
| GO:0035812 | Biological process | 0.0198467 | -1.58870165 | 14 |
| GO:2001259 | Biological process | 0.00241079 | -1.58603534 | 57 |
| GO:0010881 | Biological process | 0.02436529 | -1.5855713 | 19 |
| GO:0021696 | Biological process | 0.01883392 | -1.58554765 | 27 |
| GO:0098742 | Biological process | 2.42E-06 | -1.58450588 | 151 |
| GO:0008038 | Biological process | 0.00869804 | -1.58399479 | 38 |
| GO:0021575 | Biological process | 0.00735211 | -1.58369598 | 35 |
| GO:0060411 | Biological process | 0.00326757 | -1.58327763 | 60 |
| GO:0010644 | Biological process | 0.01329582 | -1.58262919 | 25 |
| GO:0061035 | Biological process | 0.00310261 | -1.58136101 | 56 |
| GO:0007215 | Biological process | 0.00188222 | -1.58125139 | 61 |

| | | | | |
|---|---|---|---|---|
| GO:0048167 | Biological process | 2.69E-06 | -1.58069299 | 147 |
| GO:0007638 | Biological process | 0.01762553 | -1.58056257 | 11 |
| GO:0035640 | Biological process | 0.02298407 | -1.57913548 | 18 |
| GO:0050805 | Biological process | 0.00392796 | -1.57663618 | 55 |
| GO:1901888 | Biological process | 1.39E-06 | -1.57649632 | 161 |
| GO:0099010 | Biological process | 0.02543047 | -1.57560967 | 14 |
| GO:0033604 | Biological process | 0.02435814 | -1.5754108 | 10 |
| GO:0051924 | Biological process | 1.58E-07 | -1.57537825 | 186 |
| GO:0099173 | Biological process | 9.44E-06 | -1.57524552 | 143 |
| GO:0032330 | Biological process | 0.01159065 | -1.57501271 | 40 |
| GO:0051589 | Biological process | 0.02298407 | -1.57478275 | 13 |
| GO:0007157 | Biological process | 0.01469293 | -1.57446075 | 32 |
| GO:0051968 | Biological process | 0.03105429 | -1.57444523 | 20 |
| GO:1905314 | Biological process | 0.01469293 | -1.57430854 | 32 |
| GO:0048566 | Biological process | 0.01318265 | -1.57347237 | 28 |
| GO:0030048 | Biological process | 7.89E-05 | -1.57232315 | 112 |
| GO:0022037 | Biological process | 0.00047467 | -1.57125025 | 91 |
| GO:0048546 | Biological process | 0.0097107 | -1.56967372 | 42 |
| GO:0007620 | Biological process | 0.02824282 | -1.56946381 | 14 |
| GO:0007626 | Biological process | 1.03E-05 | -1.56934902 | 146 |
| GO:0099150 | Biological process | 0.02872097 | -1.56928861 | 10 |
| GO:1902656 | Biological process | 0.02436529 | -1.56916937 | 17 |
| GO:0099587 | Biological process | 0.00371838 | -1.56885505 | 69 |
| GO:0014033 | Biological process | 0.00253551 | -1.56881472 | 74 |
| GO:0032409 | Biological process | 4.37E-08 | -1.56868048 | 215 |
| GO:0001964 | Biological process | 0.02406634 | -1.56820755 | 12 |
| GO:1904861 | Biological process | 0.03588006 | -1.56793631 | 20 |
| GO:0006937 | Biological process | 4.84E-05 | -1.5673719 | 123 |
| GO:0099563 | Biological process | 0.03719504 | -1.56632166 | 20 |
| GO:0035641 | Biological process | 0.02441186 | -1.56596322 | 12 |
| GO:0043269 | Biological process | 7.16E-09 | -1.56595343 | 503 |
| GO:0097091 | Biological process | 0.02455846 | -1.56543007 | 12 |
| GO:0003177 | Biological process | 0.03366306 | -1.5638302 | 19 |
| GO:0008090 | Biological process | 0.03021053 | -1.56296954 | 18 |
| GO:0008356 | Biological process | 0.04872042 | -1.56160977 | 15 |
| GO:0003197 | Biological process | 0.02497148 | -1.56021073 | 36 |
| GO:0098698 | Biological process | 0.03656552 | -1.56003616 | 16 |
| GO:0060412 | Biological process | 0.02600597 | -1.55969204 | 36 |
| GO:0019935 | Biological process | 8.30E-06 | -1.55932487 | 152 |
| GO:0031111 | Biological process | 0.01320475 | -1.55886008 | 37 |
| GO:0003170 | Biological process | 0.00509103 | -1.55654365 | 53 |
| GO:0014855 | Biological process | 0.00826414 | -1.55641097 | 50 |
| GO:0042476 | Biological process | 0.00133074 | -1.55442948 | 89 |
| GO:0002027 | Biological process | 0.00384986 | -1.55438233 | 74 |
| GO:0006029 | Biological process | 0.00509103 | -1.55389358 | 68 |
| GO:0099509 | Biological process | 0.04097282 | -1.55334667 | 16 |
| GO:0002026 | Biological process | 0.04266471 | -1.55233651 | 20 |
| GO:0030206 | Biological process | 0.04895488 | -1.55114803 | 22 |

| | | | | |
|---|---|---|---|---|
| GO:0060021 | Biological process | 0.00551245 | -1.54989737 | 67 |
| GO:0097205 | Biological process | 0.03276376 | -1.5498729 | 21 |
| GO:0050650 | Biological process | 0.02209981 | -1.54978102 | 26 |
| GO:0050877 | Biological process | 7.16E-09 | -1.54937476 | 649 |
| GO:0010919 | Biological process | 0.03253904 | -1.54921251 | 12 |
| GO:0006939 | Biological process | 0.00186865 | -1.54911059 | 82 |
| GO:0007271 | Biological process | 0.03813286 | -1.54888391 | 24 |
| GO:0086091 | Biological process | 0.02872097 | -1.54862939 | 29 |
| GO:0051928 | Biological process | 0.0015748 | -1.54777985 | 87 |
| GO:0098810 | Biological process | 0.04223084 | -1.54758855 | 19 |
| GO:0003012 | Biological process | 7.16E-09 | -1.54752923 | 333 |
| GO:0071340 | Biological process | 0.03125941 | -1.54724753 | 11 |
| GO:0048484 | Biological process | 0.03199651 | -1.54715325 | 11 |
| GO:0007618 | Biological process | 0.0198467 | -1.54520231 | 28 |
| GO:0034762 | Biological process | 7.16E-09 | -1.54508656 | 419 |
| GO:0051149 | Biological process | 0.00633311 | -1.54480934 | 68 |
| GO:0099072 | Biological process | 0.0065883 | -1.54439397 | 53 |
| GO:2000826 | Biological process | 0.02004622 | -1.54311046 | 28 |
| GO:0051668 | Biological process | 0.00241079 | -1.54221616 | 82 |
| GO:0007610 | Biological process | 7.16E-09 | -1.54100746 | 444 |
| GO:0030902 | Biological process | 0.0002301 | -1.5405882 | 126 |
| GO:0072507 | Biological process | 7.16E-09 | -1.54056208 | 358 |
| GO:1903115 | Biological process | 0.03203993 | -1.54037033 | 33 |
| GO:0044057 | Biological process | 7.16E-09 | -1.5402216 | 450 |
| GO:0007193 | Biological process | 0.01169613 | -1.54020754 | 50 |
| GO:0097479 | Biological process | 0.01244184 | -1.53995649 | 48 |
| GO:0055117 | Biological process | 0.00603655 | -1.53900041 | 62 |
| GO:0007187 | Biological process | 2.36E-05 | -1.53882873 | 148 |
| GO:0021533 | Biological process | 0.04988816 | -1.53785652 | 16 |
| GO:0002028 | Biological process | 0.00588382 | -1.53732799 | 72 |
| GO:0050890 | Biological process | 2.27E-07 | -1.53603593 | 230 |
| GO:0051703 | Biological process | 0.02401121 | -1.53511788 | 35 |
| GO:0030534 | Biological process | 0.00170233 | -1.53487908 | 93 |
| GO:0061005 | Biological process | 0.02600597 | -1.53455838 | 43 |
| GO:0021681 | Biological process | 0.0455696 | -1.53430388 | 10 |
| GO:0050795 | Biological process | 0.02051488 | -1.534255 | 42 |
| GO:1901862 | Biological process | 0.02600597 | -1.53425186 | 43 |
| GO:0043583 | Biological process | 3.44E-05 | -1.53406165 | 157 |
| GO:0010523 | Biological process | 0.04704893 | -1.53400902 | 13 |
| GO:0046839 | Biological process | 0.02524596 | -1.53352552 | 40 |
| GO:0061351 | Biological process | 0.00062751 | -1.53348935 | 108 |
| GO:0034330 | Biological process | 7.16E-09 | -1.53231978 | 560 |
| GO:0003203 | Biological process | 0.02303474 | -1.53165661 | 28 |
| GO:0021819 | Biological process | 0.04842179 | -1.53162645 | 10 |
| GO:0051932 | Biological process | 0.04328162 | -1.53061406 | 30 |
| GO:0090090 | Biological process | 3.44E-05 | -1.53029575 | 153 |
| GO:0043576 | Biological process | 0.04825688 | -1.53029483 | 18 |
| GO:0045777 | Biological process | 0.03481418 | -1.52970522 | 25 |

| | | | | |
|---|---|---|---|---|
| GO:0048483 | Biological process | 0.02406634 | -1.52956397 | 28 |
| GO:0070528 | Biological process | 0.04696841 | -1.5293719 | 24 |
| GO:0098657 | Biological process | 1.11E-05 | -1.52857679 | 170 |
| GO:0032411 | Biological process | 0.00284753 | -1.52855608 | 89 |
| GO:0060080 | Biological process | 0.04550703 | -1.52831548 | 12 |
| GO:2000177 | Biological process | 0.01035593 | -1.52827412 | 64 |
| GO:0003231 | Biological process | 0.00256526 | -1.52798965 | 97 |
| GO:0048747 | Biological process | 0.03138185 | -1.52776343 | 41 |
| GO:0051962 | Biological process | 7.16E-09 | -1.52746967 | 436 |
| GO:0008277 | Biological process | 0.00153374 | -1.52707487 | 100 |
| GO:0038063 | Biological process | 0.04842179 | -1.52704821 | 11 |
| GO:0015695 | Biological process | 0.03697491 | -1.52689887 | 39 |
| GO:0060538 | Biological process | 0.000309 | -1.52638543 | 120 |
| GO:0003013 | Biological process | 7.16E-09 | -1.52584293 | 420 |
| GO:0033189 | Biological process | 0.04895488 | -1.52499457 | 11 |
| GO:0007189 | Biological process | 0.00256526 | -1.5248162 | 93 |
| GO:1904889 | Biological process | 0.049406 | -1.52359861 | 11 |
| GO:0099601 | Biological process | 0.02223458 | -1.52261387 | 49 |
| GO:0003171 | Biological process | 0.0496252 | -1.52261358 | 21 |
| GO:0086005 | Biological process | 0.03746982 | -1.52197031 | 26 |
| GO:0050771 | Biological process | 0.01036144 | -1.52172743 | 61 |
| GO:0048011 | Biological process | 0.03746982 | -1.52157021 | 26 |
| GO:0042551 | Biological process | 0.03593226 | -1.5212521 | 38 |
| GO:0051216 | Biological process | 8.16E-05 | -1.52064616 | 150 |
| GO:0003206 | Biological process | 0.00151823 | -1.52053493 | 95 |
| GO:0031279 | Biological process | 0.03846236 | -1.52051707 | 26 |
| GO:0043268 | Biological process | 0.04842179 | -1.52050904 | 34 |
| GO:0021885 | Biological process | 0.03056688 | -1.51873103 | 40 |
| GO:0007528 | Biological process | 0.03247905 | -1.5181165 | 43 |
| GO:1903350 | Biological process | 0.01216257 | -1.51749571 | 53 |
| GO:0001956 | Biological process | 0.04921569 | -1.51741915 | 17 |
| GO:0048168 | Biological process | 0.03981465 | -1.51651032 | 41 |
| GO:0045661 | Biological process | 0.04248097 | -1.51457251 | 41 |
| GO:1901021 | Biological process | 0.049406 | -1.51450386 | 31 |
| GO:1901385 | Biological process | 0.0325171 | -1.51438598 | 28 |
| GO:0055123 | Biological process | 0.00112442 | -1.5143315 | 111 |
| GO:0003176 | Biological process | 0.0325171 | -1.5140736 | 28 |
| GO:1900271 | Biological process | 0.04526397 | -1.51311252 | 41 |
| GO:0060485 | Biological process | 1.75E-06 | -1.51242505 | 219 |
| GO:0097485 | Biological process | 2.58E-06 | -1.51218289 | 215 |
| GO:0008037 | Biological process | 0.00600559 | -1.51166726 | 88 |
| GO:0051960 | Biological process | 7.16E-09 | -1.51112547 | 742 |
| GO:0002062 | Biological process | 0.00560141 | -1.51033149 | 85 |
| GO:0140115 | Biological process | 0.03593467 | -1.5088765 | 45 |
| GO:0003279 | Biological process | 0.00390967 | -1.50868461 | 93 |
| GO:0007019 | Biological process | 0.04201386 | -1.50764309 | 38 |
| GO:0071869 | Biological process | 0.0088803 | -1.50516192 | 70 |
| GO:0043062 | Biological process | 5.94E-08 | -1.50467504 | 293 |

| | | | | |
|---|---|---|---|---|
| GO:0042471 | Biological process | 0.0064296 | -1.50461591 | 82 |
| GO:0051339 | Biological process | 0.03635891 | -1.50435298 | 28 |
| GO:0030166 | Biological process | 0.02436529 | -1.5031932 | 50 |
| GO:0032835 | Biological process | 0.02138973 | -1.50305228 | 47 |
| GO:0034764 | Biological process | 0.00011245 | -1.50114219 | 159 |
| GO:0097756 | Biological process | 0.02440567 | -1.49859771 | 60 |
| GO:1902305 | Biological process | 0.02551075 | -1.49852021 | 55 |
| GO:0048662 | Biological process | 0.04038709 | -1.49850362 | 40 |
| GO:0090257 | Biological process | 3.15E-05 | -1.49751714 | 181 |
| GO:0015800 | Biological process | 0.02435814 | -1.49737754 | 54 |
| GO:0072511 | Biological process | 7.16E-09 | -1.49654124 | 350 |
| GO:0014013 | Biological process | 0.00519354 | -1.49634634 | 90 |
| GO:0050770 | Biological process | 0.00018527 | -1.49615831 | 153 |
| GO:0032990 | Biological process | 7.16E-09 | -1.49304115 | 574 |
| GO:0030001 | Biological process | 7.16E-09 | -1.49284811 | 637 |
| GO:0051145 | Biological process | 0.03084343 | -1.49241023 | 50 |
| GO:0010975 | Biological process | 7.16E-09 | -1.4919327 | 431 |
| GO:0034329 | Biological process | 1.27E-08 | -1.49136785 | 333 |
| GO:0010771 | Biological process | 0.01244184 | -1.4900807 | 83 |
| GO:0014015 | Biological process | 0.02665192 | -1.49006409 | 57 |
| GO:0003205 | Biological process | 0.00055818 | -1.49004532 | 134 |
| GO:0031345 | Biological process | 0.00032524 | -1.48902569 | 155 |
| GO:0034766 | Biological process | 0.009281 | -1.48820583 | 75 |
| GO:0048640 | Biological process | 0.00741714 | -1.48701336 | 90 |
| GO:0048644 | Biological process | 0.041606 | -1.48594995 | 49 |
| GO:0003281 | Biological process | 0.02440567 | -1.48578063 | 61 |
| GO:1903305 | Biological process | 0.00206034 | -1.48555256 | 126 |
| GO:0045664 | Biological process | 7.16E-09 | -1.48552816 | 540 |
| GO:1990138 | Biological process | 0.00062167 | -1.48539812 | 145 |
| GO:0048645 | Biological process | 0.03968197 | -1.48497161 | 44 |
| GO:0021987 | Biological process | 0.01061338 | -1.4837998 | 87 |
| GO:0007517 | Biological process | 6.55E-07 | -1.48370722 | 284 |
| GO:0007585 | Biological process | 0.03186927 | -1.48244144 | 47 |
| GO:0061564 | Biological process | 7.16E-09 | -1.48189988 | 418 |
| GO:0007267 | Biological process | 7.16E-09 | -1.48126057 | 1300 |
| GO:0098739 | Biological process | 0.00473278 | -1.48030516 | 117 |
| GO:0045666 | Biological process | 2.75E-07 | -1.4795556 | 309 |
| GO:0010977 | Biological process | 0.00177847 | -1.47913109 | 130 |
| GO:0043270 | Biological process | 2.09E-05 | -1.47883185 | 214 |
| GO:0032989 | Biological process | 7.16E-09 | -1.47881195 | 645 |
| GO:0032410 | Biological process | 0.03552992 | -1.47854906 | 60 |
| GO:0042475 | Biological process | 0.02855447 | -1.47839371 | 61 |
| GO:1902904 | Biological process | 0.00174772 | -1.47818557 | 128 |
| GO:0045685 | Biological process | 0.03419201 | -1.47721448 | 54 |
| GO:0042472 | Biological process | 0.02497148 | -1.47670304 | 67 |
| GO:0098655 | Biological process | 7.16E-09 | -1.47626033 | 646 |
| GO:0007423 | Biological process | 7.16E-09 | -1.47507996 | 389 |
| GO:0106027 | Biological process | 0.01669793 | -1.47502038 | 81 |

| | | | | |
|---|---|---|---|---|
| GO:0015672 | Biological process | 7.16E-09 | -1.47445326 | 385 |
| GO:0031102 | Biological process | 0.04578966 | -1.47397818 | 49 |
| GO:0060840 | Biological process | 0.01469293 | -1.47331568 | 75 |
| GO:0021537 | Biological process | 0.0001173 | -1.47195641 | 191 |
| GO:0003208 | Biological process | 0.049406 | -1.47195044 | 44 |
| GO:0006812 | Biological process | 7.16E-09 | -1.47141551 | 843 |
| GO:0042692 | Biological process | 1.47E-06 | -1.47112184 | 280 |
| GO:0003007 | Biological process | 0.00017054 | -1.47111651 | 190 |
| GO:0061061 | Biological process | 7.16E-09 | -1.46993833 | 499 |
| GO:0050772 | Biological process | 0.04038709 | -1.46845472 | 71 |
| GO:0048667 | Biological process | 7.16E-09 | -1.46838587 | 486 |
| GO:0021510 | Biological process | 0.02643864 | -1.46736322 | 62 |
| GO:0045665 | Biological process | 0.0002864 | -1.46733844 | 175 |
| GO:0006940 | Biological process | 0.04114972 | -1.46645699 | 48 |
| GO:0071300 | Biological process | 0.04412238 | -1.46641744 | 50 |
| GO:0035107 | Biological process | 0.00544812 | -1.46630031 | 113 |
| GO:0042733 | Biological process | 0.04707236 | -1.4659454 | 55 |
| GO:0098660 | Biological process | 7.16E-09 | -1.46591115 | 630 |
| GO:0090596 | Biological process | 0.00027315 | -1.46563827 | 178 |
| GO:0019932 | Biological process | 2.79E-07 | -1.464713 | 322 |
| GO:0035150 | Biological process | 0.00813697 | -1.46439184 | 99 |
| GO:0048762 | Biological process | 0.00012801 | -1.46333709 | 180 |
| GO:0048588 | Biological process | 0.00022295 | -1.4621948 | 187 |
| GO:0019233 | Biological process | 0.03398352 | -1.46205691 | 68 |
| GO:0050848 | Biological process | 0.02113189 | -1.46191955 | 83 |
| GO:0021953 | Biological process | 0.00908297 | -1.46107759 | 118 |
| GO:0048562 | Biological process | 6.30E-05 | -1.46100616 | 215 |
| GO:0051261 | Biological process | 0.00845791 | -1.46087681 | 101 |
| GO:0060560 | Biological process | 8.10E-05 | -1.4606821 | 196 |
| GO:0051651 | Biological process | 0.00040457 | -1.4605754 | 175 |
| GO:0019722 | Biological process | 0.00036753 | -1.46036217 | 164 |
| GO:0017156 | Biological process | 0.04988816 | -1.457488 | 50 |
| GO:0055065 | Biological process | 7.16E-09 | -1.45698516 | 466 |
| GO:0090287 | Biological process | 0.00010054 | -1.45636367 | 221 |
| GO:0035113 | Biological process | 0.01211344 | -1.45618644 | 100 |
| GO:0051961 | Biological process | 1.57E-05 | -1.45452827 | 238 |
| GO:0060537 | Biological process | 2.99E-06 | -1.45402593 | 290 |
| GO:0051146 | Biological process | 6.74E-05 | -1.45389052 | 210 |
| GO:0021543 | Biological process | 0.00406617 | -1.45337509 | 128 |
| GO:0003151 | Biological process | 0.04455025 | -1.45318459 | 61 |
| GO:0034763 | Biological process | 0.01778627 | -1.45262932 | 93 |
| GO:0048666 | Biological process | 7.16E-09 | -1.45219443 | 904 |
| GO:0043010 | Biological process | 6.27E-05 | -1.45163829 | 223 |
| GO:0007224 | Biological process | 0.00700965 | -1.45104277 | 114 |
| GO:0060041 | Biological process | 0.0104891 | -1.45016721 | 96 |
| GO:0010721 | Biological process | 2.01E-05 | -1.4496223 | 248 |
| GO:0030182 | Biological process | 7.16E-09 | -1.44945315 | 1065 |
| GO:0060322 | Biological process | 7.16E-09 | -1.44559657 | 605 |

| | | | | |
|---|---|---|---|---|
| GO:0048706 | Biological process | 0.01952656 | -1.44548134 | 93 |
| GO:0007507 | Biological process | 1.70E-08 | -1.44431657 | 444 |
| GO:0014902 | Biological process | 0.03284049 | -1.44338906 | 77 |
| GO:0010769 | Biological process | 3.19E-05 | -1.44321425 | 258 |
| GO:0048634 | Biological process | 0.01367338 | -1.44285566 | 101 |
| GO:0048880 | Biological process | 1.70E-05 | -1.44273011 | 267 |
| GO:0055001 | Biological process | 0.00523831 | -1.44131764 | 130 |
| GO:0008217 | Biological process | 0.00275934 | -1.43987162 | 138 |
| GO:0034220 | Biological process | 7.16E-09 | -1.43981697 | 855 |
| GO:0006814 | Biological process | 0.00102663 | -1.43927907 | 168 |
| GO:0010976 | Biological process | 5.81E-05 | -1.43892366 | 249 |
| GO:0030178 | Biological process | 0.00177847 | -1.43751153 | 179 |
| GO:0007600 | Biological process | 5.95E-06 | -1.43726937 | 298 |
| GO:0022008 | Biological process | 7.16E-09 | -1.43675158 | 1272 |
| GO:0035725 | Biological process | 0.00503207 | -1.43630558 | 129 |
| GO:1901879 | Biological process | 0.04126538 | -1.43568579 | 79 |
| GO:0060284 | Biological process | 7.16E-09 | -1.43491831 | 749 |
| GO:0045165 | Biological process | 0.00253551 | -1.43396117 | 149 |
| GO:0007186 | Biological process | 7.16E-09 | -1.43134103 | 538 |
| GO:0007417 | Biological process | 7.16E-09 | -1.43028949 | 770 |
| GO:0048593 | Biological process | 0.03366306 | -1.42958199 | 87 |
| GO:0022612 | Biological process | 0.02113189 | -1.42838233 | 100 |
| GO:0000904 | Biological process | 7.16E-09 | -1.42737229 | 610 |
| GO:0051147 | Biological process | 0.0104891 | -1.4263903 | 126 |
| GO:0048736 | Biological process | 0.00640154 | -1.42604707 | 134 |
| GO:0021700 | Biological process | 0.00033693 | -1.42588501 | 213 |
| GO:0031110 | Biological process | 0.04236705 | -1.42442582 | 76 |
| GO:0048738 | Biological process | 0.00593108 | -1.42259181 | 157 |
| GO:0031344 | Biological process | 7.16E-09 | -1.41753232 | 585 |
| GO:0006873 | Biological process | 4.32E-08 | -1.41709343 | 483 |
| GO:0030900 | Biological process | 6.46E-05 | -1.41647054 | 275 |
| GO:0071772 | Biological process | 0.03105429 | -1.41557718 | 118 |
| GO:0090288 | Biological process | 0.01491871 | -1.41049979 | 122 |
| GO:0048705 | Biological process | 0.0039473 | -1.41009024 | 159 |
| GO:0061448 | Biological process | 0.00091662 | -1.4099903 | 203 |
| GO:0017157 | Biological process | 0.0028662 | -1.40534012 | 176 |
| GO:0048592 | Biological process | 0.04113802 | -1.40460215 | 105 |
| GO:0023061 | Biological process | 2.31E-07 | -1.40399924 | 448 |
| GO:0010720 | Biological process | 3.89E-07 | -1.40398704 | 447 |
| GO:0051494 | Biological process | 0.01160874 | -1.40248305 | 135 |
| GO:0001764 | Biological process | 0.02643864 | -1.40242957 | 116 |
| GO:1905114 | Biological process | 9.68E-08 | -1.39993201 | 498 |
| GO:0001501 | Biological process | 7.73E-06 | -1.39836685 | 381 |
| GO:0048598 | Biological process | 2.71E-07 | -1.39727302 | 452 |
| GO:0000902 | Biological process | 7.16E-09 | -1.38898141 | 841 |
| GO:0001649 | Biological process | 0.00482955 | -1.38680326 | 176 |
| GO:0050679 | Biological process | 0.01199827 | -1.38640071 | 155 |
| GO:0060348 | Biological process | 0.0123465 | -1.38548824 | 155 |

| | | | | |
|---|---|---|---|---|
| GO:0048863 | Biological process | 0.00240679 | -1.38440029 | 210 |
| GO:0035051 | Biological process | 0.04223084 | -1.38217871 | 108 |
| GO:0006865 | Biological process | 0.03593226 | -1.3818369 | 115 |
| GO:0006023 | Biological process | 0.04704893 | -1.38177505 | 94 |
| GO:0009887 | Biological process | 7.16E-09 | -1.38131401 | 780 |
| GO:0031346 | Biological process | 6.09E-05 | -1.38091375 | 336 |
| GO:0060070 | Biological process | 0.00060394 | -1.37972925 | 271 |
| GO:0035265 | Biological process | 0.03593226 | -1.37811507 | 127 |
| GO:0006811 | Biological process | 7.16E-09 | -1.37532253 | 1227 |
| GO:0048638 | Biological process | 0.0012568 | -1.37513992 | 255 |
| GO:0045596 | Biological process | 5.96E-07 | -1.37304916 | 539 |
| GO:0001503 | Biological process | 0.00027324 | -1.37234674 | 313 |
| GO:0090101 | Biological process | 0.04704893 | -1.37221846 | 96 |
| GO:0042063 | Biological process | 0.00237051 | -1.36855367 | 236 |
| GO:0055085 | Biological process | 7.16E-09 | -1.36689204 | 1182 |
| GO:0022604 | Biological process | 1.13E-05 | -1.36499794 | 421 |
| GO:0030030 | Biological process | 7.16E-09 | -1.36491598 | 1272 |
| GO:0007389 | Biological process | 0.00032524 | -1.36478457 | 296 |
| GO:0050801 | Biological process | 4.12E-07 | -1.36385054 | 596 |
| GO:0010001 | Biological process | 0.00946751 | -1.36091289 | 181 |
| GO:0003018 | Biological process | 0.02665192 | -1.36090947 | 137 |
| GO:0022610 | Biological process | 7.16E-09 | -1.35953361 | 1061 |
| GO:0030308 | Biological process | 0.02600597 | -1.35740885 | 152 |
| GO:0001935 | Biological process | 0.0434146 | -1.35430955 | 128 |
| GO:0048568 | Biological process | 0.00036753 | -1.34997462 | 339 |
| GO:0071695 | Biological process | 0.01812017 | -1.34867437 | 169 |
| GO:0072359 | Biological process | 7.16E-09 | -1.34747487 | 874 |
| GO:0001655 | Biological process | 0.00353985 | -1.34728166 | 258 |
| GO:2000027 | Biological process | 0.01311441 | -1.34423449 | 197 |
| GO:0048017 | Biological process | 0.04101383 | -1.34378607 | 154 |
| GO:0045597 | Biological process | 3.67E-08 | -1.34307627 | 787 |
| GO:0051093 | Biological process | 1.43E-07 | -1.34266164 | 734 |
| GO:0030278 | Biological process | 0.03633427 | -1.33832121 | 153 |
| GO:0090092 | Biological process | 0.01947787 | -1.33479019 | 189 |
| GO:0045595 | Biological process | 7.16E-09 | -1.33360181 | 1441 |
| GO:0033002 | Biological process | 0.02900511 | -1.33308767 | 160 |
| GO:0016358 | Biological process | 0.01669793 | -1.33066955 | 206 |
| GO:1990778 | Biological process | 0.00220536 | -1.32800926 | 301 |
| GO:0009612 | Biological process | 0.03755888 | -1.32520649 | 167 |
| GO:0001944 | Biological process | 1.04E-05 | -1.32246094 | 600 |
| GO:0030111 | Biological process | 0.00275934 | -1.32186365 | 306 |
| GO:0051094 | Biological process | 7.16E-09 | -1.3216475 | 1084 |
| GO:0070848 | Biological process | 3.31E-05 | -1.3204094 | 561 |
| GO:0019725 | Biological process | 2.39E-06 | -1.31690463 | 717 |
| GO:0048729 | Biological process | 0.00013469 | -1.31416231 | 511 |
| GO:0045926 | Biological process | 0.02728038 | -1.31253289 | 196 |
| GO:0051049 | Biological process | 7.16E-09 | -1.30836386 | 1403 |
| GO:0003002 | Biological process | 0.02026405 | -1.30671257 | 227 |

| | | | | |
|---|---|---|---|---|
| GO:0048589 | Biological process | 0.00036753 | -1.3036167 | 512 |
| GO:0060562 | Biological process | 0.01709122 | -1.30127893 | 258 |
| GO:0048514 | Biological process | 0.00037166 | -1.30124397 | 498 |
| GO:0042330 | Biological process | 0.00050296 | -1.2979398 | 475 |
| GO:0031589 | Biological process | 0.01061338 | -1.29648405 | 304 |
| GO:0051235 | Biological process | 0.02223458 | -1.29633881 | 269 |
| GO:0098609 | Biological process | 8.64E-05 | -1.29287571 | 620 |
| GO:0048646 | Biological process | 2.39E-06 | -1.2921749 | 863 |
| GO:0035239 | Biological process | 2.85E-05 | -1.28818378 | 688 |
| GO:0051270 | Biological process | 8.48E-06 | -1.28107291 | 823 |
| GO:0032970 | Biological process | 0.01631121 | -1.28046285 | 331 |
| GO:0022603 | Biological process | 8.30E-06 | -1.27977145 | 893 |
| GO:0035295 | Biological process | 2.13E-05 | -1.27667323 | 837 |
| GO:0072659 | Biological process | 0.049406 | -1.26947464 | 255 |
| GO:1902903 | Biological process | 0.02440567 | -1.26912443 | 315 |
| GO:0051272 | Biological process | 0.002624 | -1.26844781 | 470 |
| GO:0097435 | Biological process | 0.00136442 | -1.26328409 | 590 |
| GO:0030029 | Biological process | 0.00038313 | -1.26287205 | 653 |
| GO:0060341 | Biological process | 4.51E-05 | -1.26092878 | 847 |
| GO:0002009 | Biological process | 0.00646177 | -1.25710829 | 443 |
| GO:0009790 | Biological process | 7.99E-05 | -1.25595133 | 821 |
| GO:0016049 | Biological process | 0.00784867 | -1.25595115 | 399 |
| GO:0051240 | Biological process | 6.49E-07 | -1.25303615 | 1366 |
| GO:0048870 | Biological process | 1.78E-06 | -1.2530329 | 1257 |
| GO:0090066 | Biological process | 0.00738759 | -1.24857288 | 413 |
| GO:0198738 | Biological process | 0.01883872 | -1.24700918 | 422 |
| GO:0072657 | Biological process | 0.00326757 | -1.24690825 | 577 |
| GO:0001667 | Biological process | 0.04328162 | -1.2437892 | 344 |
| GO:0051046 | Biological process | 0.0085123 | -1.23572274 | 545 |
| GO:0007167 | Biological process | 0.00067296 | -1.23033099 | 853 |
| GO:0040008 | Biological process | 0.01153053 | -1.22114706 | 528 |
| GO:0060627 | Biological process | 0.0248504 | -1.21565144 | 449 |
| GO:0040007 | Biological process | 0.00389997 | -1.21424781 | 765 |
| GO:0051241 | Biological process | 0.0017235 | -1.20973588 | 929 |
| GO:0048878 | Biological process | 0.00312672 | -1.20949193 | 879 |
| GO:0044087 | Biological process | 0.0049504 | -1.20734077 | 818 |
| GO:0008284 | Biological process | 0.00977624 | -1.20172397 | 673 |
| GO:0023056 | Biological process | 7.78E-05 | -1.2016284 | 1443 |
| GO:0009611 | Biological process | 0.03862418 | -1.19962337 | 509 |
| GO:0051050 | Biological process | 0.01036144 | -1.19436219 | 745 |
| GO:0051493 | Biological process | 0.03982617 | -1.19122512 | 474 |
| GO:0007169 | Biological process | 0.04328162 | -1.18984418 | 605 |
| GO:0051130 | Biological process | 0.01524939 | -1.16112705 | 1023 |
| GO:0042127 | Biological process | 0.00732104 | -1.16026082 | 1245 |
| GO:0023057 | Biological process | 0.02009484 | -1.15716681 | 1132 |
| GO:0071495 | Biological process | 0.02440567 | -1.15464869 | 1060 |
| GO:0007010 | Biological process | 0.02455333 | -1.14682817 | 1133 |
| GO:0060429 | Biological process | 0.04987734 | -1.14129363 | 890 |

| | | | | |
|---|---|---|---|---|
| GO:0009719 | Biological process | 0.03589778 | -1.12848659 | 1265 |
| GO:0002832 | Biological process | 0.045028 | 1.43678152 | 76 |
| GO:0055092 | Biological process | 0.02441186 | 1.54646723 | 69 |
| GO:0097006 | Biological process | 0.04633114 | 1.55503768 | 74 |
| GO:0060333 | Biological process | 0.04412238 | 1.56294728 | 74 |
| GO:0031424 | Biological process | 0.04707236 | 1.56621167 | 44 |
| GO:0060760 | Biological process | 0.03853316 | 1.57797677 | 51 |
| GO:0034381 | Biological process | 0.03717109 | 1.60609591 | 44 |
| GO:0007586 | Biological process | 0.00558481 | 1.65180319 | 85 |
| GO:0002753 | Biological process | 0.03104276 | 1.65686115 | 61 |
| GO:0046503 | Biological process | 0.00839066 | 1.72133413 | 55 |
| GO:0009072 | Biological process | 0.04770379 | 1.72467804 | 26 |
| GO:0050892 | Biological process | 0.01827796 | 1.84236255 | 32 |
| GO:0006641 | Biological process | 0.00039976 | 1.86673779 | 76 |
| GO:0034143 | Biological process | 0.03188257 | 1.8730902 | 15 |
| GO:0044241 | Biological process | 0.04262721 | 1.88123023 | 12 |
| GO:0042304 | Biological process | 0.00481621 | 1.89750499 | 41 |
| GO:0010984 | Biological process | 0.02440567 | 1.9062174 | 15 |
| GO:0034370 | Biological process | 0.04566297 | 1.92516477 | 10 |
| GO:0019731 | Biological process | 0.01311441 | 1.96701995 | 18 |
| GO:0090207 | Biological process | 0.00640154 | 1.97000611 | 32 |
| GO:0019433 | Biological process | 0.00389997 | 2.03269544 | 26 |
| GO:0061365 | Biological process | 0.01601713 | 2.04408459 | 10 |
| GO:0030277 | Biological process | 0.00558481 | 2.04581695 | 14 |
| GO:0046461 | Biological process | 0.0012417 | 2.09817928 | 33 |
| GO:0090208 | Biological process | 0.00549587 | 2.09898317 | 19 |
| GO:0098856 | Biological process | 0.00588382 | 2.10434705 | 13 |
| GO:0032480 | Biological process | 9.75E-05 | 2.16415277 | 39 |
| GO:0010896 | Biological process | 0.00027324 | 2.2653967 | 11 |
| GO:0034340 | Biological process | 6.40E-07 | 2.28973293 | 75 |
| GO:0045211 | Cellular component | 2.95E-09 | -1.76942886 | 179 |
| GO:0034703 | Cellular component | 2.95E-09 | -1.75198893 | 145 |
| GO:0097060 | Cellular component | 2.95E-09 | -1.74833507 | 262 |
| GO:0099240 | Cellular component | 2.95E-09 | -1.73891702 | 106 |
| GO:0043204 | Cellular component | 2.95E-09 | -1.72698866 | 124 |
| GO:0044306 | Cellular component | 2.95E-09 | -1.71092059 | 119 |
| GO:1990351 | Cellular component | 2.95E-09 | -1.69686795 | 220 |
| GO:0098794 | Cellular component | 2.95E-09 | -1.65229512 | 503 |
| GO:0098978 | Cellular component | 2.95E-09 | -1.64008525 | 292 |
| GO:0098793 | Cellular component | 2.95E-09 | -1.61892864 | 402 |
| GO:0045202 | Cellular component | 2.95E-09 | -1.61845973 | 1044 |
| GO:0098984 | Cellular component | 2.95E-09 | -1.59846223 | 304 |
| GO:0044297 | Cellular component | 2.95E-09 | -1.59556937 | 482 |
| GO:0150034 | Cellular component | 2.95E-09 | -1.58717344 | 255 |
| GO:0030424 | Cellular component | 2.95E-09 | -1.58716107 | 524 |
| GO:0036477 | Cellular component | 2.95E-09 | -1.58143476 | 675 |
| GO:0097447 | Cellular component | 2.95E-09 | -1.52950062 | 486 |
| GO:0098797 | Cellular component | 2.95E-09 | -1.50997708 | 414 |

| | | | | |
|---|---|---|---|---|
| GO:0043005 | Cellular component | 2.95E-09 | -1.50649123 | 1022 |
| GO:0031226 | Cellular component | 2.95E-09 | -1.50078847 | 1076 |
| GO:0098590 | Cellular component | 2.95E-09 | -1.39637954 | 937 |
| GO:0042734 | Cellular component | 1.29E-08 | -1.70336242 | 107 |
| GO:0042383 | Cellular component | 4.27E-08 | -1.6732618 | 111 |
| GO:0031012 | Cellular component | 4.58E-08 | -1.43687743 | 381 |
| GO:0044309 | Cellular component | 1.33E-07 | -1.59756471 | 148 |
| GO:0099081 | Cellular component | 2.01E-07 | -1.32892601 | 670 |
| GO:0098936 | Cellular component | 5.38E-07 | -1.71269167 | 74 |
| GO:0099634 | Cellular component | 1.33E-06 | -1.68792673 | 77 |
| GO:0009986 | Cellular component | 1.79E-06 | -1.3404372 | 563 |
| GO:0034705 | Cellular component | 2.78E-06 | -1.72486118 | 59 |
| GO:0062023 | Cellular component | 2.78E-06 | -1.42246393 | 309 |
| GO:0031674 | Cellular component | 3.00E-06 | -1.61990335 | 106 |
| GO:0043235 | Cellular component | 5.90E-06 | -1.42862387 | 290 |
| GO:0043292 | Cellular component | 1.09E-05 | -1.51334803 | 173 |
| GO:0098839 | Cellular component | 1.15E-05 | -1.68137213 | 63 |
| GO:0016528 | Cellular component | 1.66E-05 | -1.65941435 | 66 |
| GO:0098796 | Cellular component | 1.66E-05 | -1.26151179 | 938 |
| GO:0098589 | Cellular component | 4.94E-05 | -1.39451637 | 287 |
| GO:0048786 | Cellular component | 6.98E-05 | -1.6650471 | 62 |
| GO:0070382 | Cellular component | 0.00011432 | -1.47047364 | 177 |
| GO:0043195 | Cellular component | 0.00013425 | -1.67749925 | 44 |
| GO:0044304 | Cellular component | 0.00013425 | -1.65032151 | 54 |
| GO:0098948 | Cellular component | 0.00015106 | -1.67040106 | 46 |
| GO:0098982 | Cellular component | 0.00016868 | -1.67780578 | 41 |
| GO:0099146 | Cellular component | 0.00027528 | -1.68345734 | 35 |
| GO:0098889 | Cellular component | 0.00034518 | -1.62426573 | 54 |
| GO:0099080 | Cellular component | 0.00034518 | -1.21057745 | 934 |
| GO:0098878 | Cellular component | 0.00045973 | -1.66494557 | 32 |
| GO:0030672 | Cellular component | 0.00051848 | -1.54272626 | 80 |
| GO:0034385 | Cellular component | 0.00062161 | 2.15713125 | 17 |
| GO:0030315 | Cellular component | 0.00074688 | -1.65331556 | 43 |
| GO:0098685 | Cellular component | 0.00074688 | -1.56438218 | 70 |
| GO:0031594 | Cellular component | 0.00077903 | -1.56352572 | 66 |
| GO:0001533 | Cellular component | 0.0010629 | 2.12009596 | 17 |
| GO:0060076 | Cellular component | 0.00110368 | -1.6525021 | 31 |
| GO:0033017 | Cellular component | 0.00136045 | -1.64673838 | 31 |
| GO:0030427 | Cellular component | 0.00202047 | -1.40737156 | 164 |
| GO:0043198 | Cellular component | 0.00250306 | -1.60599839 | 33 |
| GO:0034704 | Cellular component | 0.00365963 | -1.55382461 | 49 |
| GO:0098802 | Cellular component | 0.00376869 | -1.41583155 | 140 |
| GO:0099568 | Cellular component | 0.00447351 | -1.37092985 | 186 |
| GO:0030133 | Cellular component | 0.00447351 | -1.29204075 | 330 |
| GO:0015629 | Cellular component | 0.00447351 | -1.26252752 | 406 |
| GO:0044853 | Cellular component | 0.00468153 | -1.48115139 | 92 |
| GO:0005911 | Cellular component | 0.00468153 | -1.26240186 | 395 |
| GO:0014704 | Cellular component | 0.00494319 | -1.57151938 | 42 |

| | | | | |
|---|---|---|---|---|
| GO:0035578 | Cellular component | 0.00519977 | 1.79164541 | 75 |
| GO:0031209 | Cellular component | 0.00528529 | -1.58912708 | 10 |
| GO:0032994 | Cellular component | 0.00528529 | 1.92085809 | 28 |
| GO:0090665 | Cellular component | 0.00538172 | -1.62405405 | 17 |
| GO:0044291 | Cellular component | 0.00545675 | -1.53082246 | 60 |
| GO:0005901 | Cellular component | 0.00625264 | -1.50187777 | 68 |
| GO:0031256 | Cellular component | 0.00685683 | -1.38421329 | 145 |
| GO:0032838 | Cellular component | 0.00812826 | -1.38865529 | 151 |
| GO:0070161 | Cellular component | 0.00847109 | -1.18858763 | 717 |
| GO:0042627 | Cellular component | 0.00959656 | 2.03841544 | 10 |
| GO:0034706 | Cellular component | 0.01011092 | -1.62510141 | 19 |
| GO:0034364 | Cellular component | 0.01027318 | 1.90933162 | 17 |
| GO:0098644 | Cellular component | 0.01312308 | -1.56599242 | 16 |
| GO:0048787 | Cellular component | 0.01314797 | -1.54965796 | 26 |
| GO:0005583 | Cellular component | 0.01478033 | -1.55539309 | 10 |
| GO:0030285 | Cellular component | 0.01531845 | -1.53963701 | 24 |
| GO:0005811 | Cellular component | 0.01531845 | 1.61276083 | 82 |
| GO:0031252 | Cellular component | 0.01683969 | -1.24356965 | 378 |
| GO:0042824 | Cellular component | 0.0172694 | 1.99219415 | 10 |
| GO:0005604 | Cellular component | 0.01746914 | -1.42936812 | 83 |
| GO:0012507 | Cellular component | 0.01811352 | 1.8107233 | 50 |
| GO:0001518 | Cellular component | 0.01837559 | -1.55990337 | 12 |
| GO:0005891 | Cellular component | 0.01862798 | -1.50343507 | 29 |
| GO:0030134 | Cellular component | 0.02616356 | 1.6045832 | 76 |
| GO:0032281 | Cellular component | 0.02644198 | -1.53846437 | 15 |
| GO:0044305 | Cellular component | 0.02661778 | -1.57024534 | 19 |
| GO:0043034 | Cellular component | 0.02661778 | -1.52991846 | 18 |
| GO:0098945 | Cellular component | 0.02966539 | -1.52288186 | 16 |
| GO:0099513 | Cellular component | 0.02996753 | -1.20024495 | 497 |
| GO:0044298 | Cellular component | 0.03235041 | -1.51345939 | 21 |
| GO:0098803 | Cellular component | 0.03235041 | 1.53367533 | 77 |
| GO:0032839 | Cellular component | 0.0350039 | -1.48417218 | 30 |
| GO:0030964 | Cellular component | 0.0350039 | 1.55784136 | 48 |
| GO:0030027 | Cellular component | 0.03532002 | -1.28729084 | 188 |
| GO:0030863 | Cellular component | 0.03574641 | -1.39582108 | 90 |
| GO:0031045 | Cellular component | 0.03706533 | -1.55567809 | 19 |
| GO:0031430 | Cellular component | 0.03851414 | -1.50169127 | 21 |
| GO:0032589 | Cellular component | 0.03873292 | -1.47406326 | 40 |
| GO:0098799 | Cellular component | 0.04095409 | 1.72501394 | 22 |
| GO:0031225 | Cellular component | 0.0410716 | -1.35241474 | 96 |
| GO:0005929 | Cellular component | 0.0410716 | -1.19156342 | 421 |
| GO:0120111 | Cellular component | 0.04222857 | -1.37677177 | 80 |
| GO:0098686 | Cellular component | 0.04235002 | -1.48098802 | 26 |
| GO:0045259 | Cellular component | 0.04247879 | 1.78557127 | 19 |
| GO:0042611 | Cellular component | 0.04296381 | 1.75009389 | 16 |
| GO:0031253 | Cellular component | 0.04462015 | -1.23711783 | 277 |
| GO:0043083 | Cellular component | 0.04527661 | -1.50771961 | 12 |
| GO:0030175 | Cellular component | 0.04527661 | -1.34231769 | 95 |

| | | | | |
|---|---|---|---|---|
| GO:0005244 | Molecular function | 1.04E-08 | -1.70256164 | 123 |
| GO:0022836 | Molecular function | 1.04E-08 | -1.69848205 | 206 |
| GO:0005261 | Molecular function | 1.04E-08 | -1.68510605 | 213 |
| GO:0022803 | Molecular function | 1.04E-08 | -1.59585481 | 306 |
| GO:0046873 | Molecular function | 1.04E-08 | -1.55442237 | 287 |
| GO:0060089 | Molecular function | 1.04E-08 | -1.48701757 | 605 |
| GO:0008324 | Molecular function | 1.04E-08 | -1.48107435 | 437 |
| GO:0005509 | Molecular function | 1.04E-08 | -1.4409129 | 461 |
| GO:0015075 | Molecular function | 1.04E-08 | -1.40985607 | 606 |
| GO:0005198 | Molecular function | 1.12E-07 | -1.4104776 | 492 |
| GO:0005215 | Molecular function | 1.21E-07 | -1.32526905 | 840 |
| GO:0008092 | Molecular function | 2.83E-07 | -1.32213791 | 806 |
| GO:0015079 | Molecular function | 3.61E-07 | -1.6721462 | 93 |
| GO:0022843 | Molecular function | 3.99E-07 | -1.70183138 | 82 |
| GO:0004930 | Molecular function | 3.99E-07 | -1.54062734 | 193 |
| GO:0005516 | Molecular function | 5.25E-07 | -1.57623925 | 160 |
| GO:0015077 | Molecular function | 1.08E-06 | -1.48527426 | 253 |
| GO:0005267 | Molecular function | 2.15E-06 | -1.70972638 | 67 |
| GO:0015276 | Molecular function | 1.11E-05 | -1.65680835 | 80 |
| GO:0005201 | Molecular function | 1.96E-05 | -1.59162429 | 114 |
| GO:0044325 | Molecular function | 5.55E-05 | -1.56180462 | 113 |
| GO:0097157 | Molecular function | 7.00E-05 | -1.72261821 | 11 |
| GO:0005249 | Molecular function | 0.00010603 | -1.67436012 | 50 |
| GO:0030594 | Molecular function | 0.00012071 | -1.72859394 | 40 |
| GO:0005102 | Molecular function | 0.000124 | -1.22561001 | 1108 |
| GO:0015085 | Molecular function | 0.00021794 | -1.56625549 | 94 |
| GO:0016247 | Molecular function | 0.00021794 | -1.53826162 | 110 |
| GO:0099094 | Molecular function | 0.00023483 | -1.60644541 | 69 |
| GO:0072509 | Molecular function | 0.00064629 | -1.50024528 | 120 |
| GO:0099106 | Molecular function | 0.00075962 | -1.54336331 | 90 |
| GO:0030020 | Molecular function | 0.00077777 | -1.68980821 | 29 |
| GO:0098960 | Molecular function | 0.00078913 | -1.68839663 | 29 |
| GO:0022835 | Molecular function | 0.00086341 | -1.68106503 | 28 |
| GO:0005272 | Molecular function | 0.00086341 | -1.66893806 | 33 |
| GO:0005539 | Molecular function | 0.00086341 | -1.43466481 | 162 |
| GO:0001653 | Molecular function | 0.00106455 | -1.58426586 | 65 |
| GO:0005230 | Molecular function | 0.00223833 | -1.64269088 | 37 |
| GO:0099529 | Molecular function | 0.00234609 | -1.65576414 | 26 |
| GO:0015631 | Molecular function | 0.00239089 | -1.33245962 | 303 |
| GO:0004970 | Molecular function | 0.00517012 | -1.6341946 | 11 |
| GO:0008066 | Molecular function | 0.00517012 | -1.6341946 | 11 |
| GO:0019199 | Molecular function | 0.00635287 | -1.52056436 | 70 |
| GO:0005246 | Molecular function | 0.00653101 | -1.61259665 | 36 |
| GO:0015081 | Molecular function | 0.00677018 | -1.45701317 | 103 |
| GO:0005248 | Molecular function | 0.00686653 | -1.62070424 | 18 |
| GO:0003954 | Molecular function | 0.00686653 | 1.99090091 | 44 |
| GO:0008195 | Molecular function | 0.00817407 | -1.61334161 | 12 |
| GO:0030545 | Molecular function | 0.0102475 | -1.30815093 | 274 |

| | | | | |
|---|---|---|---|---|
| GO:0030276 | Molecular function | 0.01121945 | -1.5130676 | 59 |
| GO:0016655 | Molecular function | 0.01121945 | 1.83849343 | 53 |
| GO:0008017 | Molecular function | 0.01234072 | -1.33518063 | 220 |
| GO:0003779 | Molecular function | 0.01274933 | -1.27413638 | 345 |
| GO:0004143 | Molecular function | 0.0130602 | -1.58869401 | 10 |
| GO:0005245 | Molecular function | 0.0130602 | -1.58160815 | 28 |
| GO:0005518 | Molecular function | 0.01485405 | -1.48605777 | 63 |
| GO:0051018 | Molecular function | 0.0158376 | -1.5303568 | 44 |
| GO:0019838 | Molecular function | 0.02256206 | -1.40060194 | 118 |
| GO:0008307 | Molecular function | 0.02271404 | -1.5620524 | 30 |
| GO:0015459 | Molecular function | 0.02391774 | -1.54704519 | 36 |
| GO:0000149 | Molecular function | 0.03639626 | -1.39179666 | 96 |
| GO:0005251 | Molecular function | 0.03747855 | -1.54638722 | 18 |
| GO:0008146 | Molecular function | 0.04393674 | -1.53805049 | 32 |
| GO:0004714 | Molecular function | 0.04393674 | -1.48131609 | 54 |
| GO:0017080 | Molecular function | 0.04974294 | -1.50646909 | 33 |

**Supplementary Figure 1.** Vulcano plot of KO genes detected in shotgun sequencing. Differentially abundand KO genes (padj < 0.05) have been highlighted in blue and red for lower and higher abundance in anaemia, respectively. Non significant genes have been coloured in black, according to the legend.

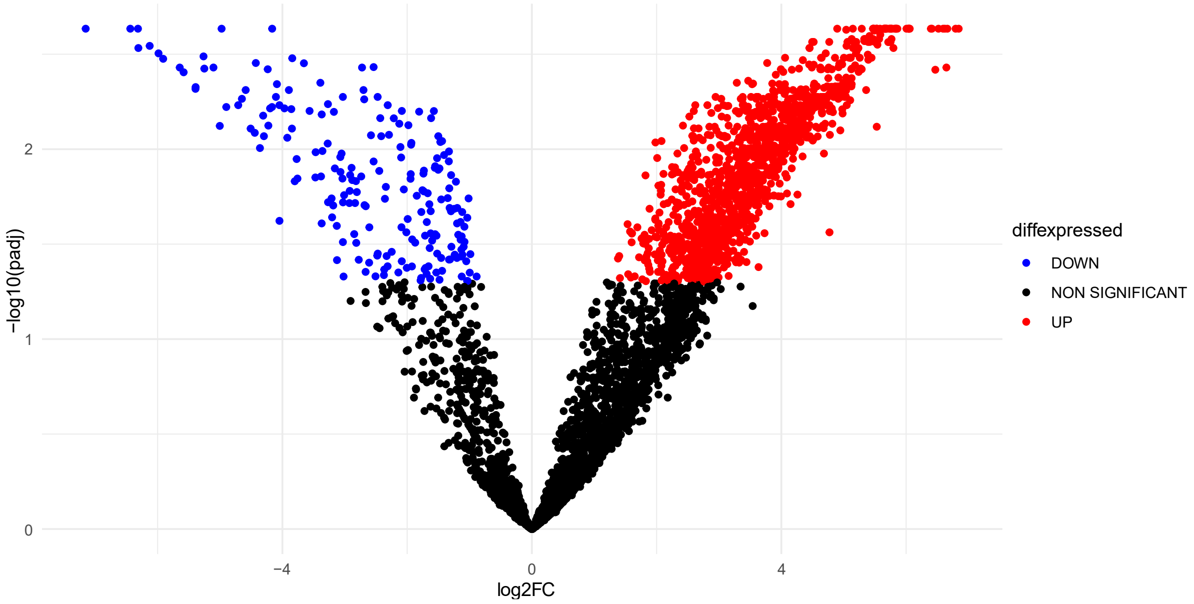

**Supplementary Figure 2**. Biosynthesis of amino acids pathway (KEGG) coloured according to $log_2FC$ values of statistically significant KO genes. Colour legend: amino acids – producing pathways with increased relative abundance during IDA (green squares); amino acids – producing pathways with decreased relative abundance during IDA (orange squares)

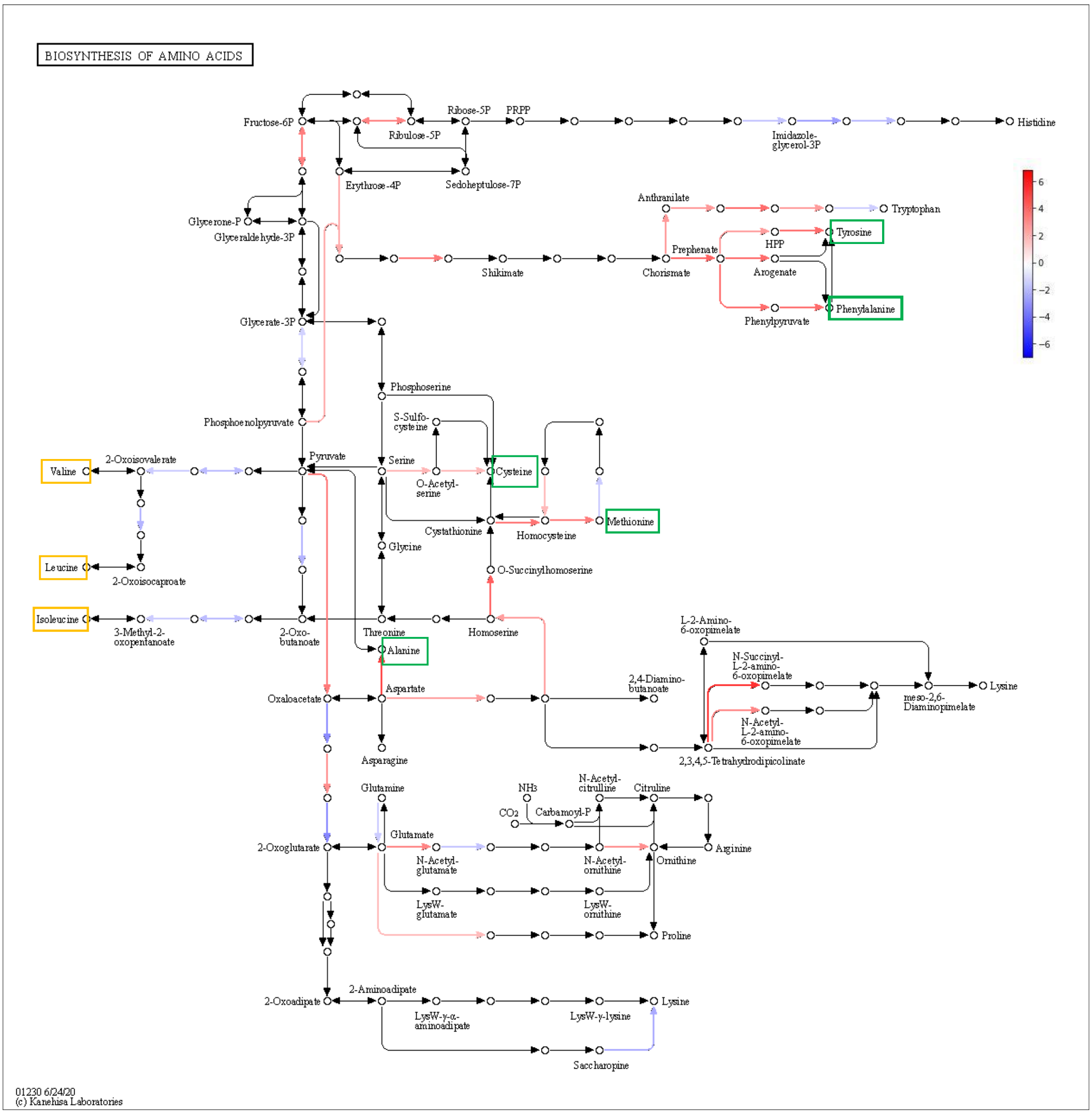

**Supplementary Figure 3.** Amino sugar and nucleotide metabolism pathway (KEGG) coloured according to log$_2$FC values of statistically significant KO genes. Colour legend: synthesis of cell wall components (green), metabolism of sugar intermediates (blue)

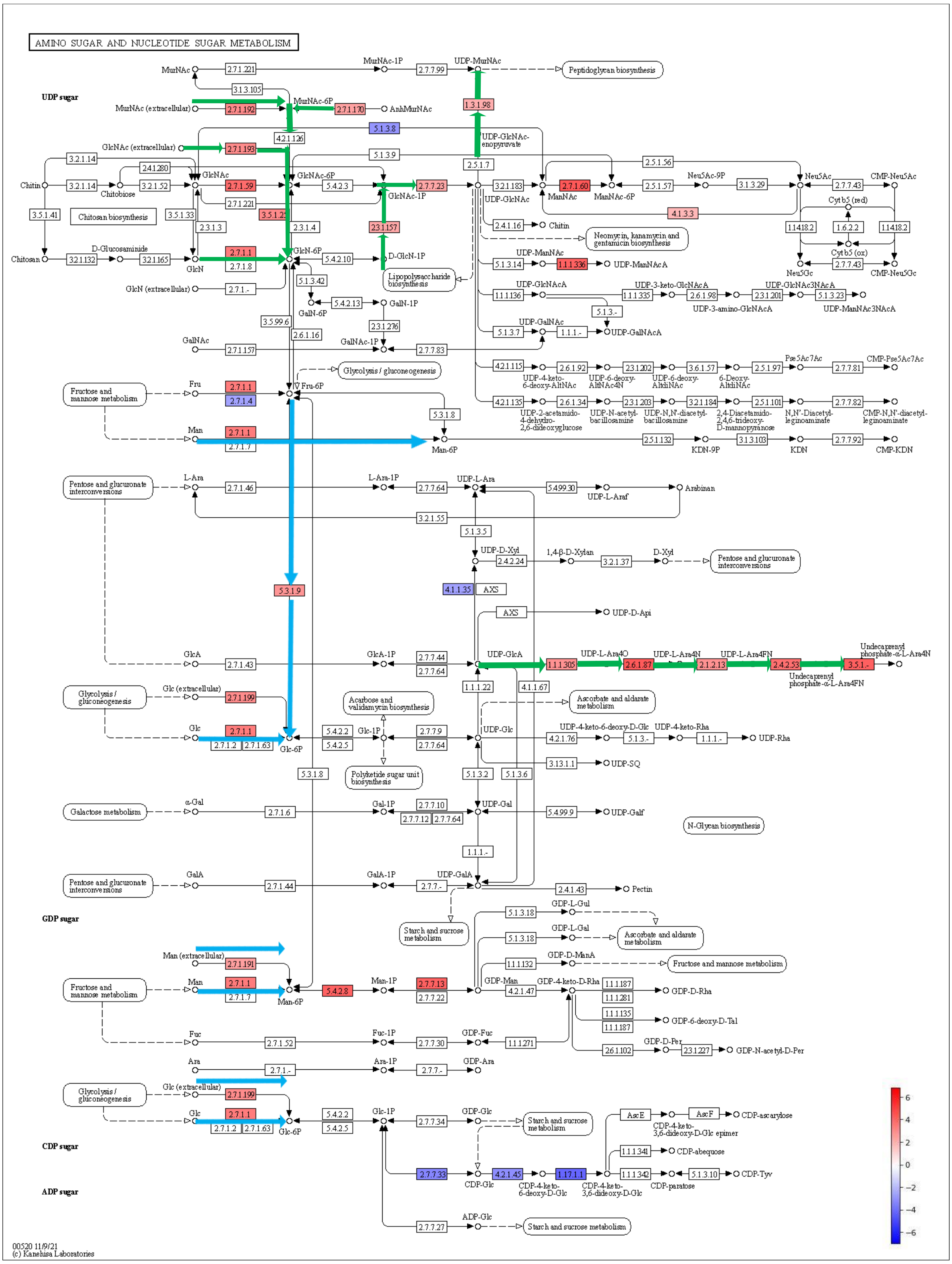

**Supplementary Figure 4.** Glycolysis and gluconeogenesis pathway (KEGG) coloured according to $\log_2$FC values of statistically significant KO genes. Colour legend: metabolism of carbohydrates and sugar intermediates towards the formation of pyruvate and acetyl-CoA (blue)

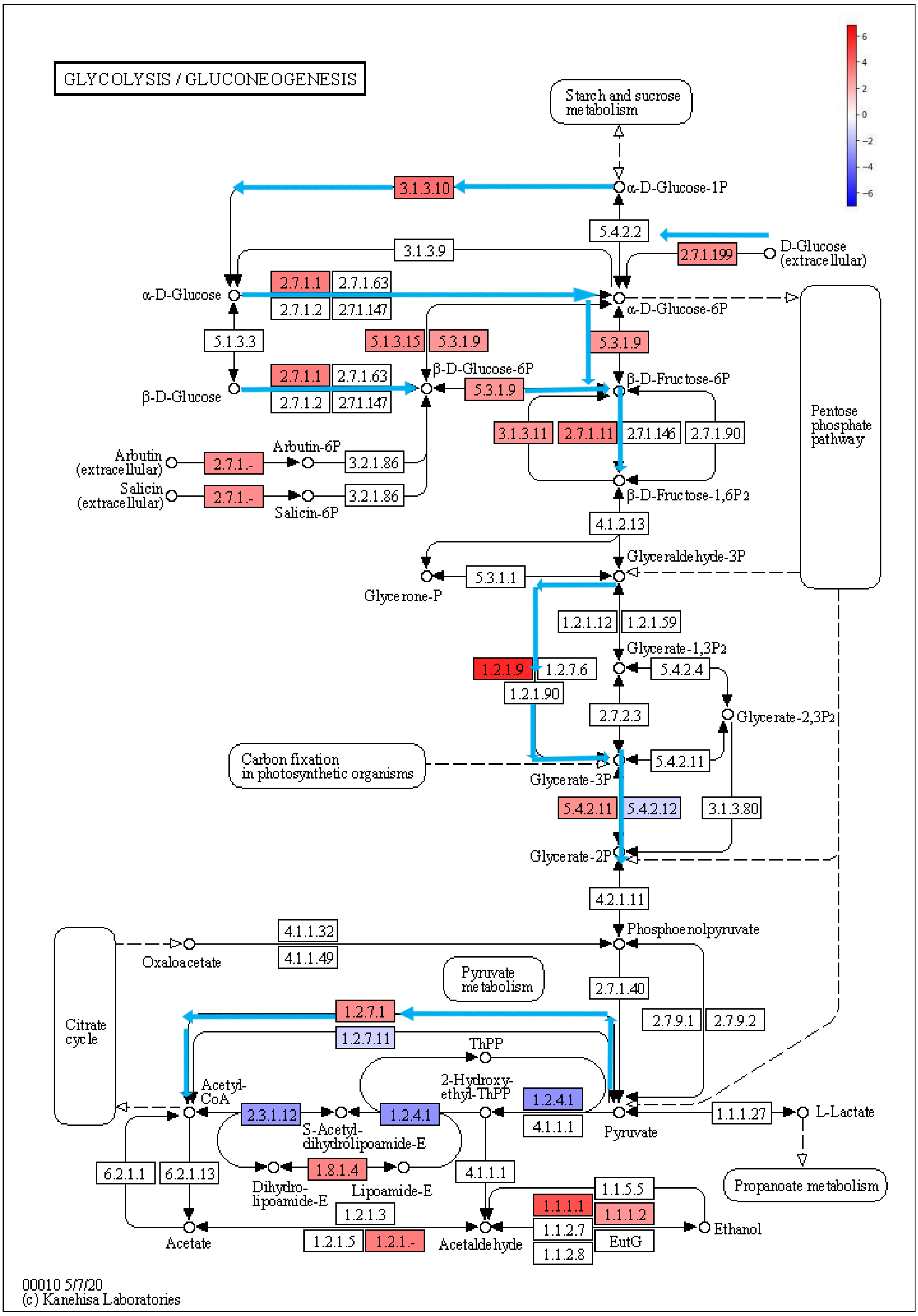

**Supplementary Figure 5.** Pyruvate metabolism pathway (KEGG) coloured according to $log_2$FC values of statistically significant KO genes. Colour legend: metabolism of pyruvate towards the formation for formate and succinate (blue), lactate and acetate (green)

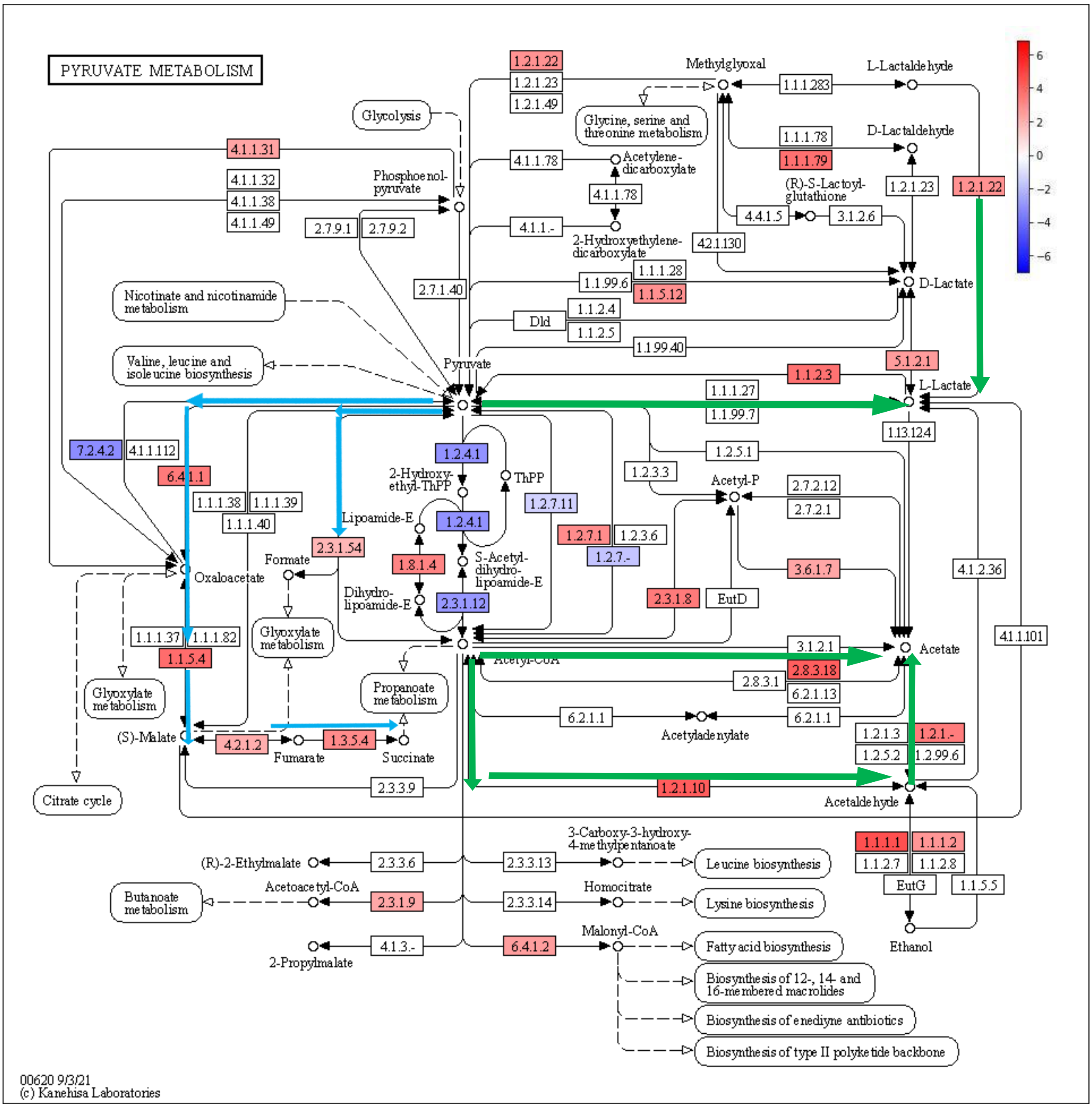

**Supplementary Figure 6.** Colonic SCFAs concentration in anaemic and control animals. Statistical differences are indicated as follows: * p>0.05, ** p>0.01, ***p>0.001, **** p>0.0001

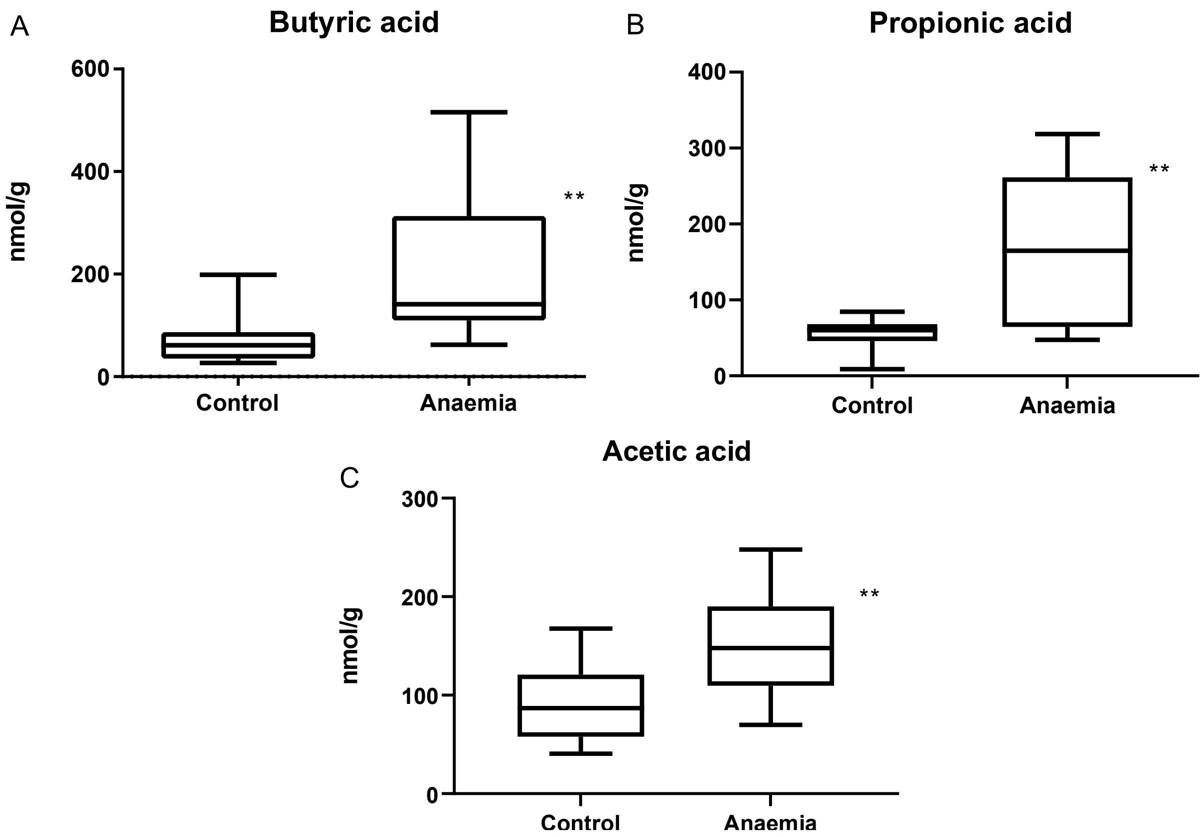

**Supplementary Figure 7**. Purine metabolism pathway (KEGG) coloured according to $log_2FC$ values of statistically significant KO genes. Colour legend: formation of purine-derived nucleotides (green) and nucleotides derivatives (purple)

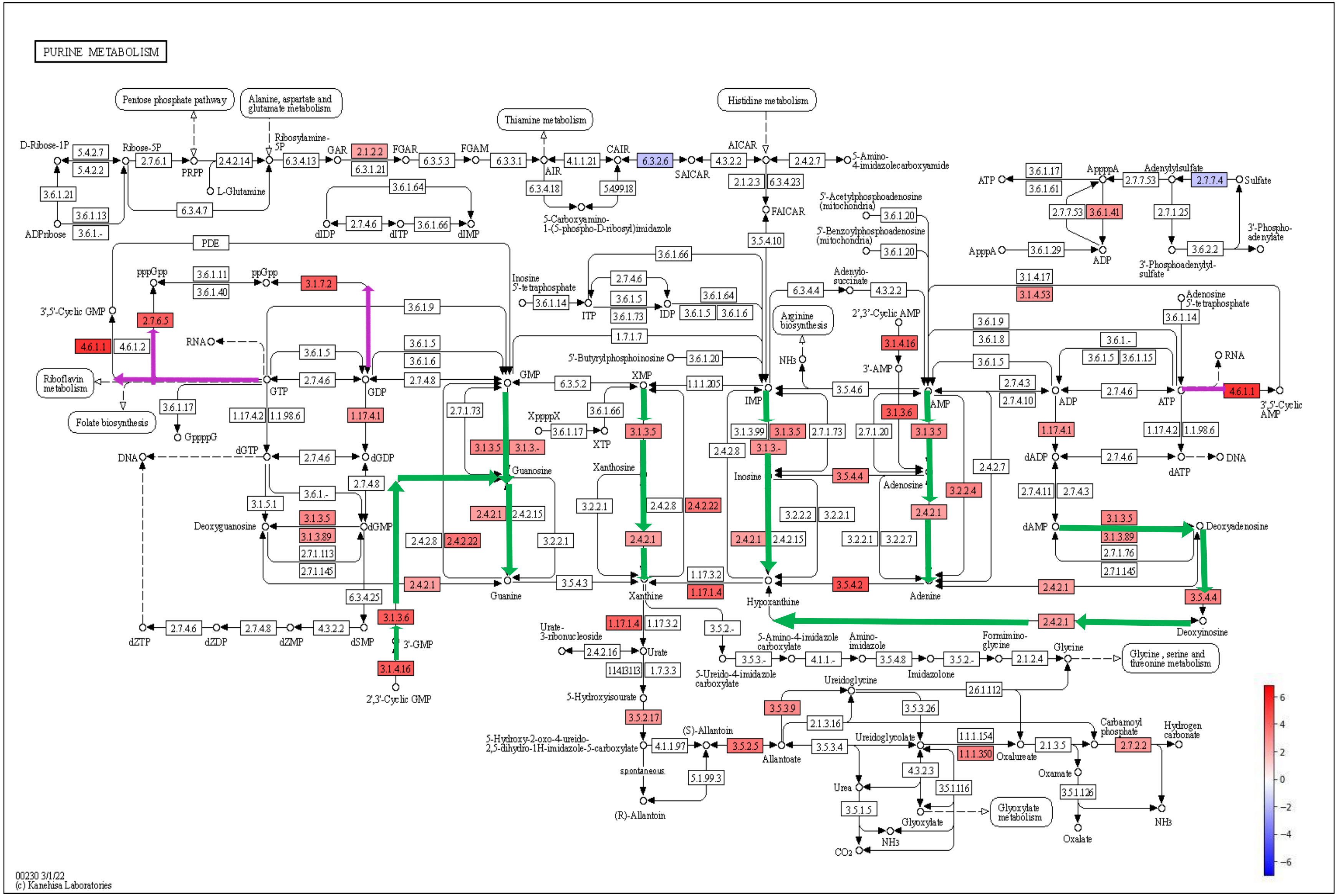

**Supplementary Figure 8**. Pyrimidine metabolism (KEGG) coloured according to $log_2FC$ values of statistically significant KO genes. Colour legend: formation of pyrimidine-derived nucleotides (green)

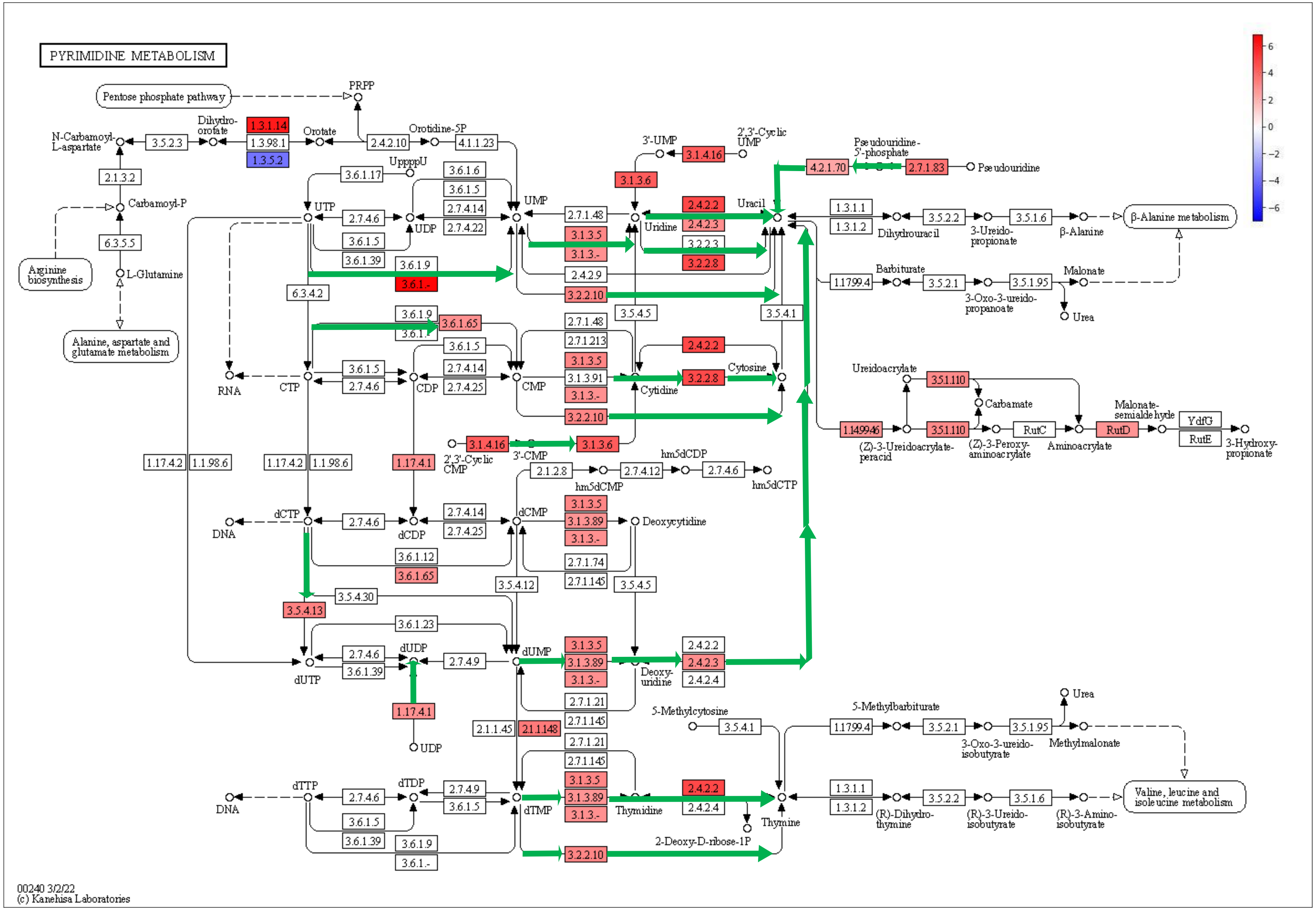

**Supplementary Figure 9**. Bacterial load in colonic contents of control and anaemic animals quantified by qPCR. Mean and standard deviations are shown for biological replicates. Statistical significance is expressed as follows: Statistical significance is expressed as follows: *p < 0.05; **p < 0.01, ***p < 0.001 and ****p<0.0001

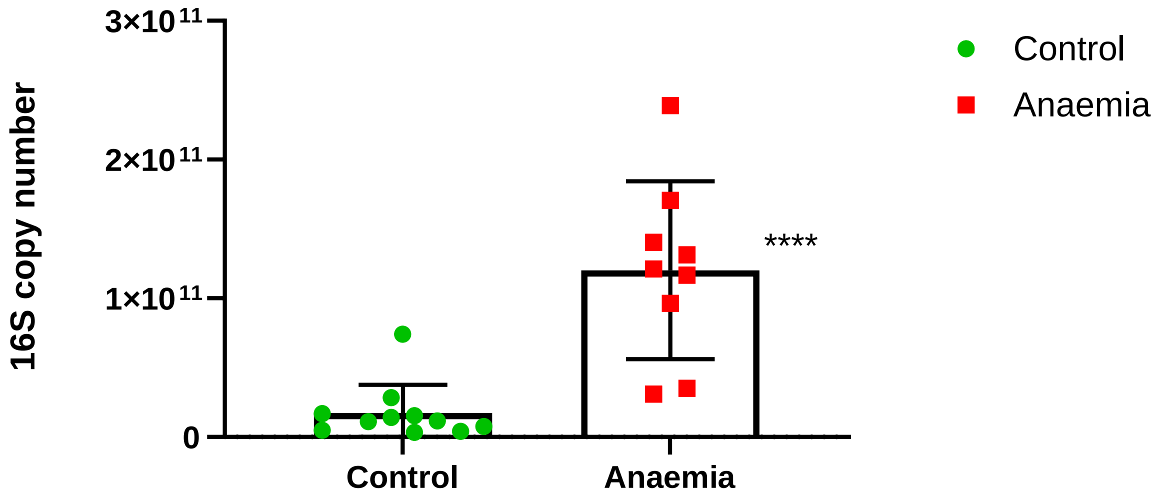

**Supplementary Figure 10.** Histological analyses of the colonic epithelium. (A) Representative colon histological sections of control and anaemic animals. Arrows and asterisks indicate major histopathological differences between groups: leukocyte infiltration (black arrow), epithelial damage and ulceration (red arrow), reduction of goblet cells (asterisk) and reduction of mitosis (green circles). (B) Histological scores of control and anaemic animals (n=5)

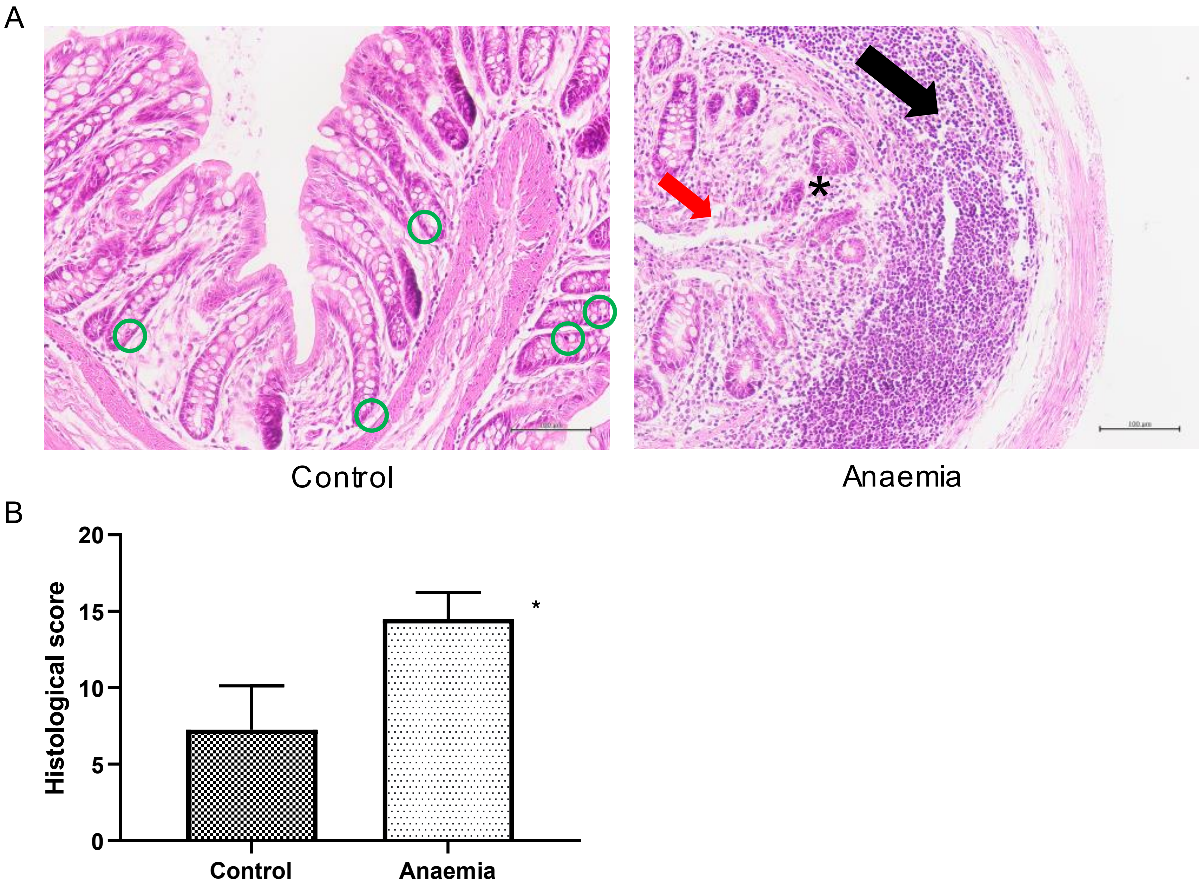

**Supplementary Figure 11**. Study of HIF1α target genes by qPCR. Normalized mRNA expression was calculated using PPIB as housekeeping gene. Mean and standard deviations are shown for biological replicates

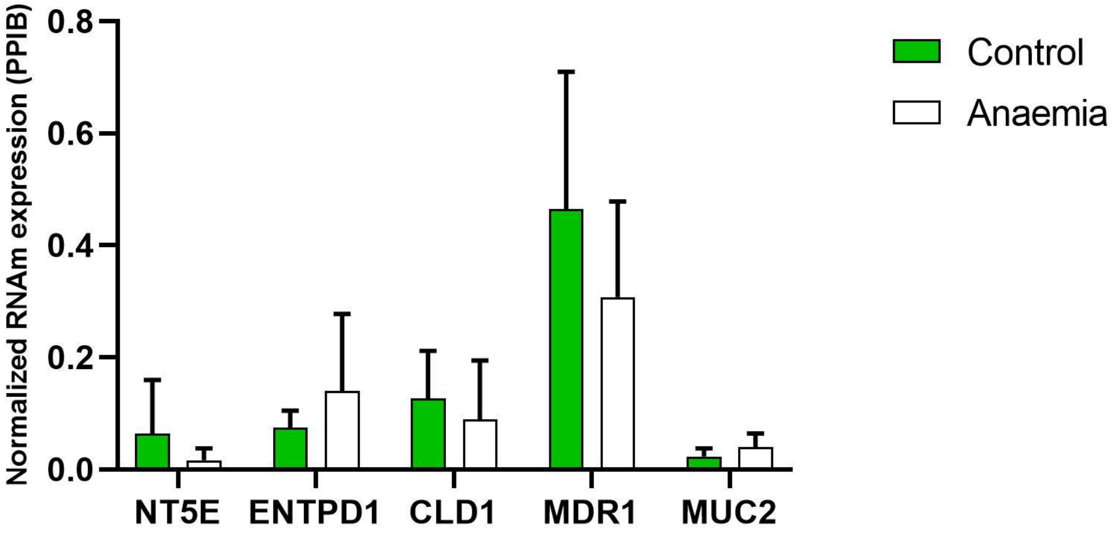
